# Brain maps of general cognitive function and spatial correlations with neurobiological cortical profiles

**DOI:** 10.1101/2024.12.17.628670

**Authors:** Joanna E. Moodie, Colin Buchanan, Anna Furtjes, Eleanor Conole, Aleks Stolicyn, Janie Corley, Karen Ferguson, Maria Valdes Hernandez, Susana Munoz Maniega, Tom C. Russ, Michelle Luciano, Heather Whalley, Mark E. Bastin, Joanna Wardlaw, Ian Deary, Simon Cox

## Abstract

In this paper, we attempt to answer two questions: 1) which regions of the human brain, in terms of morphometry, are most strongly related to individual differences in domain-general cognitive functioning (*g*)? and 2) what are the underlying neurobiological properties of those regions? We meta-analyse vertex-wise *g*-cortical morphometry (volume, surface area, thickness, curvature and sulcal depth) associations using data from 3 cohorts: the UK Biobank (UKB), Generation Scotland (GenScot), and the Lothian Birth Cohort 1936 (LBC1936), with the meta-analytic *N* = 38,379 (age range = 44 to 84 years old). These *g-*morphometry associations vary in magnitude and direction across the cortex (|β| range = -0.12 to 0.17 across morphometry measures) and show good cross-cohort agreement (mean spatial correlation *r =* 0.57, *SD* = 0.18). Then, to address (2), we bring together existing -and derive new -cortical maps of 33 neurobiological characteristics from multiple modalities (including neurotransmitter receptor densities, gene expression, functional connectivity, metabolism, and cytoarchitectural similarity). We discover that these 33 profiles spatially covary along four major dimensions of cortical organisation (accounting for 65.9% of the variance) and denote aspects of neurobiological scaffolding that underpin the spatial patterning of MRI-cognitive associations we observe (significant |*r*| range = 0.21 to 0.56). Alongside the cortical maps from these analyses, which we make openly accessible, we provide a compendium of cortex-wide and within-region spatial correlations among general and specific facets of brain cortical organisation and higher order cognitive functioning, which we hope will serve as a framework for analysing other aspects of behaviour-brain MRI associations.

## 1 Introduction

Individual differences in human cognitive function have well-established though modest associations with individual differences in brain structure. For example, larger total brain volumes are reliably associated with higher general cognitive function (*g*) scores (e.g., ^1^, *N =* 18,363, *r =* 0.275, 95% C.I. = [0.252, 0.299]). The strength of associations between *g* and brain volume varies by brain region (^2,1,3^), and brain-cognition associations also vary by region for other morphometry measures, such as surface area (^4,5^), cortical thickness (^6,5^), curvature (^7^) and sulcal depth (^8, 9^). The parieto-frontal integration theory (P-FIT, ^2^) provides a theoretical basis for the involvement of parieto-frontal brain regions over others in cognition, and there have been expansions and additions to that framework (e.g., ^10^ , ^92, 11^). However, explanations of what regional morphometry-phenotypic association patterns tell us are far from complete. Interpretations are complicated because measures of morphometry from brain MRI are a conflation of multifarious underlying biological properties which also vary by brain region. Thus, in the current paper, we aim to characterise the spatial concordance between two types of brain map, i.e., 1) *g*-morphometry associations and 2) neurobiological profiles. We argue that this could help to decode the neurobiological principles of cortical organisation that facilitate our complex cognitive skills. Formally quantifying that spatial concordance, in turn, might further inform a mechanistic understanding of how cognitive functioning differs between individuals in health and dysfunction.

Until recently, inferences about the underlying biology of brain morphometry-behaviour associations have been predominantly descriptive or indirect, reliant upon findings from unrelated studies to draw together narrative conclusions. This is mainly due to practical limitations in directly relating in vivo MRI findings to information taken postmortem, limitations in the number of biological properties that can be measured in the same individuals, and generally low participant numbers in instances that combine imaging and post-mortem work. However, group-level summary data brain maps for several neurobiological measures are increasingly being made open-source (^12^F,^13^), and can now be straightforwardly registered to the same common brain space as association maps (^12^), allowing for direct quantitative comparisons. Royer et al. (2024) provide a detailed perspective paper discussing the recent rise in the creation and use of cortical profiles to make discoveries about brain organisation (^14^). A landmark study tested spatial associations between neurotransmitter receptor distributions and cortical patterns from case/control analyses of 13 disorders, including depression, obsessive-compulsive disorder, schizophrenia and Parkinson’s disease; and identified spatial co-patterning between neurotransmitter receptors and functional imaging significance patterns derived from Neurosynth for a general factor of cognitive terms (including terms such as attention, stress, and planning) (^15^).

Brain structural differences related to general cognitive functioning have been robustly established and have wide-reaching associations with important life outcomes including everyday function, health, illness, dementia, and death (^16,17,18^). There are increasingly robust analyses which have established cognitive and brain structural associations (e.g., ^3, 19, 20, 21^), yet there remain no large-scale meta-analytic estimates of general cognitive-MRI associations at the level of the cortical vertex. The measure of general cognitive function, as a principal component or latent factor ‘*g*’, offers several relevant properties that make it a behavioural measure of suitable quality for such analyses. It captures the general tendency for cognitive test scores to be positively correlated, and is somewhat invariant to cognitive test content, provided that multiple cognitive domains are captured (^2, 23, 24^). It is one of the most replicated phenomena in psychological science (^25,26^); and its individual differences tend to be quite highly stable across the healthy lifespan (^27^). We bring together *g*-brain structure associations with biological cortical profiles to allow direct (quantitative) inferences about the organising principles of the brain that underlie the cognitive-MRI signals which we observe. Moreover, we produce an extension to prior analytic approaches whereby we go beyond cortex-level spatial correlations (e.g., ^15, 28, 29, 30, 31^), to additionally include regional-level spatial correlations. These regional-level spatial correlations (here, using the Desikan-Killiany atlas, with 34 left/right paired cortical regions) provide nuanced information about 1) the relative strengths of the spatial correlations in different regions and 2) the homogeneity of co-patterning across regions.

In the current paper, we ask two main questions: 1) which regions of the human brain, in terms of their morphometry, are most strongly related to individual differences in domain-general cognitive function? and 2) what are the underlying neurobiological properties of those regions? We address these two important gaps in our knowledge by (see *Figure 1*), first, conducting the largest vertex-wise (298,790 cortical vertices) analysis of *g-*cortex associations across 3 cohorts with 5 morphometry measures (volume, surface area, thickness, curvature, and sulcal depth) in community-dwelling adults (meta-analytic N = 38,379). Then we quantitatively test how those brain regions that are associated with *g* are spatially correlated with the patterning of 33 of the brain’s neurobiological properties across the human cerebral cortex (including neurotransmitter receptor densities, cytoarchitectural, microstructural and functional connectivity similarity gradients, and metabolism) (^32^). We assemble open-source brain maps and derive novel ones; registering them to the same common brain space as our brain morphometry-*g* meta-analytic results; and then we quantitatively test their spatial concordance. Additionally, we identify four principal components that explain the majority of the variance (65.9%) across the 33 maps of the brain’s neurobiological properties, which indicate major dimensions of fundamental brain organisation, and we test their associations with *g-*morphometry cortical profiles. These analyses implement methods for uncovering principles of cortical organisation that are associated with individual differences in our complex cognitive skills.

**Figure 1.**
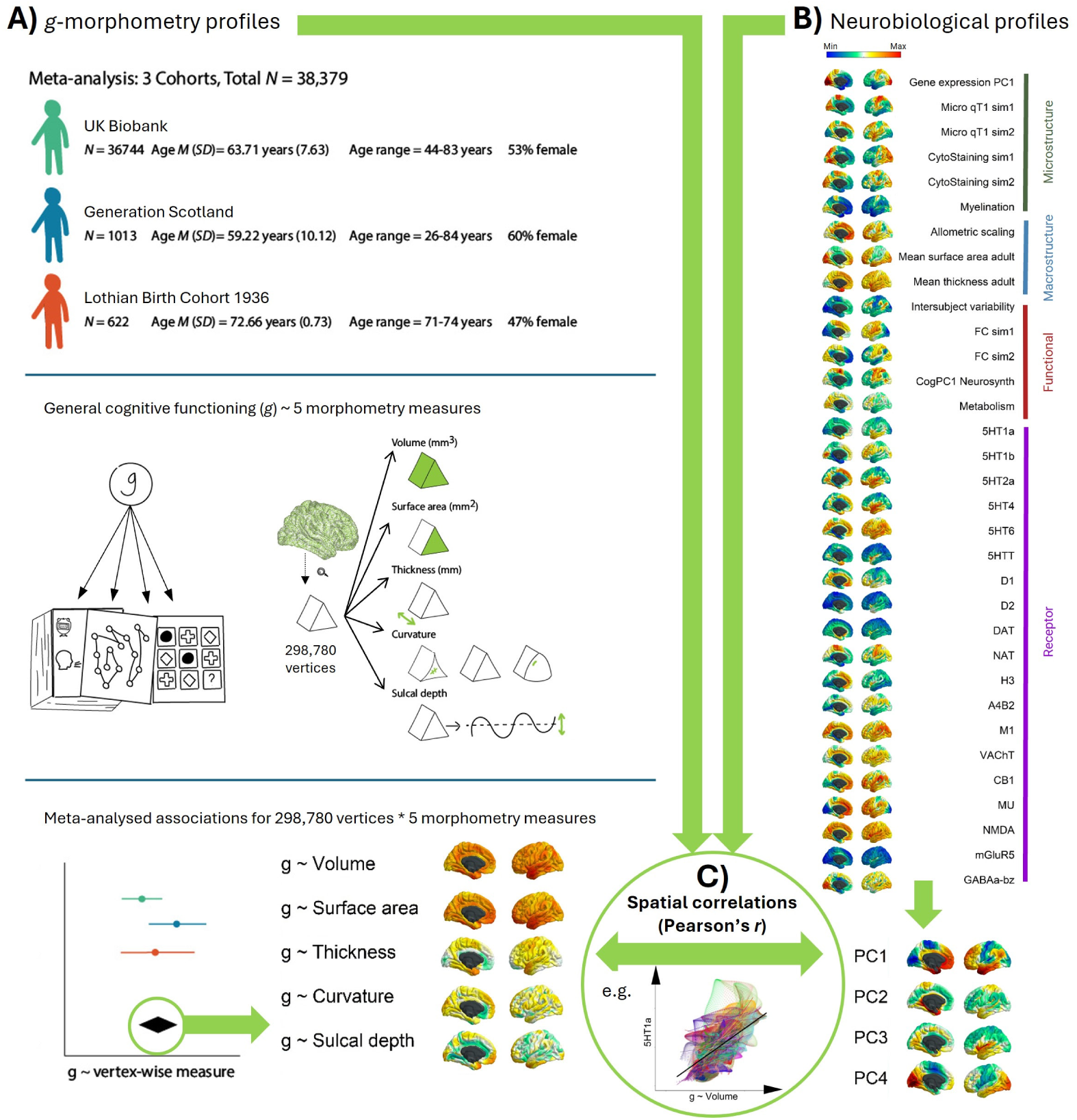
Overview of the methodological approach. Figure 1 note A) Associations between g and 5 measures of brain morphometry (volume, surface area, thickness, curvature and sulcal depth) were estimated for each of three cohorts of community-dwelling adults (UKB, GenScot and LBC1936). These vertex-wise association maps were then meta-analysed, which is the primary outcome of the first step. B) We curated and derived new maps of 33 neurobiological characteristics that vary across the cortex, and registered them to the same anatomical space as the vertex-wise meta-analyses described in A. We also conduct a principal components analysis which identifies four major dimensions of neurobiological organisation across the cortex. C) Finally, we calculate the spatial correlations between g-morphometry profiles and neurobiological profiles, to identify which principles of cortical organisation are most likely candidates for supporting complex cognitive skills.

## 2 Methods

### 2.1 Methods for identifying individual differences

#### 2.1.1 Participants

Data from three cohorts were used to calculate associations between general cognitive functioning (*g*) (and age and sex) and 5 measures of vertex-wise morphometry (volume, surface area, thickness, curvature, and sulcal depth) – the UK Biobank (UKB), Generation Scotland: Scottish Family Health Study (GenScot), and the Lothian Birth Cohort 1936 (LBC1936). They were also used to calculate meta-analysed means for the 5 morphometry measures. These maps will be openly available on publication in fsaverage space at github.com/JoannaMoodie/moodie-brainmaps-cognition.

The UKB (http://www.ukbiobank.ac.uk, ^33^) is a study of ∼500,000 participants, and the data of 40,383 participants who attended the first neuroimaging visit (which included collection of cognitive test data and brain MRI scans) are used in the present analyses. Participants were excluded from the present analysis if their self-reported medical history, taken by a nurse at the data collection appointment, recorded a diagnosis of, for example, dementia, Parkinson’s disease, stroke, other chronic degenerative neurological problems or other demyelinating conditions, including multiple sclerosis and Guillain–Barré syndrome, and brain cancer or injury (a full list of exclusion criteria is provided in *Table S1*). For the global and subcortical brain structures analysis (see *Supplementary Analysis 2*), the sample consisted of *N =* 39,250 (53% female, mean age = 63.91 years, *SD =*7.67 years, and range = 44 to 83 years). For the vertex-wise analyses, participants were included if qcaching in FreeSurfer ran successfully for all 5 morphometry measures. The final *N* for vertex-wise analyses was 36,744 participants (53 % female, mean age = 63.71 years, SD *=* 7.63 years, and range = 44-83 years). The UKB was given ethical approval by the NHS Research Ethics Committee (REC reference 11/NW/0382) and the current analyses were conducted under UKB application number 10279. All participants provided informed consent. More information on the consent procedure can be found at https://biobank.ctsu.ox.ac.uk/crystal/label.cgi?id=100023.

The GenScot imaging sample is a population-based study, developed from the Generation Scotland: Scottish Family Health Study (^34^). Data are available for a maximum of *N =* 1188 participants. Cognitive and MRI data are available for *N =* 1043 participants (60% female, mean age *=* 59.29 years, *SD =* 10.12 years, and range = 26 to 84 years). All 1043 participants were used in the current global and subcortical brain structures analyses (see *Supplementary Analysis 2*). For the vertex-wise analysis, qcaching in FreeSurfer ran successfully for all measures for *N* = 1013 participants (60% female), mean age *=* 59.22 years (*SD =* 10.12 years), age range = 26 to 84 years. GenScot received ethical approval from the NHS Tayside Research Ethics Committee (14/SS/0039), and all participants provided informed consent.

The LBC1936 is a longitudinal study of a sample of community-dwelling older adults who were born in 1936, most of whom took part in the Scottish Mental Survey of 1947 when they were ∼11 years old, and who volunteered to participate in this cohort study at ∼70 years old (^35,36^) https://lothian-birth-cohorts.ed.ac.uk/. The current analysis includes data from the second wave of data collection, which is the first wave at which head MRI scans are available. In total, 731 participants agreed to MRI scanning. After image collection and processing, *N =* 636 participants were included in the specific brain structures analyses conducted in *Supplementary Analysis 2* (47% female, mean age = 72.67 years, *SD =* 0.71 years, and range = 70 to 74 years). Qcaching was unsuccessful for 14 participants, leaving a final *N* for vertex-wise analyses of 622 (47% female, mean age = 72.66 years, *SD =* 0.73 years, and range = 71 to 74 years). The LBC1936 study was given ethical approval by the Multi-Centre Research Ethics Committee for Scotland, (MREC/01/0/56), the Lothian Research Ethics Committee (LREC/2003/2/29) and the Scotland A Research Ethics Committee (07/MRE00/58). All participants gave written informed consent.

#### 2.1.2 Cognitive tests

All three cohorts have data collected across several cognitive tests, covering several cognitive domains (e.g. memory, reasoning and processing speed), which enables the estimation of a latent factor, *g*. The cognitive tests in each cohort have been described in detail elsewhere: UKB (^37^, 10 tests included: Reaction time, Number span, Verbal and numerical reasoning, Trail making B, Matrix pattern, Tower task, Digit-symbol substitution, Pairs matching, Prospective memory, and Paired associates), GenScot (^34^, 5 tests included: Matrix reasoning, Verbal fluency, Mill Hill vocabulary, Digit-symbol substitution, and Logical memory), and LBC1936 (^35,38,39^, 13 tests included: Matrix reasoning, Block design, Spatial span, National Adult Reading Test (NART), Weschler Test of Adult Reading (WTAR), Verbal fluency, Verbal paired associates, Logical memory, Digit span backwards, Symbol search, Digit-symbol substitution, Inspection time, and Four-choice reaction time), see *Supplementary Tables* S2 to S7 for more details. The cognitive tests from each cohort cover various cognitive domains, including Crystallised (verbal) Ability, Reasoning, Processing speed, and aspects of Memory.

A latent factor of *g –* capturing shared variance in performance across all cognitive tests – was estimated for each cohort in a structural equation modelling framework. For UKB and GenScot, no residual covariances between individual cognitive tests were included. For the LBC1936, which has a larger cognitive battery that includes multiple tests for each cognitive domain, *g* has previously been modelled with a hierarchical confirmatory factor analysis approach, to incorporate defined cognitive domains (^38, 39^). Here, in keeping with these previous models, within-domain residual covariances were added for four cognitive domains (visuospatial skills, crystallised ability, verbal memory and processing speed). Latent *g* model fits were assessed using the following fit indices: Comparative Fit Index (CFI), Tucker Lewis Index (TLI), Root Mean Square Error of Approximation (RMSEA), and the Root Mean Square Residual (SRMR). All models had CFI > 0.95, TLI > 0.88, RMSEA < 0.08 and SRMR < 0.04. For specific details of the model fits, see *Table S9*. Results of the *g* measurement models are summarised in *Figure S1* and *Table S8*. For all cohorts, all estimated paths to latent *g* were statistically significant at the *p <* .001 level. To be clear, a *g* factor was found in each of the three cognitive test batteries (that is, a model with a *g* factor had a good fit to the data [i.e., the cognitive tests’ covariance matrices]) and was not imposed upon them.

The latent *g* scores were extracted for all participants (these were calculated with the slightly larger samples that were included in the global and subcortical structures analysis, see *Supplementary Analysis 2*, and these same scores were used for the vertex-wise analysis, which had a slightly smaller sample size due to qcaching failures). All *g* scores were scaled so that higher scores reflected better cognitive performance.

#### 2.1.3 MRI protocols

Detailed information for MRI protocols in all three cohorts are reported elsewhere: UKB (^40^), LBC1936 (^41^) and GenScot (^34^) but are briefly summarised here. In the present sample, UKB participants attended one of four testing sites: Cheadle (∼60%), Reading (∼14%), Newcastle (*∼*25%), and Bristol (∼0.13%). The same type of scanner was used in all four testing sites, a 3T Siemens Skyra, with a 32-channel Siemens head radiofrequency coil. The UK Biobank MRI protocol includes various MRI acquisitions (more details available here https://www.fmrib.ox.ac.uk/ukbiobank/protocol/V4_23092014.pdf) but in this work we exclusively used the T1-weighted MPRAGE volumes. For T1-weighted images, 208 sagittal slices were acquired with a field view of 256 mm and a matrix size of 256 x 256 pixels, giving a resolution of 1 x 1 x 1 mm^3^. The repetition time was 2000 ms and the echo time was 2.01 ms (^40^).

GenScot had 2 testing sites: Aberdeen (in the present sample, *N =* 528, 51% of the total sample) and Dundee (*N =* 515, 49% of the total sample). Detailed information about the GenScot structural image acquisitions is available here https://wellcomeopenresearch.org/articles/4-185. For the current analysis, we used the T1-weighted fast gradient echo with magnetisation preparation volume. The Aberdeen site used a 3T Philips Achieva TX-series MRI system (Philips Healthcare, Best, Netherlands) with a 32-channel phased-array head coil and a back facing mirror (software version 5.1.7; gradients with maximum amplitude 80 mT/m and maximum slew rate 100 T/m/s). For T1-weighted images, 160 sagittal slices were acquired with a field of view of 240 mm and a matrix size of 240 x 240 pixels, giving a resolution of 1 x 1 x 1 mm^3^. Repetition time was 1968 ms, echo time was 3.8 ms and inversion time was 1031 ms. In Dundee, the scanner was a Siemens 3T Prisma-FIT (Siemens, Erlangen, Germany) with 20 channel head and neck phased array coil and a back facing mirror (Syngo E11, gradient with max amplitude 80 mT/m and maximum slew rate 200 T/m/s). For T1-weighted images 208 sagittal slices were acquired with a field of view of 256 mm and matrix size 256 x 256 pixels giving a resolution of 1 x 1 x 1 mm^3^. Repetition time was 1740 ms, echo time was 2.62 ms, and inversion time was 900 ms ^34^.

All LBC1936 participants were scanned in the same scanner in the same clinic, using a GE Signa LX 1.5T Horizon HDx clinical scanner (General Electric, Milwaukee, WI) with a manufacturer supplied 8-channel phased array head coil. More information on the structural image acquisitions for the LBC1936 cohort is available in (^41^). For T1-weighted images (3D IR-Prep FSPGR), 160 coronal slices were acquired, with a field of view of 256 mm and a matrix size of 192 x 192 pixels (zero filled to 256 x 256) giving a resolution of 1 x 1 x 1.3 mm^3^. The repetition time was 10 ms, echo time was 4 ms and inversion time was 500 ms.

For all cohorts, the FreeSurfer image analysis suite (http://surfer.nmr.mgh.harvard.edu/) was used for cortical reconstruction and volumetric segmentation. The 46 global and subcortical structures (including grey matter, white matter and ventricles), used in *Supplementary Analysis 2*, were available for each cohort in the aseg FreeSurfer outputs. Vertex-wise surface values for 5 morphometry measures (volume, surface area, thickness, curvature and sulcal depth) were available at 9 smoothing tolerances (0, 5, 10, 15, 20, 25, 30, 35 and 40 mm FWHM, full width half maximum Gaussian kernel) by running the -qcache flag.

Each cohort used a different version of FreeSurfer: UKB = v6.0, GenScot = v5.3, LBC1936 = v5.1. The LBC1936 and GenScot parcellations have previously undergone quality control, with manual editing to rectify parcellation issues including skull stripping, tissue identification and regional boundary lines. The UKB regional data were extracted from the bulk-downloaded aseg files provided by the UK Biobank. For the current study, UKB values more than 4 standard deviations from the mean for any global or subcortical brain structure volume were excluded (corresponding to < 1.2% of the data per variable; *M =* 87.97, *SD =* 121.75, range = 0 to 474 participants) – note, outlier values were excluded by region, rather than participant-wise exclusions

#### 2.1.4 Morphometry measures

The morphometry measurements are illustrated in *Figure 1A* (middle panel). Volume is the amount of three-dimensional space of a vertex, surface area is the total area of the cortical sheet section of the vertex, and thickness is the distance between the pial and white matter cortical surfaces. If thickness were uniform across the vertex, volume would be the product of surface area and thickness, but this relationship is more complex in practice. For curvature, a value of zero represents no curvature “– “; those with negative values are curving up (convex) “◠“; those with positive values are curving down (concave) “◡“. The sulcal depth is a measure of how removed a vertex is from a theoretical mid-surface that is estimated between the gyri and sulci (vertices on the mid surface receive a value of 0). A more positive sulcal depth suggests a deeper location (i.e., away from the scalp) and a more negative value is shallower (i.e., towards the scalp). Deep sulci tend to have more concave curvature, shallower regions tend to have curvature magnitudes nearer to zero, and gyri (defined here as regions with negative values for “sulcal depth”) tend to correspond to convex curvature. The measure of curvature provides information about how the cortex folds at the local level, while sulcal depth provides a more macroscopic perspective on the depth of sulci relative to the midpoint of the cortical surface, offering insights into the overall brain folding complexity.

#### 2.1.5 Meta-analyses

We chose a 20 mm FWHM smoothing tolerance for our main cohort meta-analyses , in line with our previous work (^42, 43^). For each cohort, a standardised β was estimated between *g* and each vertex for the 5 vertex-wise morphometry measures. Each participant’s cortical surface was aligned to the fsaverage template. Out of 327,684 initial vertices along the fsaverage surface, there are 298,790 vertices labelled as “cortex”, and these vertices are analysed here.

We first checked the cross-cohort agreement of the means of the five measures of cortical morphometry across the three cohorts. Spatial variations in measures of cortical volume, surface area, thickness, curvature and sulcal depth were highly stable between cohorts -all r > 0.843 (see *Table S*10, for the global means see *Figure S*3, and for the meta-analysed mean cortical maps see *Figure* S2). From these analyses, the meta-analysed mean profiles for surface area and thickness were included in the spatial correlation analyses in section 3.4.

For cohort-based association analyses, all brain measures were controlled for head position in the scanner (X, Y and Z coordinates, from UKB codes 25756, 25757, and 25758; and estimated in FreeSurfer for GenScot and LBC1936), testing site (for UKB and GenScot only) and, for LBC1936 only, time lag (because it was the only cohort with a time lag between cognitive and MRI appointments, *M* lag *=*65.08 days, *SD =* 37.77). Age and sex were included as covariates in models when they were not the variable of interest.

To characterise which regions of the human brain, in terms of morphometry, are most strongly related to individual differences in *g*, we then meta-analysed the vertex-wise *g* associations between the three cohorts with random effects models. This type of model was deemed the most appropriate due to the differing characteristics (e.g., age range) between the cohorts. Vertex-wise brain maps for age and sex associations were meta-analysed in the same way. For vertex-wise analyses of age, only GenScot and UKB cohorts were included due to the narrow age range of the LBC1936 cohort (mean age = 72.67 years, *SD =* 0.71). For all meta-analyses, between-cohort age moderation analyses were additionally conducted (i.e., mean age for each cohort was included as a moderator in the *rma* function in the *metafor* package ^90^). UKB and GenScot have larger age ranges, and lower mean age (*M* = 63.91 years, range = 44-83; *M* = 59.29 years, range = 26-84 respectively) compared to the LBC1936 (*M* = 72.67 years, range = 70-74). Therefore, although we included age as a covariate within cohorts, it remains possible that between-cohort age differences affect the brain associations. Any between-cohort age moderation analyses significant at the α < .05 level are reported below.

### 2.2 Neurobiological cortical profiles

The 33 included neurobiological cortical profiles were derived from several modalities, including: in vivo MRI, rsfMRI (resting state functional MRI), fcMRI (resting-state functional connectivity MRI), PET (positron emission tomography) scans and postmortem tissue. Several of these cortical profiles are openly available online through neuromaps (^12^), and BigBrainWarp (^13^), and we registered all profiles to fsaverage space using neuromaps. We include maps of metabolism (we calculated a principal component, derived from previously published, open source maps of cerebral blood flow, oxygen metabolism, and glucose metabolism, ^79^); similarity gradients of cytoarchitecture (staining intensity), functional connectivity, and microstructure (^13^); the first principal component of gene expression from the abagen toolbox (^44^); cortical myelination T1/T2 ratio (^45^) and 19 neurotransmitter receptor densities (^12^). These maps are described in more detail in Table 1 and in the subheadings below.

**Table 1.** Description of the neurobiological cortical profiles. Descriptive statistics of all vertex-wise cortical profiles in this paper are available in Table S13.

| Map | Source | Original source | Data source | Participants | Category | Type | Higher value = |
| --- | --- | --- | --- | --- | --- | --- | --- |
| Gene expression PC1 | Neuromaps | abagen toolbox <sup>44</sup> | Postmortem (gene expression) | *N* = 6 adult human donors, 1 female, ages 24 to 57 years | Microstructure | Principal component 1 of gene expression | A higher positive component score for PC1 |
| Microstructural similarity | BigBrainWarp | Paquola et al. <sup>13,77</sup> | In Vivo (qT1) | *N* = 50 healthy adults <sup>46</sup>, 23 women, age mean = 29.54 (SD = 5.62) | Microstructure | Eigenvectors 1 and 2 | A higher positive eigenvector score |
| Cytoarchitectural similarity | BigBrainWarp | Paquola et al. <sup>13</sup> | Postmortem (BigBrain, histology) | *N* = 1 donor, 65-year-old male | Microstructure | Eigenvectors 1 and 2 from BigBrain staining intensity profiles | A higher positive eigenvector score |
| Myelination (T1/T2 contrast) | Neuromaps | Glasser <sup>45</sup> | In vivo (T1/T2) | *N* = 449 young adults (ages 22–35) from the Human Connectome Project (HCP) | Microstructure | T1w to T2w ratio | Higher myelin density |
| Allometric scaling | Calculated for the current paper | Current paper | In vivo (T1, MRI) | *N* = 38,379 adults, from UKB, GenScot and LBC1936; age range: 26–84 | Macrostructure | Log-log regression coefficient for vertex surface area predicted by total surface area | Higher coefficient (greater deviation from isometry) |
| Mean surface area adult | Calculated for the current paper | Current paper | In vivo (T1, MRI) | *N* = 38,379 adults, from UKB, GenScot and LBC1936; age range: 26–84 | Macrostructure | Mean | Larger surface area |
| Mean thickness adult | Calculated for the current paper | Current paper | In vivo (T1, MRI) | *N* = 38,379 adults, from UKB, GenScot and LBC1936; age range: 26–84 | Macrostructure | Mean | Thicker |
| Intersubject variability | Neuromaps | Mueller et al. <sup>47</sup> | In vivo (fcfMRI) | *N* = healthy subjects (age 51.8±6.99, 9 female) | Functional | Variability in fcfMRI data | More variability in rsfMRI |
| Functional connectivity similarity | BigBrainWarp | Paquola et al. <sup>13</sup> | In vivo (rsfMRI) | 50 healthy adults <sup>46</sup>, 23 women, age mean = 29.54 (SD = 5.62) | Functional | Eigenvectors 1 and 2 from rsfMRI data | A higher positive eigenvector score |
| Cognition PC1 Neurosynth | Neuromaps | Yarkoni et al. <sup>48</sup> | In vivo (fMRI) | Unknown | Functional | Principal component 1 of fMRI patterns associated with various terms in Neurosynth | A higher positive component score for PC1 |
| Metabolism | Neuromaps, and PC1 calculated for the current paper | Vaishnavi et al.<sup>79</sup> | In vivo (PET) | *N* = 33 neurologically normal young adults, 19 women, 14 men, 20-33 years old | Functional | Principal component 1 of cerebral blood flow, oxygen metabolism and glucose metabolism maps | Higher positive values on PC1 (higher values on the maps for CBF, CMRO2 and CMRGlu) |
| 5HT1a (Serotonin) | Neuromaps, (Hansen et al.) | Savli et al.<sup>49</sup> | In vivo (PET) | *N* = 35, age mean = 26.3 (SD = 5.2) | Receptor density | Group averaged PET data | Higher density of each receptor |
| 5HT1b (Serotonin) | Neuromaps, (Hansen et al.) | Gallezot et al.<sup>50</sup> | In vivo (PET) | *N* = 65, age mean = 33.7 (SD = 9.7) | Receptor density | Group averaged PET data | Higher density of each receptor |
| 5HT2A (Serotonin) | Neuromaps, (Hansen et al.) | Beliveau et al.<sup>51</sup> | In vivo (PET) | *N* = 29, age mean = 22.6 (SD = 2.7) | Receptor density | Group averaged PET data | Higher density of each receptor |
| 5HT4 (Serotonin) | Neuromaps, (Hansen et al.) | Beliveau et al.<sup>51</sup> | In vivo (PET) | *N* = 59, age mean = 25.9 (SD = 5.3) | Receptor density | Group averaged PET data | Higher density of each receptor |
| 5HT6 (Serotonin) | Neuromaps, (Hansen et al.) | Radhakrishnan et al.<sup>52</sup> | In vivo (PET) | *N* = 30, age mean = 36.6, SD = 9) | Receptor density | Group averaged PET data | Higher density of each receptor |
| 5HTT (Serotonin) | Neuromaps, (Hansen et al.) | Beliveau et al.<sup>51</sup> | In vivo (PET) | *N* = 100, age mean = 25.1 (SD = 5.8) | Receptor density | Group averaged PET data | Higher density of each receptor |
| D1 (Dopamine) | Neuromaps, (Hansen et al.) | Kaller et al.<sup>53</sup> | In vivo (PET) | *N* = 13, age mean = 33 years (SD = 13) | Receptor density | Group averaged PET data | Higher density of each receptor |
| D2 (Dopamine) | Neuromaps, (Hansen et al.) | Smith et al. and Sandiego et al.<sup>54, 55</sup> | In vivo (PET) | *N* = 37, age mean = 48.4 years (SD = 16.9);<br>*N* = 55, age mean = 32.5 years (SD = 9.7) | Receptor density | Group averaged PET data | Higher density of each receptor |
| DAT (Dopamine) | Neuromaps, (Hansen et al.) | Dukart et al.<sup>56</sup> | In vivo (PET) | *N* = 174, age mean = 61 years (SD = 11) | Receptor density | Group averaged PET data | Higher density of each receptor |
| NAT (Norepinephrine) | Neuromaps, (Hansen et al.) | Ding et al.<sup>57</sup> | In vivo (PET) | *N* = 77, age mean = 33.4 (SD = 9.2) | Receptor density | Group averaged PET data | Higher density of each receptor |
| H3 (Histamine) | Neuromaps, (Hansen et al.) | Gallezot et al.<sup>58</sup> | In vivo (PET) | *N* = 8, age mean = 31.7 (SD = 9.0) | Receptor density | Group averaged PET data | Higher density of each receptor |
| A4B2 (Acetylcholine) | Neuromaps, (Hansen et al.) | Hillmer et al.<sup>59</sup> | In vivo (PET) | *N* = 30, age mean = 33.5 years (SD = 10.7) | Receptor density | Group averaged PET data | Higher density of each receptor |
| M1<br>(Acetylcholine) | Neuromaps,<br>(Hansen et al.) | Naganawa et<br>al. <sup>60</sup> | In vivo (PET) | $N = 24$, age<br>mean = 40.5<br>(SD = 11.7) | Receptor density | Group<br>averaged PET<br>data | Higher<br>density of<br>each<br>receptor |
| VACht<br>(Acetylcholine) | Neuromaps,<br>(Hansen et al.) | PIs: Tuominen,<br>L. & Guimond,<br>S.;<br>Aghourian et<br>al.<sup>61</sup>;<br>Bedard et al. <sup>62</sup> | In vivo (PET) | $N = 4$, age mean<br>= 37 (DF =<br>10.2);<br>$N = 18$ (age<br>mean = 66.8, SD<br>= 6.8);<br>$N = 5$, age mean<br>= 68.3 (SD =<br>3.1) | Receptor density | Group<br>averaged PET<br>data | Higher<br>density of<br>each<br>receptor |
| CB1<br>(Cannabinoid) | Neuromaps,<br>(Hansen et al.) | Normandin et<br>al. <sup>63</sup> | In vivo (PET) | $N = 77$, age<br>mean = 30<br>years (SD = 8.9) | Receptor density | Group<br>averaged PET<br>data | Higher<br>density of<br>each<br>receptor |
| MU<br>(Opioid) | Neuromaps,<br>(Hansen et al.) | Kantonen et al<br>, <sup>64</sup> | In vivo (PET) | $N = 204$, age<br>mean = 32.3<br>years (SD =<br>10.8) | Receptor density | Group<br>averaged PET<br>data | Higher<br>density of<br>each<br>receptor |
| NMDA<br>(Glutamate) | Neuromaps,<br>(Hansen et al.) | Galovic et al.<sup>65</sup>,<br><sup>66</sup> | In vivo (PET) | $N = 29$, age<br>mean = 40.9<br>years (SD =<br>12.7) | Receptor density | Group<br>averaged PET<br>data | Higher<br>density of<br>each<br>receptor |
| mGluR5<br>(Glutamate) | Neuromaps,<br>(Hansen et al.) | Smart et al. <sup>67</sup>;<br>PIs: Rosa-Neto,<br>P. &<br>Kobayashi, E.;<br>DuBois et al. <sup>68</sup> | In vivo (PET) | $N = 73$, age<br>mean = 19.9<br>(SD = 3.04);<br>$N = 22$, age<br>mean = 67.9<br>(SD = 9.6);<br>$N = 28$, age<br>mean = 33.1<br>(SD = 11.2) | Receptor density | Group<br>averaged PET<br>data | Higher<br>density of<br>each<br>receptor |
| GABAA-bz<br>(GABA) | Neuromaps,<br>(Hansen et al.) | Nørgaard et<br>al.<sup>69</sup> | In vivo (PET) | $N = 16$, age<br>mean = 32.3<br>(SD = 10.8) | Receptor density | Group<br>averaged PET<br>data | Higher<br>density of<br>each<br>receptor |

#### 2.2.1 Gene expression

The gene expression map was the first principal component of gene expression from the abagen toolbox (^44^). It is available in neuromaps in fsaverage 10k space, and we resampled it to fsaverage 164k using the transforms function in neuromaps. Seemingly due to registration error, there were more vertices outside the cortical mask than for the association maps, and most of the other neurobiological profiles. There were 292076 vertices included in the cortical mask.

#### 2.2.2 T1/T2 ratio derived myelination

The map of cortical myelin content was previously derived from T1w to T2w ratios (maps calculated in ^45^, method described in detail in ^70^). T1/T2 ratio is thought by some to be a good estimate of relative myelin content across the cortical surface ^70^, although it is important to note that this method only provides a proxy for myelin content, and also reflects tissue microstructures other than myelin, such as axon density and dendrite density and iron content ^71^. Indeed, in some contexts, T1/T2 ratios do not make a good proxy for myelination ^72^. Nevertheless, we call this map “myelination”, in line with the source paper.

#### 2.2.3 Allometric scaling

To obtain a map of allometric scaling, we calculated associations between total surface area and vertex-wise surface for UKB, GenScot and LBC1936 cohorts and then meta-analysed them. Allometric scaling was calculated based on previous work (^73^), with a log-log regression coefficient for vertex surface area predicted by total surface area. Allometric scaling shows which vertices have a disproportionately larger surface area in people with bigger brains. Comparable maps of allometric scaling are available for younger cohorts compared to the current sample (the Philadelphia Neurodevelopmental Cohort, PNC, and a National Institutes of Health, NIH, sample). These maps were created with samples 8 to 23 years old (*N =* 1373) and 5 to 25 years old (*N* = 792). These previously calculated maps were correlated at *r =* 0.679, with each other, and at *r =* 0.430 and *r =* 0.378, respectively with our log vertex area ∼ log total surface area maps (which are created with data on adults, age range = 26-83 years old). As most maps of interest included in the current study are derived from adult data, we use the allometry map that we created from our current samples in our further analyses. In our calculations of allometric scaling, the standardised estimates were strongly spatially correlated between cohorts (LBC1936-GenScot *r =* 0.764, GenScot-UKB *r =* 0.773 and UKB-LBC1936, *r =* 0.730, all *p* < 2.2x10^-16^), showing that across cohorts, the regions that tended to be larger with increasing brain size were consistent.

#### 2.2.4 Mean surface area and mean thickness

Meta-analysed mean values for surface area and thickness were calculated using UKB, GenScot and LBC1936 data (meta-analytic N = 38,379). These are mapped to the cortex in *Figure S*2 (between cohort spatial correlations were all r > 0.843, see *Table S*10).

#### 2.2.5 Intersubject variability

Intersubject variability in rsfMRI varies spatially across the cortex (^47^)F70. In other words, for some regions, rsfMRI is similar across participants, whilst in other regions, it is more variable. An openly available cortical map of intersubject variability was available at 1k density in fsaverage space in neuromaps (^12^), and we registered it to 164k density in fsaverage space.

#### 2.2.6 Cognition PC1 from Neurosynth

Component 1 from a principal components analysis of cognitive terms in Neurosynth (which is a database of task-based fMRI results) (^48^). It is available as a cortical map in MNI152 2mm space, and we registered it to fsaverage_164k space (^12^).

#### 2.2.7 Similarity gradients: cytoarchitecture, functional connectivity and microstructure

Cortical similarity gradients of cytoarchitecture, functional connectivity and microstructure are readily available in fsaverage_164k space in BigBrainWarp https://bigbrainwarp.readthedocs.io/en/latest/. The BigBrain data (*N* = 1) (^74^) was used to create the cytoarchitectural similarity map, and the microstructural and functional maps were based on *N* = 50 healthy adults, for whom multiscale MRI data is openly available^75^. The cytoarchitectural gradients data are based on staining intensity profiles. The microstructural gradients are based on qT1 intensity, a quantitative measure of longitudinal relaxation time, which provides an in vivo proxy for cortical microstructure. The functional connectivity gradients are based on rsfMRI-derived functional connectomes. The methods used to obtain these maps are available in detail in the documentation for BigBrainWarp https://bigbrainwarp.readthedocs.io/en/latest/ and micapipe https://micapipe.readthedocs.io/en/latest/. Briefly, the cytoarchitectural and functional gradients were calculated with diffusion map embedding, which is a nonlinear manifold learning technique (^76^), applied to cross-correlations of vertex-wise staining intensity profiles (^77^), and the microstructural and functional connectivity axes are calculated using a microstructural profile covariance (MPC) approach (^78^), which provides eigenvectors of common variation. The percentage of variance explained by the first two eigenvectors for each measure were: cytoarchitectural similarity 1 = 42% and 2 = 35%, for functional connectivity 1 = 12.9% and 2 = 6.5%, and for microstructural similarity 1 = 59.0% and 2 = 10.5%. In an attempt to make the cortical similarity gradients from BigBrainWarp of comparable granularity to our individual difference association maps (20 mm FWHM smoothing), we performed additional smoothing on the BigBrainWarp-sourced maps. The cytoarchitectural gradients were previously smoothed by 2 mm FWHM Gaussian kernel (^13^), and so we smoothed these with an additional 18 mm FWHM kernel. With the approximate guideline that the rsfMRI data approximately has a smoothing kernel of 6 mm (^13^), we smoothed the functional connectivity gradients with an additional 14 mm kernel. To our knowledge, the microstructural similarity gradients available in BigBrainWarp have not previously been smoothed, and here we smoothed them with a 20 mm FWHM kernel.

#### 2.2.8 Metabolism

Metabolism data, available in neuromaps (^12^), was originally collected in 2010 by Vaishnavi et al. (^79^). These data are an average of the PET maps across 33 young adults at rest. Here, we looked at 3 measures of cortical metabolism, cerebral blood flow, oxygen metabolism and glucose metabolism. We registered them from fsLR_164k to fsaverage_164k in neuromaps (^12^). These maps were all highly spatially correlated with each other (all *r* > 0.8), and the first principal component explained 88% of the variance, with all three loadings > 0.5, see *Figure S*17. It is this first principal component of cortical metabolism that we used for the “metabolism” map, included in our spatial correlation analyses. More positive values denote higher metabolic activity.

#### 2.2.9 Neurotransmitter receptor densities

Hansen et al. recently collected receptor density maps for serotonin (5HT1A, 5HT1B, 5HT2A, 5HT4, 5HT6, and 5HTT), dopamine (D1, D2, DAT), norepinephrine (NAT), histamine (H3), acetylcholine (α4β2, M1, VAChT), cannabinoid (CB1), opioid (Mu), glutamate (NMDA, mGluR5) and GABA (GABA A/BZ) neurotransmitter receptors (^15^). They are available through neuromaps and we registered them from MNI152 space to fsaverage_164k space, also using neuromaps (^12^). Due to the lower spatial resolution of PET data, no further smoothing was performed.

### 2.3 Spatial correlations

The *g-*morphometry maps described above (*Figure 1A*) were then spatially correlated with 1) 33 neurobiological maps, and 2) 4 PCs derived from the 33 neurobiological maps and denoting core components of cortical neurobiological organisation (*Figure 1B*). Spatial correlations were calculated using Pearson’s *r* (e.g. for each *g-*morphometry map, the vector of cortical vertices was correlated with each other map’s vector of cortical vertices). Alexander-Bloch’s spin-based permutation test was used to calculate *p*-values^80^. Each *g-* morphometry map was spun randomly 10000 times, and from the resulting null distributions of the correlations, *p* values were calculated. The Pearson’s *r* and *p_spin* values for spatial correlations between all maps included in the main correlation analyses in the current paper are available in the Supplementary Tabular Data file.

All 33 neurobiological maps were inputted into a PCA which was calculated in R using the *prcomp* function, with the aim to identify core components of neurobiological spatial cortical organisation. With all vertices in the cortical mask, across the 33 maps, the vertex count was *N* = 292,056 for the PCA. Four components were extracted, based on the variance they explained (together, 65.9%), and rotated with the varimax method.

We also calculated within-region vertex-wise spatial correlations for *g*-morphometry and neurobiological map correlations. To do this, we used the fsaverage annotation files to identify which vertices were included in each region according to the Desikan-Killiany atlas (34 left/right paired cortical regions), and then calculated the spatial correlations separately for each region. These within-region analyses offer important nuance to the cortex-wide spatial correlation statistics in the form of two additional features: 1) the relative strength of spatial correlations within different regions, and 2) the homogeneity of correlations among regions.

### 2.4 Supplementary analyses

We conducted three supplementary analyses. *Supplementary Analysis 1* addresses the current lack of consensus about optimal smoothing parameters. Noise in the data, due to registration inaccuracies, is minimized when the cortex is parcellated into larger regions (i.e., greater smoothing) but, when the cortex is parcellated into smaller regions (i.e., less smoothing), the % variance explained increases (^81^) due to the additional information provided. Thus, at the vertex-wise level, there is a balance to be struck between the benefits of reducing noise in the data, and the problem that increasing to higher levels of smoothing will, at a point, remove fine-grained spatial information and thus reduce the spatial specificity of detected associations. Lerch and Evans (2005) analysed the effect of different smoothing tolerances on cortical thickness measurement sensitivity, and they concluded an optimal kernel size of 30 mm FWHM (^82^, *N =* 25). Some studies use 30 mm (^83,84^), and other common choices are 5 mm (^4^), 10 mm (^85,86^), 15 mm (^87,88^) or 20 mm (^42,89^). We investigated the effects of smoothing tolerances on *g*-morphometry associations here (see *Supplementary Analysis 1*), and the results suggest that generally, across morphometry measures, 10-20 mm FWHM tends to maximise noise reduction while maintaining localised effects. These results may aid future smoothing tolerance choices for similar analyses.

*Supplementary Analysis 2* focuses on global and sub-cortical associations with *g*, age, and sex. Although much previous work on *g-*brain associations focuses on the cortex, sub-cortical structures are becoming increasingly recognized for their associations with cognitive function.

*Supplementary Analysis 3* tests whether *g-*morphometry associations differ by sex when they are calculated separately for each sex. The spatial correlations were all *r* > 0.753 for UKB, suggesting that meaningful sex differences do not exist in the general population.

### 2.5 Analysis software

Within-cohort vertex-wise analyses were conducted in surfstat http://www.math.mcgill.ca/keith/surfstat/ in MATLAB. All meta-analyses (metafor, ^90^), structural equation models (lavaan, ^91^), and spatial correlations were conducted in R 4.0.2.

(R Core Team, 2020). Structural equation models were estimated with the full information maximum likelihood method.

### 2.6 Data Availability

All UKB data analysed here were provided under project reference 10279. A guide to access UKB data is available from http://www.ukbiobank.ac.uk/register-apply/. To access data from the GenScot study, see https://www.research.ed.ac.uk/en/datasets/stratifying-resilience-and-depression-longitudinally-GenScot-a-dep, and to access the Lothian Birth Cohort data, see https://www.ed.ac.uk/lothian-birth-cohorts/data-access-collaboration. The BigBrainWarp toolbox, released by Paquola et al. (^13^), is available for download at https://bigbrainwarp.readthedocs.io/en/latest/. The neuromaps toolbox is available at https://github.com/netneurolab/neuromaps. Analysis script templates and the vertex-wise β estimate cortical maps for *g*, age, sex and allometric scaling, along with the meta-analytic means, the principal component of the metabolism maps, and the four principal components derived from the 33 neurobiological maps that are calculated in the current paper will be available on publication here: github.com/JoannaMoodie/moodie-brainmaps-cognition.

## 3 Results

### 3.1 Associations between general cognitive functioning and brain morphometry: cross-cohort replicability and meta-analysis results

#### 3.1.1 Global morphometry associations with *g*

At the global cortical level (measures summed across all vertices, associations calculated for each cohort, then meta-analysed), participants with higher general cognitive function had a greater total cortical volume (β = 0.178, SE = 0.035, *p* = 3.18x10^-7^), higher total cortical surface area (β = 0.154, SE = 0.021, *p* = 5.90x10^-12^), and (nominally) thicker cortex on average (β = 0.073, SE = 0.037, *p* = .049). Higher *g* was marginally associated with greater overall concave curvature (β = 0.080, SE = 0.005, *p* = 6.20x10^-60^) although, as shown below, the direction and magnitude of the association substantially depended on region (range vertex-wise β = -0.10 to 0.09) Average sulcal depth was not associated with *g* (β = 0.018, SE = 0.025, *p* = .472). This appears to be due to regional variation in the direction of effects, which cancel each other out (range vertex-wise β = -0.12 to 0.13; see *Figure 2*).

**Figure 2.**
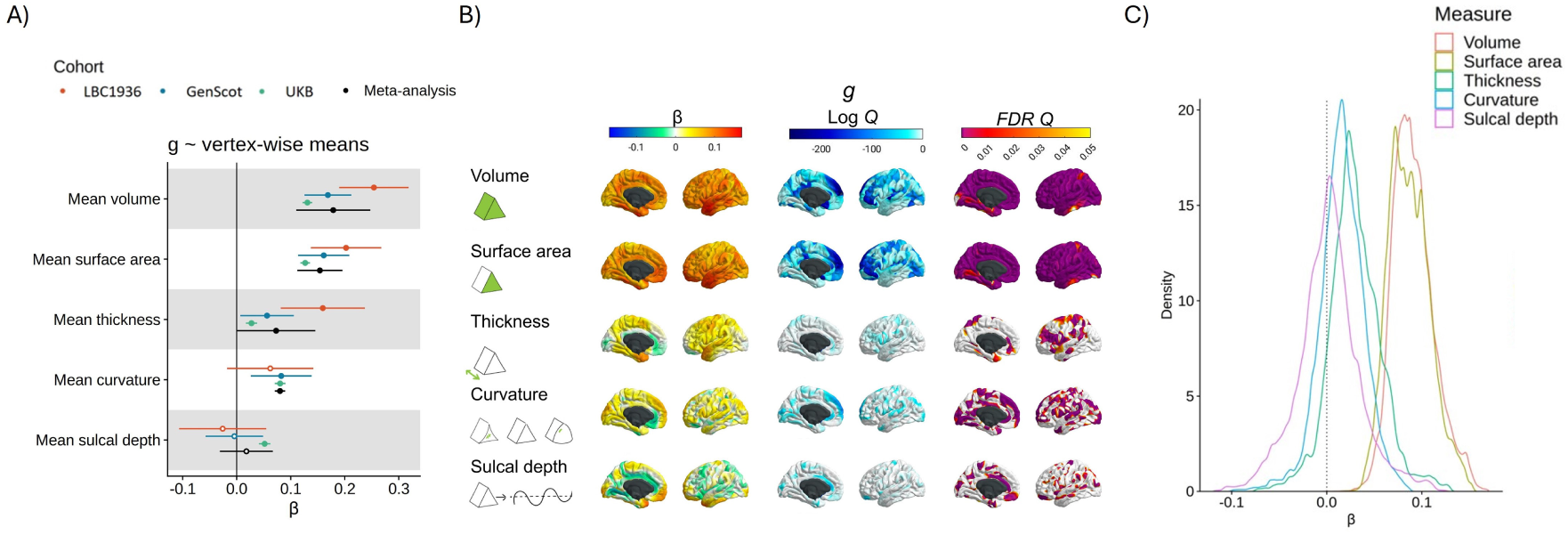
Vertex-wise g-morphometry associations. Figure 2 note A) Associations between g and cortex-level means for all 5 morphometry measures, for the 3 cohorts (UKB, GenScot and LBC1936) and the meta-analysis. B) Vertex-wise g associations, mapped to the cortex. The lower scale limit for log Q maps is set at the minimum available value for any morphometry measure (which is -263.24, or FDR Q = 4.75x10-115). C) Density distributions for the meta-analysed g ∼ morphometry associations for the 5 measures of morphometry (volume, surface area, thickness curvature and sulcal depth). The vertical dotted line marks β = 0.

#### 3.1.2 Vertex-wise *g*-morphometry associations: cross-cohort replicability

We ran vertex-wise *g-*morphometry analyses in each of the three cohorts. We used a 20 mm FWHM smoothing tolerance, which provided a good balance between noise reduction and loss of fine-grained cortical information (*Supplementary Analysis 1*). The patterning of associations between general cognitive function and brain cortical measures showed good between-cohort spatial agreement with moderate-to-strong correlations (see *Table 2*, correlations ranged from Pearson’s *r =* 0.174 to 0.581, all *p* < 2.2x10^-16^). The mean between-cohort spatial correlation for *g* profiles was *r* = 0.424, SD = 0.132, which provides further evidence for the utility of *g* for replicable brain-behaviour analyses ( ^24, 23^). Note that even for traits that have high reliability like sex, the between-cohort correlation is not *r* = 1 (*r* = 0.710, SD = 0.073, see *Table 1* for details). The lowest *g-*association agreements involved the LBC1936, which has an older and narrower-age-range compared to the other two cohorts (mean age = 72.67 years, *SD =* 0.71). Spatial correlations between GenScot and UKB were all *r* > 0.345. Notably, the magnitude of the associations between vertex-wise cortical measures and *g* did not change significantly across mean cohort age groups (there were no between-cohort age moderation effects, *FDR Q* > .05). The *g-*association maps for each cohort are shown in *Figures S*7 to 11 and the density distributions of the β values are summarised in *Figure S*12. Associations between subcortical and global volumes and *g* found in the current work are presented and discussed in detail in *Supplementary Analysis 2*.

**Table 2.** Spatial agreement across cohorts in the patterning of vertex-wise associations with g, age, and sex for cortical volume, surface area, thickness, curvature and sulcal depth. All p < 2.2x10^-16^. Table 2 Note Pearson’s r is shown, indicating a spatial correlation of vectors between each pairwise combination of cohorts (LBC1936, GenScot, and UKB). The vector for each cohort is a list of standardised β at each cortical vertex, denoting the cortex-wide association between morphometry (volume, surface area, thickness, curvature and sulcal depth) and g, or age, or sex.

| Measure | *g* |  |  | Age | Sex |  |  |
| --- | --- | --- | --- | --- | --- | --- | --- |
| Cohorts | LBC1936 | GenScot- | UKB- | GenScot- | LBC1936 | GenScot- | UKB- |
|  | -GenScot | UKB | LBC1936 | UKB | -GenScot | UKB | LBC1936 |
| Volume | 0.177 | 0.435 | 0.254 | 0.553 | 0.765 | 0.780 | 0.749 |
| Surface area | 0.292 | 0.579 | 0.391 | 0.785 | 0.627 | 0.723 | 0.630 |
| Thickness | 0.455 | 0.539 | 0.579 | 0.438 | 0.723 | 0.622 | 0.583 |
| Curvature | 0.273 | 0.345 | 0.297 | 0.625 | 0.681 | 0.781 | 0.677 |
| Sulcal depth | 0.581 | 0.538 | 0.452 | 0.663 | 0.718 | 0.847 | 0.742 |

#### 3.1.3 Vertex-wise *g*-morphometry associations: meta-analysis results

We further meta-analysed the *g-*vertex associations with random effects models (mapped to the cortex in *Figure 2* and shown in extended detail in *Figures S*13 and S14). Qualitative summaries of the cortical regions with the strongest positive and negative associations for *g* are in *Table S1*2. Across volume and surface area (respective β ranges = < 0.001 to 0.17, and 0.01 to 0.15), there were positive associations in lateral temporal, lateral frontal and parietal regions of the cortex. These loci are broadly consistent with the P-FIT (^2^) and other results from single-cohort analyses. These results offer substantially greater spatial fidelity than prior ROI-based analyses.

These meta-analyses also provide novel information about cognitive-cortical associations. For thickness, some regions had positive associations and others had negative associations. These associations (β range = -0.08 to 0.13, *M* = 0.03, SD = 0.03) were most strongly positive in the temporal pole and entorhinal cortex and were most strongly negative in the anterior cingulate, medial orbitofrontal and medial occipital regions, where a thinner cortex predicted higher *g*. Curvature and sulcal depth tended to be absent in prior ROI-based analyses, and so have not been considered in detail in reviews and previously published meta-analyses (e.g., ^92^). For curvature (β range = -0.10 to 0.09, *M* = 0.02, *SD* = 0.02), higher *g* is associated with more concave vertices in medial frontal and medial occipital regions and more convex vertices in the anterior cingulate. Lastly, for sulcal depth itself, our vertex-wise results provide regional detail beyond the null association found when only a global measure was used. There was substantial heterogeneity in regional associations (β range = -0.12 to 0.13, *M* = <0.01, *SD =* 0.03). The results suggest that, relative to the whole cortex, deeper vertices in the medial frontal, temporal pole and parieto-frontal regions are associated with higher *g* and less deep vertices in the cingulate and hippocampal gyrus are associated with higher *g* (see Supplementary Table 12).

#### 3.1.4 Vertex-wise *g*-morphometry associations: agreement between morphometry measures

There were different regional association patterns for the 5 morphometry measures (see *Table 3*, and *Table S*11 for the absolute β value correlations). For example, surface area and thickness had negative and non-significant spatial agreement correlations with each other for both the *g* and age analysis (see *Table 2*, *g: r =* -0.182, *p_spin* = .252; age: *r =* -0.265, *p*_spin = .462). This result is consistent with previous findings that surface area and thickness associations are spatially, phenotypically and genetically distinct (F^93, 94, 95, 96^). The current results agree with the previous findings that patterns of *g* associations are not consistent between different morphometry measures (e.g., ^3, 4^). This serves as a reminder that *g*-morphometry associations do not simply tell us where *g* is in the brain; rather, they each index a conflation of multifarious biological properties which also vary by brain region. The differential nature of these *g*-morphometry associations might be explained by different underlying neurobiological factors of the brain (which we discuss further in section 3.4).

**Table 3.** Correlations (r) between directional (not absolute) g-associations for the 5 vertex-wise measures. Permutation-based p-values are available for g correlations in Table S13, and correlation charts are shown in Figure S15. If the p-values are < .05, they are presented in bold font.

|  |  | Volume | Surface area | Thickness | Curvature |
| --- | --- | --- | --- | --- | --- |
| *g* | Volume |  |  |  |  |
|  | Surface area | **0.656** |  |  |  |
|  | Thickness | **0.442** | -0.182 |  |  |
|  | Curvature | **-0.230** | **-0.163** | -0.069 |  |
|  | Sulcal depth | -0.122 | 0.085 | -0.109 | **0.176** |
| Age | Volume |  |  |  |  |
|  | Surface area | **0.602** |  |  |  |
|  | Thickness | **0.345** | -0.309 |  |  |
|  | Curvature | **-0.266** | -0.105 | 0.020 |  |
|  | Sulcal depth | 0.194 | **0.452** | **-0.342** | -0.111 |
| Sex | Volume |  |  |  |  |
|  | Surface area | **0.771** |  |  |  |
|  | Thickness | **0.600** | 0.169 |  |  |
|  | Curvature | **-0.174** | **0.139** | **-0.298** |  |
|  | Sulcal depth | 0.041 | **0.237** | -0.105 | **0.410** |

#### 3.1.5 Vertex-wise *g*-morphometry associations: within-region correlations

Within-region correlations show that 21/34 regions had a negative correlation between *g*-surface area and *g*-thickness and the top 5 regions with negative correlations, *r* range = -0.86 to -0.65 are: lateral orbitofrontal, caudal middle frontal, pericalcarine, rostral middle frontal, and temporal pole (see *Figure 4*). There were 13/34 regions with positive correlations, and the top 5 regions for which *g*-surface area are positively associated with *g*-thickness are the inferior parietal region, caudal anterior cingulate, lateral occipital, transverse temporal and frontal pole (*r* range = 0.355 to 0.785). These results show that the concordance between these two maps has a considerable amount of variation across the cortex, which is not possible to tell from the overall cortical correlation (*r* = -0.182).

### 3.2 Brain regions most related to *g* are those most susceptible to ageing

In addition to the meta-analytic *g*-morphometry association maps, we similarly calculated meta-analytic maps of associations of age, and sex with cortical morphometry. *Figure 3* shows how these age and sex associations map to the cortex, and *Table S*12 provides a qualitative description of the cortical regions that have the most strongly positive, and most strongly negative β values for each measure for age and sex associations. The *g*, age and sex association maps are in the same analysis space (fsaverage), and so we can quantitatively compare their spatial patterning across the cerebral cortex.

**Figure 3.**
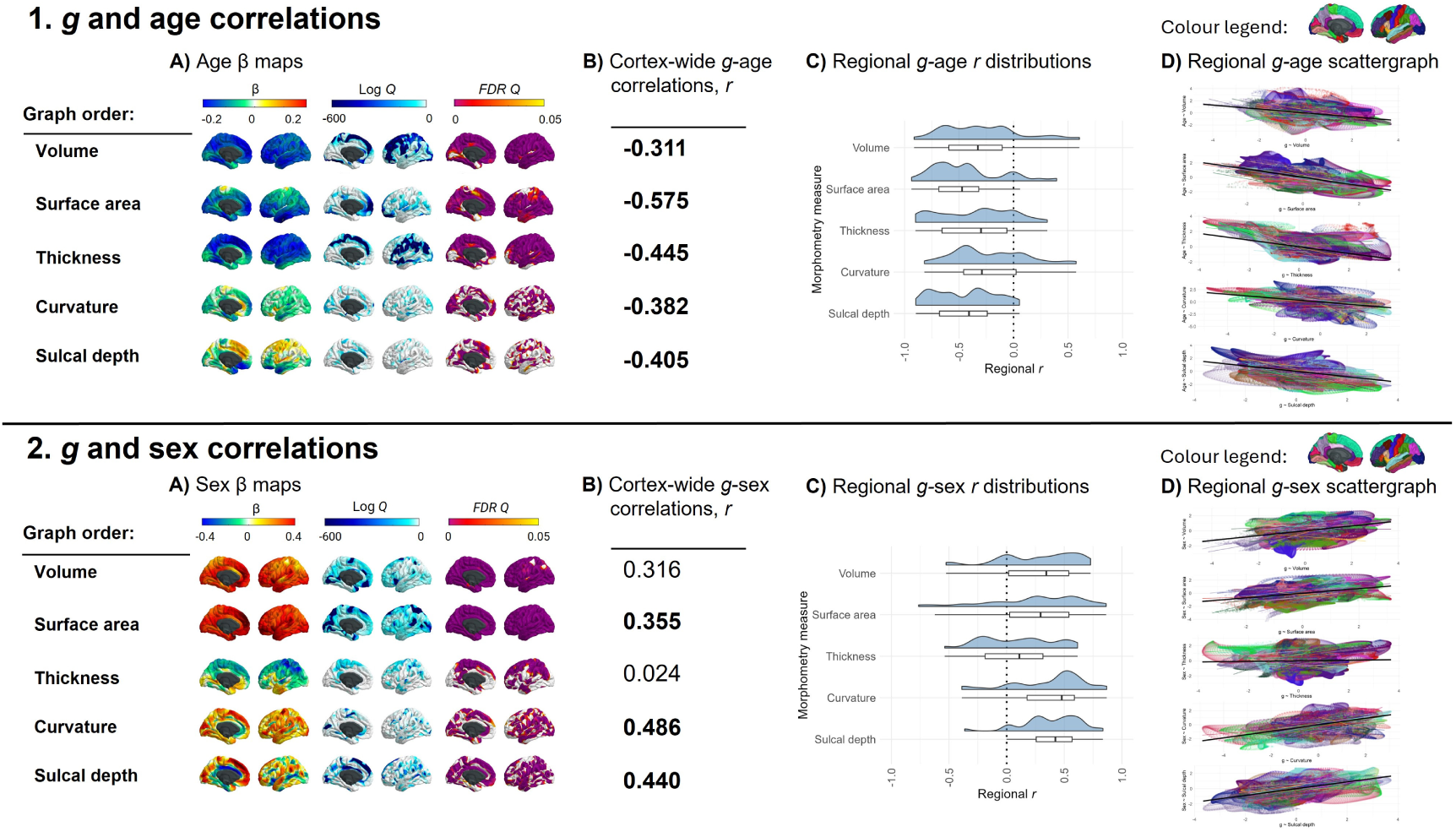
g-age and g-sex correlations. Figure 3 note A) Age (1) and sex (2) associations mapped to the cortex. Some FDR Q values were estimated to be zero, and these have been set to the closest minimum that was successfully calculated. For age and sex, the log Q limit was set at -704.3499, which is FDR Q = 1.273x10^-306^ (Note that the log of Q = .05, a typical α significance threshold, is -2.9957). B) The cortex-level spatial correlations between the g-morphometry associations with the age-and sex-morphometry associations. Note g-sex volume and g-sex thickness had p_spin values > .05, but for all others p_spin < .05. C) Distributions of regional correlations. D) Scattergraphs showing the cortex-wide correlations (representing the numbers in B) in black and showing whether and where that overall spatial agreement holds for different regions (colours represent the 34 paired left/right Desikan-Killiany regions) of the brain.

The vertex-wise age associations show that older people tend to have a smaller cortex in terms of volume and surface area, and most of the cortex also thins with age; . Frontal, lateral temporal and parietal regions are among those most strongly negatively associated with age. For curvature, vertices in the insula tend to be more concave with age, whilst most of the rest of the cortex sees an increase in convex vertices which is consistent with previous findings that show that cortical gyrification decreases with age (^105^). For sulcal depth, the anterior cingulate gyrus, medial frontal region and insula become increased sulcal depth as age increases, and the medial orbitofrontal, posterior cingulate, and lateral orbitofrontal regions are less deep as age increases.

We use these data to quantitatively assess prior observations (which have arisen mostly from qualitative inferences from disparate publications) that those parts of the brain most susceptible to ageing are also those most strongly implicated in our most complex thinking skills (^97^). We tested the spatial agreement of the vertex-wise associations for *g* with those for age (i.e., g-volume with age-volume etc.). The results show that, as previously qualitatively observed (^98, 19, 99^), regions of the brain most associated with *g* are also those that decline most with age: spatial correlations range from *r =* -0.311 to -0.575. These overall spatial correlations broadly held across most regions of the brain (mean number of negative correlations across measures = 28.8/34 regions; *Figure 3C*): for most regions, vertices associated with higher *g* tend to exhibit more ageing-related shrinkage and thinning. The 5 regions with the strongest negative correlations for age-volume and *g-*volume comparisons were the transverse temporal, isthmus cingulate, frontal pole, caudal anterior cingulate and superior temporal regions (*r* range -0.660 to -0.912). The correlation of age-cortex and *g-*cortex associations across all 46 global and subcortical measures was *r =* -0.860, *p =* 2.86x10^-13^ (see *Supplementary Analysis 2* for extended analyses). These findings are compatible with previous findings that present brain, age and *g* associations (e.g., ^19, 100, 101^). For example, this finding could be linked to the “last-in-first-out” hypothesis of ageing, whereby the neocortical regions that are responsible for more complex cognition mature later in development, and are also more vulnerable to ageing, which might be related to the high degree of dendritic plasticity and remodelling required for successful functioning (^102, 103^).

### 3.3 Sex differences paradox may be due to a compensatory volume-gyrification trade-off

The vertex-wise profiles for sex associations show that, across the cortex, males tend to have a larger volume and surface area of frontal regions than females. Females tend to have thicker superior frontal and parietal regions than males, although lateral temporal regions are thicker in males than females. Males tend to have generally more concave curvature across the cortex, compared to females, and increased sulcal depth, particularly in medial frontal regions.

Different brain regions have been shown to differentially mediate associations between sex and cognitive performance (e.g., for volume, the mediation % for verbal and numeric reasoning has been shown to range from 0.9% in the cuneus to 29.1% in the superior temporal region, in a sample of 5216 UKB participants) (^104^). Here, we also tested the strength of spatial correlations between sex and *g* vertex-wise cortical maps. The correlation between sex-and *g-*associations for subcortical and global regions was *r =* 0.305, *p =* .0496 (details available in *Supplementary Analysis 2*). For vertex-wise cortical spatial correlations, regions that tend to be larger/more concave/deeper in males than females also tend to be more associated with *g* (*r =* 0.310 to *r* = 0.486), but this was a considerably weaker, non-significant result for *g-*thickness (*r =* 0.024, *p_spin* = .912). The weaker result for thickness is understood better by looking at the within-region analysis. For the other 4 morphometry measures, the majority of *g-*sex brain map correlations are in one direction (positive, see *Figure 3*, 2D). In contrast, for thickness, there is more of a balance between positive and negative correlations. For some regions, a higher *g* is associated more with a thicker cortex in males (top 5 correlations, *r* range = 0.50 to 0.62: pericalcarine, isthmus cingulate, middle temporal, superior temporal, posterior cingulate), whereas in others, a higher *g* score is associated with thicker cortex in females (top 5, *r* range = -0.24 to -0.54 : frontal pole, paracentral, superior parietal, lateral occipital, superior frontal).

These data offer a valuable new quantitative insight into a well-documented paradox: although global brain volume exhibits clear sex differences, with mean brain volume differing significantly between males and females, these structural disparities do not translate into measurable differences in cognitive functioning between the sexes. One hypothesised explanation for this paradox involves compensatory mechanisms that mitigate volumetric sex differences, such as increased gyrification (^104^). Here, there appears to be direct quantitative evidence of that: we found that brain regions that were largest in males were also more convex in females (*r* = -0.174*, p_*spin = .016 between volume and curvature for sex). Convex vertices are associated with greater gyrification which is generally a sign of a younger, healthier brain (^105^). For this sex ∼ volume, sex ∼ curvature comparison, there were negative within-region correlations for the majority of regions (28/34 regions, see *Figure 4B*), with the top 5 correlations *r* range = -0.77 to -0.81 for the caudal middle frontal, rostral anterior cingulate, pars triangularis, paracentral, and temporal pole regions.

**Figure 4.**
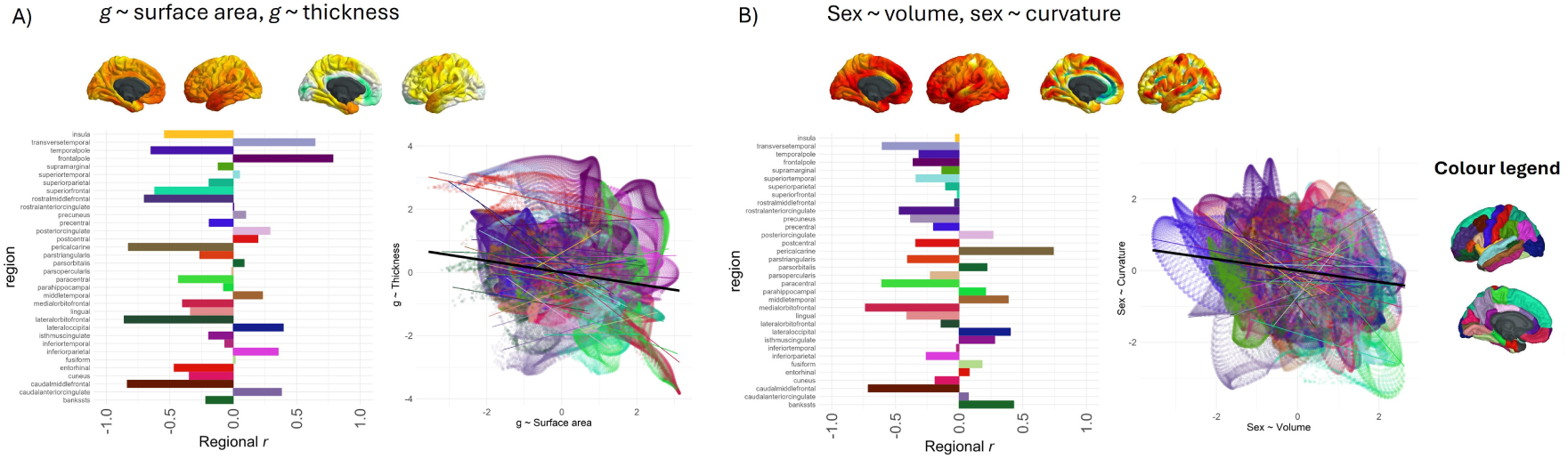
Examples of within-region spatial correlations. Figure 4 note Within-region vertex-wise spatial correlations for A) g ∼ surface area and g ∼ thickness (overall r = -0.182) and B) sex ∼ volume and sex ∼ curvature (overall r = -0.174). The overall r is shown with a black line, and the 34 paired Desikan-Killiany regions are plotted according to the colour legend on the right-hand side of the plot-These results offer regional underpinnings of cortex-wide associations. Assessing within-region correlations allows identifying relative strengths of spatial correlations in different regions across the cortex, as well as the homogeneity of the effects.

### 3.4 Neurobiological correlations between g and brain profiles - what is distinctive about regions associated with g?

We found widespread cortex-wide spatial correlations between *g*’s brain morphometry associations and 33 neurobiological cortical spatial profiles (*Figure 5*). This represents the most detailed compendium of shared spatial signatures between the structure of cognitively-relevant brain regions and, microstructural, macrostructural, functional and receptor densities to have been assembled at high regional fidelity. *g*-volume and *g*-surface area association maps were significantly correlated respectively with 14 and 15 neurobiological profile maps. The neurobiological correlations of *g-*volume and *g-*surface area are highly correlated (*r* = 0.919) suggesting that these two measures are highly similar in their relationships to underlying spatial characteristics. The results indicate that regions of the cortex where larger volume or surface area is more strongly associated with better general cognitive functioning were also those regions that show, for example, lower metabolic activity at rest (*r* = -0.41 and -0.29, respectively), lower cortical myelination (T1/T2 contrast, *r* = -0.35 and -0.47, respectively), higher receptor densities (5HT1a, 5HT2a, 5HT4, 5HT6, D1, D2 M1, VAChT, CB1, MU, NMDA, *r* range = 0.25 to 0.60), and also show significant co-localisation with the primary axis of cortical gene expression (*r* = -0.44 and - 0.48, respectively). The negative association for T1/T2 contrast-derived myelination might be explained as the most highly myelinated areas of the brain are not those involved in higher order cognition but rather are areas receiving large volumes of sensory input such as the primary motor, somatosensory, auditory and visual cortices. Myelination decreases with distance from these regions ^106^. Additionally, higher T1/T2 ratios have previously been associated with poorer outcomes such as Alzheimer’s disease (^107^) and, as discussed earlier, this ratio is perhaps not a good proxy for myelination. Neurobiological profiles without any cortex-wide correlations with *g*’s brain associations were cytoarchitectural staining similarity gradient 1, functional connectivity similarity gradient 2, 5HT1b, A4B2. However, this does not suggest that there are no meaningful spatial correlations between *g*-morphometry profiles and these neurobiological profiles at the regional level (see *Figure* S22 for an extended version of *Figure 5*, with distributions of the within-region correlations, and *Figure*s S26 to S30 for further detail).

**Figure 5.**
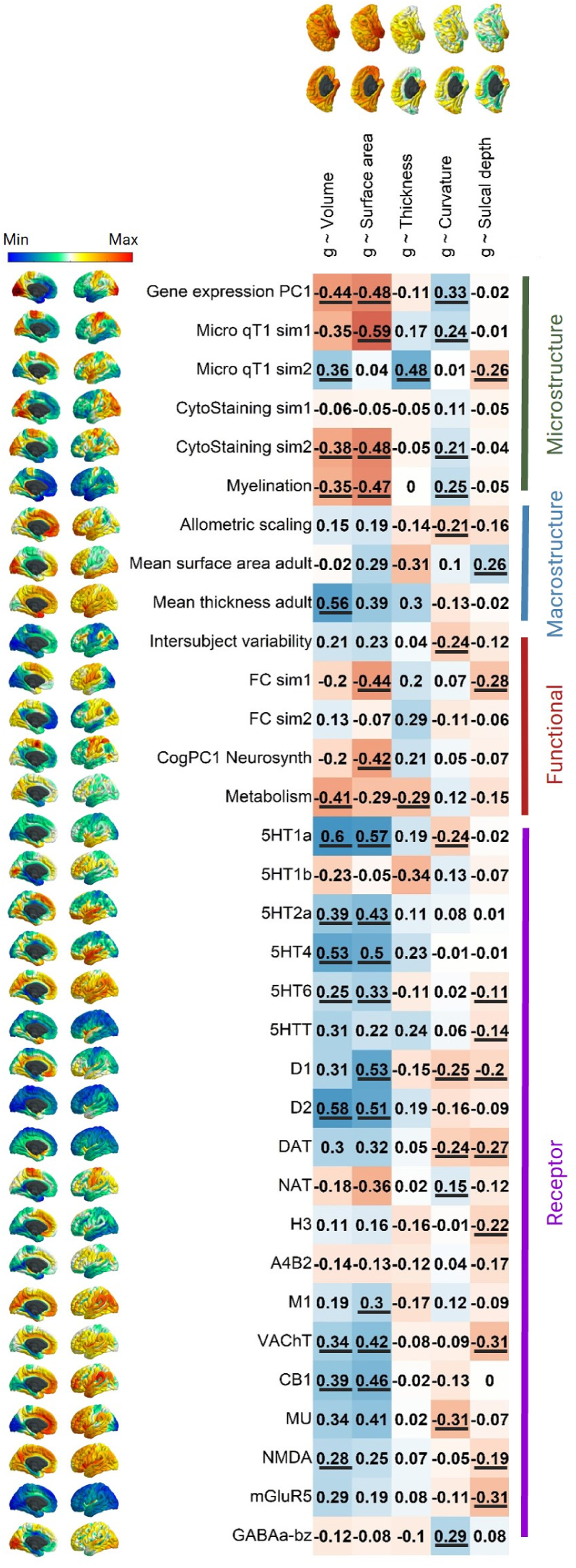
Vertex-wise spatial correlations between g-morphometry associations and 33 neurobiological cortical profiles. Figure 5 note Spatial correlations with p_spin values < .05 are underlined. An extended version of this Figure is available in Figure S22, showing the underlying regional correlation summaries, like those in Figure 4.

As reported in the analyses in section 3.1.4 above, there was a negative and non-significant cortex-wide spatial correlation between *g* ∼ surface area and *g* ∼ thickness (*r* = -0.182, *p_spin* = .252). In the current analyses, our cortical map of mean surface area (i.e., one of the neurobiological profile maps) was positively associated with *g* ∼ surface area (*r* = 0.29, *p_spin* > .05) and negatively associated with *g* ∼ thickness (*r* = -0.31, *p_spin* > .05) and these two correlations appear to cancel each other out in mean surface area’s association with *g* ∼ volume (*r* = -0.02, *p_spin* > .05). Neurobiological profiles might offer some further insights into these relationships between *g ∼* surface area and *g* ∼ thickness. While microstructure gradient 1 was correlated with *g-*surface area (*r* = -0.59), microstructure gradient 2 was correlated with *g-* thickness (*r =* 0.48). A similar pattern occurred for functional connectivity similarity gradients where the first one was significantly correlated with *g*-surface area (*r* = -0.44), and the second had a moderate correlation with *g*-thickness (*r* = 0.29, although *p_spin* = .205 The tendency of g-surface area and g-thickness to align with different microstructural and functional similarity gradients may help explain why their cortical spatial patterns are spatially distinct, potentially reflecting their unique phenotypic and genetic characteristics (^93, 94, 95, 96^).

Regions of the brain where volume and surface areas were most strongly related to *g* were also those that density show particularly high receptor density across multiple neurotransmitters (5HT1a, D2, D1, 5HT4, CB1, VAChT, and 5HT2a; *r* range = 0.34 to 0.59). For the other *g-* morphometry associations, cortex-level spatial correlations were generally smaller and positive, with some exceptions. For example, there is a relatively large negative correlation between *g-* curvature and MU (*r =* -0.31, suggesting that regions with a higher density of MU receptors are those for which more convex vertices are associated with higher *g*) and between *g-*sulcal depth and VACHT and mGluR5 (both *r =* -0.29, suggesting that regions with a higher density of VACHT and mGluR5 receptors tend to be those for which a higher *g* is associated with a more gyral vertex). At the cortex-wide level, there were no *p_spin* significant associations between the *g*-thickness map and any neurotransmitter receptor profiles, although, as shown in *Figure* S28 (and reported in the Supplementary Tabular Data File), this appears to be because there was a mix of positive and negative correlations at the regional level, which likely cancel each other out.

### 3.5 Four major dimensions explain the majority of spatial variation across 33 cortical properties

Since there were some qualitatively observable consistencies in spatial maps across the 33 cortical properties presented in *Figure 5*, we conducted a spatial component decomposition using principal components analysis (PCA) to quantitatively identify any underlying spatial similarities (i.e., statistical ‘dimensions’) that were shared across multiple maps (*Figure 6*). We conducted the PCA across all 33 maps. It might be argued that one should reduce the dimensionality of the N = 19 receptor maps first. However, there was no evidence that those receptor maps were more similar to each other than to all other maps, i.e., the absolute correlations within neurotransmitters, and those within other types of maps were not significantly different from each other (t = -0.198, *p* = .843, neurotransmitter maps’ mean |*r*| = 0.371, *SD* |*r*|= 0.237, other maps’ mean |*r*| = 0.367, *SD* |*r*| = 0.267).

**Figure 6.**
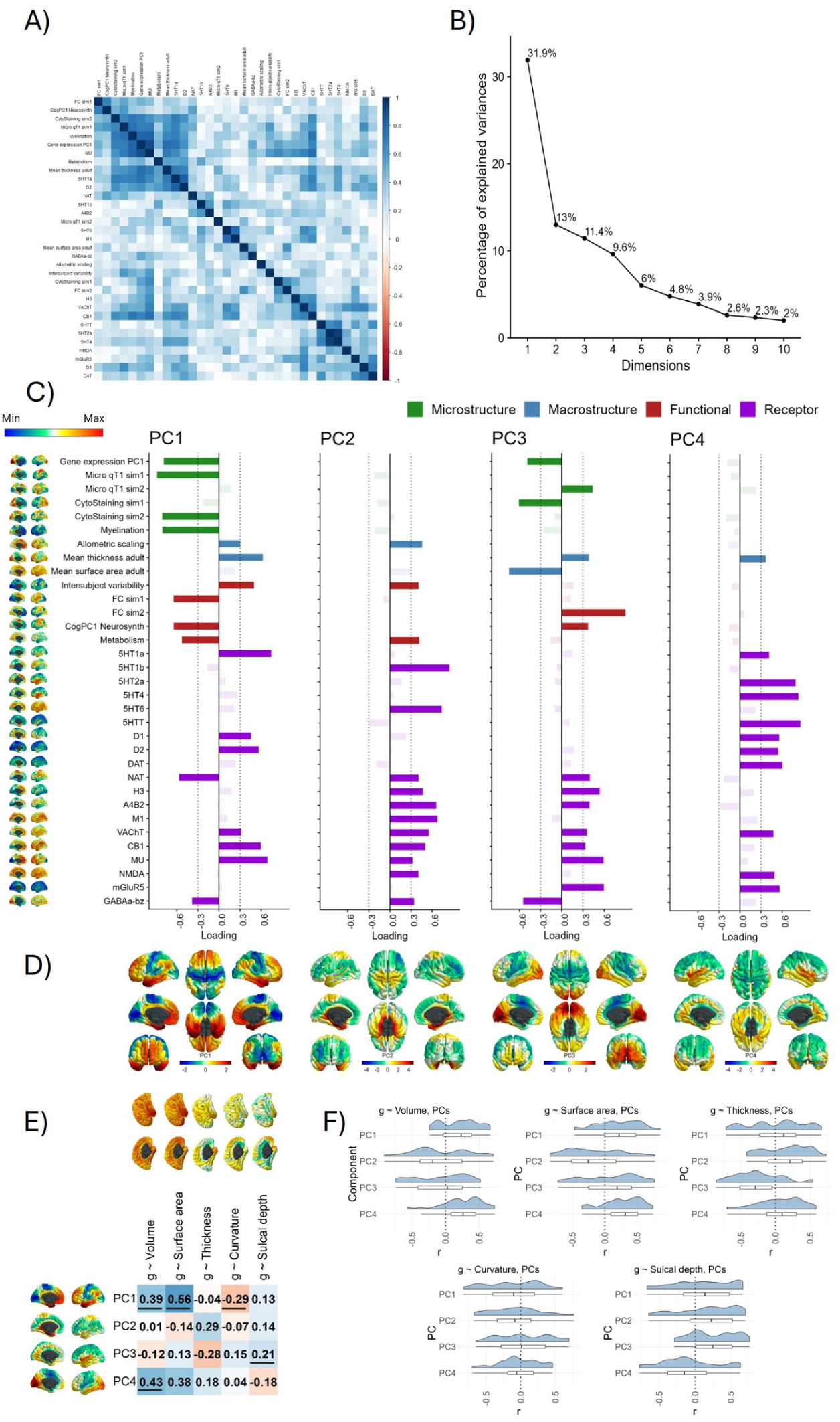
Many neurobiological profiles converge on four spatial dimensions which are spatially correlated with g-morphometry maps. Figure 6 note A) A correlation plot between the 33 neurobiological profiles. B) A scree plot showing percentage of explained variance by the first 10 PCs. The first four were extracted with varimax rotation. C) The loadings of each neurobiological profile on the first four components. Loadings < |.3| are shown in a reduced alpha to aid interpretations. D) PC scores mapped onto the cortex. E) Correlations of four PCs with g ∼ morphometry associations (with spin test < .05 underlined). F) The distributions of within-region g ∼ morphometry correlations. Note that this plot does not take into consideration the number of vertices in each region.

Consistent with our qualitative observations, the 33 maps shared only four main spatial patterns: the first four components accounted for 65.9% of the variance, and there was a marked inflection point in the variance explained after these four (*Figure 6A* and scree plot in *Figure 6B*). We extracted the first four components with varimax rotation (loadings presented in *Figure 6C*). The first component alone accounted for almost one third of the spatial variance (the loadings of PC1 were similar whether rotated or unrotated; coefficient of factor congruence = 0.900; *Figure S18*). We describe these major dimensions as mapped onto the cortical surface (see Figure 6D), which appear to reflect core organisational principles of the brain’s neurobiology across multiple scales:

- PC1 resembles previously reported latent variables of cortical macrostructure (^108, 78^). Its cortical profile is characterized by a gradient from unimodal sensory input areas (sensorimotor, primary auditory and visual / medial occipital regions) at one end of the scale, and amodal association cortices (medial frontal and temporal regions) at the other (^78^). It captures multiple aspects of neurobiological information with high loadings across microstructure, macrostructure, functional activity, and neurotransmitter receptor density categories. The largest positive receptor loadings were for 5HT1a, MU, CB1, D1 and D2. NAT and GABAa-bz had negative loadings < -0.3.
- PC2 is medial temporal and is most strongly characterized by allometric scaling, intersubject variability and metabolism showing strong parahippocampal localisation. High-loading receptor maps were A4B2, M1, VAChT, CB1, MU, mGluR5 and GABA-bz.
- PC3 is an anterior-posterior component and is associated with functional activity (both FC similarity gradient 2 and CogPC1 Neurosynth) and the first principal component of cortical gene expression, as well as with cytoarchitectural staining and microstructural
similarity profiles. There is a large negative loading for GABAa-bz, and the largest positive loadings for mGlur5, MU and H3.
- PC4 is superior/inferior, with strikingly strong component scores in the insula. It is largely a receptor-based component, with notable positive loadings for several serotonin maps (5HT1b, 5HT2a, 5HT4, 5HTT), and all three dopamine maps (D1, D2 and DAT), as well the two glutamate maps (NMDA, and mGLuR5) and VAChT (acetylcholine).

We correlated these major dimensions of neurobiological organization with the *g*-morphometry association maps (see *Figure 6E*). Notably, PC1 correlations were highest with *g-*volume (*r =* 0.39, *p_spin* =.009) and *g-*surface area (*r =* 0.56, *p_spin* = .002). The strongest cortex-wide correlation for PC2 is for *g-*thickness, although it is not significant in the spin test (*r =* 0.29, *p_spin* = .074). The only spin test *p* value < .05 for g-morphometry associations with PC3 is for sulcal depth (*r* = 0.21, *p_spin* = .049). PC4 had the strongest association with *g-*volume (*r =* 0.43, *p_spin* = .010), which appears to be led by *g-*surface area (*r* = 0.38, *p_spin* = .079) rather than *g-*thickness (*r* = 0.18, *p* = .364).

Within-region analyses results show that significant cortex-wide map correlations are underpinned by homogenous correlations at regional level (see Figure 6F). Neurobiological profiles with null correlations at the cortex level had both positive and negative correlations at the regional level, which cancel each other out but could still reveal important associations of *g*-morphometry maps and core organisational principles of the human brain. These results are presented in detail in Figure S31 and the Supplementary Tabular Data File.

Together, the results show that multiple biological properties covary together in relatively few spatial axes across the cortex. These dimensions represent multi-system neurobiological foundations of individual differences in general cognitive functioning.

## 4 Discussion

This study provides the most definitive cross-cohort characterisation of regional morphometric brain associations with *g* to-date. It demonstrates how such associations vary in strength and direction across the volume, surface area, thickness, curvature and sulcal depth of the cortex. We also provide a compendium of spatial associations between *g*-morphometry profiles and 33 neurobiological profiles and discover that these 33 profiles share four major dimensions of spatial cortical organization. We look at spatial correlations between *g-*morphometry associations and neurobiological profiles to provide insights into the neurobiological mechanisms underpinning our complex cognitive functions.

Using the spatial correlations approach, the current results provide further detail about *g* and brain-neurobiology relationships. Key findings were: 1) We provide the largest to-date meta-analytic vertex-wise associations between brain morphometry and *g* (i.e., morphometry maps) and characterise how they vary across the cortex; 2) Vertices with larger *g* associations tend to be more susceptible to age; 3) Vertices most strongly associated with *g* were largest in males and also more convex in females; 4) *g-*morphometry associations maps substantially overlap with maps of neurobiological properties; 5) The cortical spatial patterning of 33 neurobiological maps can be concisely summarised in 4 PCs, and their correlations with *g*-morphometry patterns are presented.

There were *p_spin* significant correlations between several *g-*morphometry and neurobiological profiles, such as dopamine, serotonin, VACHT, CB1 and NMDA neurotransmitter receptor densities. These neurotransmitter-cognition results are in line with previous reports e.g., dopamine and serotonin have previously been shown to be important for cognitive processes (e.g., ^109, 110, 111^), VACHT dysfunction has been shown to be related to intellectual disabilities and Parkinson’s Disease, as well as to prefrontal cortex functioning (^112,113^), acute CB1 disruption results in a decline in verbal learning and working memory performance ^114^, and NMDA has been selected as a promising target for cognitive enhancement e.g., in dementia (^115, 116^). There was a negative correlation between brain volume and metabolism, which might be explained as the metabolism data were collected a rest, so one might expect these regions to be de-coupled from the cognition-relevant regions we identify here. *g*-volume and *g-*surface area also had moderately strong correlations with PC1 gene expression, in line with our previous work which characterised *g-*morphometry and gene expression profile associations in more detail (^3^). Microstructural and functional similarity gradients had differential correlations between *g*-surface area and *g*-thickness, which could give insights into the mechanisms behind why the spatial profiles of these two *g*-morphometry profiles are different.

The spatial variation of the neurobiological profiles across the cortex was captured in four dimensions which reflect multi-scale organisational principles of the brain that support general cognitive functioning. Neurobiological PC1 resembles previously reported latent variables of cortical macrostructure (^108, 78^). Its cortical profile is characterized by a gradient from unimodal sensory input areas (sensorimotor, primary auditory and visual / medial occipital regions) at one end of the scale, and amodal association cortices (medial frontal and temporal regions) at the other. It captures multiple aspects of neurobiological information with high loadings across microstructure, macrostructure, functional activity, and neurotransmitter receptor density categories. The consistency with previous reports suggests that our applied methods are promising and that the core dimensions of cortical organisation may be replicated across different measures and analysis strategies. PC1 had moderate correlations with *g*-volume and *g*-surface area, suggesting that this dimension of neurobiological characteristics is important for cognition-brain organisation.

In the current study we also offer a framework to extend the increasingly popular method of calculating spatial correspondences between two cortical profiles (usually represented by a single correlation of assumed linear correspondence). Our approach to examine vertex-wise within-region spatial agreement offers important insights about the relative strength of correlations for different regions and the extent of homogeneity of cortex-wide correlations across regions. For correlations between age-brain and *g-*brain maps, the vast majority of within-region correlations are negative. For example, for cortical volume – across regions, vertices for which a higher *g* is associated with higher volume tend to be the same vertices for which a higher age is associated with lower volume (i.e., appears to decline more with age). On the other hand, for example, for the *g* ∼ surface area and *g ∼* thickness comparison, there are 13 regions with positive associations (*r*s M = 0.27, SD = 0.24, range = 0.006 to 0.785) and 21 with negative associations (*r*s M = -0.397, SD = 0.269, range = -0.012 to -0.859), showing that the concordance between these two maps has a considerable amount of variation across the cortex, which is not possible to tell from the overall cortical correlation (*r* = -0.182). Sometimes a cortical correlation might be null because positive and negative within-region correlations cancel each other out. For example, for *g* ∼ thickness, there are no *p_spin* significant cortex-wise correlations with any neurotransmitter receptor density maps, but at the regional level, there are both large positive and negative associations for several receptor types. These within-region correlations cancel each other out at the cortex-level and conceal potentially meaningful spatial correlations between *g* ∼ thickness and neurotransmitters receptor densities.

The extent to which brain-behaviour associations are stable and replicable is a subject of current debate (^117, 118^), and here we formally quantified the extent to which the patterning of associations is stable across cohorts. The results show good cross-cohort agreement, suggesting that it is not always the case that thousands of individuals are required to produce reproducible brain-wide associations. We also provide a critical evaluation of smoothing tolerances for these associations, which suggests that between 10-20 mm FWHM is a good choice across morphometry measures. We further conducted cross-cohort meta-analytic data on subcortical associations with *g,* age, and sex (described in detail in *Supplementary Analysis 2*) which contribute significantly to the literature on this topic.

For our meta-analyses, methods were matched, where possible, between the UKB, GenScot and LBC1936 cohorts (e.g., obtaining brain morphometric measures from FreeSurfer and including multiple different cognitive test scores in our calculation of latent *g* scores). This consistency allowed for direct quantitative comparison between cohorts and leads to improved confidence in the final meta-analytic estimates of *g-b*rain associations. With that said, some differences in MRI data and processing protocols between the three cohorts might differentially affect the cortical surface results: 1) each of the three cohorts used different scanners for MRI acquisition and, although T1-weighted data provides consistent between-scanner measures (^119^), we cannot rule out between-cohort scanner-specific effects; 2) Desikan-Killiany parcellations were visually inspected and manually edited for LBC1936 and GenScot, which would also affect the vertex-wise surfaces, but manual inspections were not carried out for UKB; and 3) different FreeSurfer versions were used for each cohort and are likely to have contributed to some differences in estimations, alongside different types and quantity of cognitive tests. It is therefore encouraging that the spatial correlations were fairly stable and that the meta-analytic results also show significant associations with multi-modal biological data from independent sources.

All cortical maps included in the current analyses were registered to fsaverage space. Registration differences might have had a small impact on the results – e.g., there were a few vertices around the cortical mask that were present in some transformed fsaverage maps, and not in others (e.g. there were fewer vertices included in the cortical mask for the gene expression PC1). However, we only included vertices within the cortical mask across all maps in each analysis, and we would expect the effects of such registration inaccuracies to be small. Additionally, efforts were made to harmonise smoothing tolerances between maps for different data types, but the original sampling density across the cortex was different between maps. Cortical data obtained at lower spatial resolutions (e.g. neurotransmitter density maps, derived from PET data) may have contributed to some uncertainty in our vertex-wise analyses, particularly within smaller cortical structures. The cortex-wide *r* values should thus be interpreted alongside the corresponding *p_spin* values, as this limits the spatial autocorrelation effects that tend to increase with increased smoothing.

Although the spin test goes some way to improve the validity of spatial correlations, it is important to note, too, that using Pearson’s *r* to test spatial correlations of cortical profiles assumes that we are working with linear spatial associations. This is not always the case (e.g., see *Figure* S15), as non-linear patterns are often found. The spin test does not address the heterogeneity of spatial autocorrelation effects across the cortex. Further methods are currently being developed to address this issue and more accurately characterise spatial concordance between cortical maps (^120^). The methods and cortical maps we provide here could aid more detailed investigations into how and where spatial concordance/discordance between neurobiological and brain structural profiles is relevant to brain-behaviour relationships.

The methods used in the current paper rely on all cortical profiles having results at all vertices. The publication and release of summary data for all vertices should be encouraged, not only those vertices that reach certain criteria (e.g., significance thresholds). While for example, cluster-based analyses can be useful for identifying the main regions of interest, presenting results for all vertices provides information about the relative patterning of effects across the whole cortex that can be directly compared with other whole-cortex profiles.

A limitation of this study is that participants were likely to be in relatively good health, as we chose to exclude participants with self-declared neurological issues from the UKB sample. Having said that, our UKB exclusion criteria did not include GP, hospital or death records, so it is likely that some participants with such conditions remain in the UKB sample, and may influence the findings, as some neurological diseases e.g. stroke or brain lesions are likely to affect brain-cognition associations. The GenScot imaging sample was biased to have more participants with past or current depression than would likely be typical in the general population, in line with the initial aims of their study. Depression status was not controlled for in the current analyses, and it is possible that it could affect brain-cognition associations. All three cohorts used to calculate *g*-brain associations are also largely white and northern European, and so it is not clear whether these findings apply in other world regions or ethnic groups. Additionally, whereas the cognitive-MRI data do not include childhood and adolescence (and therefore the results may not relate directly to those parts of the life span), the good adulthood age coverage, absence of age moderation of the meta-analytic estimates within-cohort, and clear agreement across cohorts suggests that the well-powered results capture adulthood brain-*g* correlations. The open-source neurobiological maps that we use here are also limited in terms of generalisation due to sample characteristics, which are also not directly comparable with the cohorts used to calculate *g*-morphometry associations. For example, gene expression PC1 was calculated based on data from 6 donors, aged 24 to 57 years old, which is a small sample, and has a younger age range than the current study (age range = 44 to 84 years old); the cytoarchitectural similarity maps were based on just one donor, and the PET maps tended to be conducted on young healthy adults, with sample sizes ranging from 8 to 204. At this stage, the results should be thought of as being at a high-level overview stage, and be used for hypothesis generation, rather than being taken as direct evidence of brain-behaviour relationships.

As further group-level brain-wide maps are made openly available, the correlational structures between different cortical profiles should continue to be examined and updated. Moreover, the future collection or release of individual-level data across different neurobiological characteristics would enable individual differences analyses, for example, to test the relationship between the density of certain receptor types and *g*-morphometry associations at the individual level. This would allow for more direct associations between neurobiological characteristics and *g*, and longitudinal studies with within-participant cognitive, brain imaging and neurobiological data could also directly reveal whether which neurobiological characteristics underpin the clear similarity between *g* and age-related brain patterns.

## 5 Conclusion

This study advances our understanding of how different neurobiological profiles in the human cortex share spatial patterning with *g*-structural morphometry profiles. We discovered four principal components, which explain 65.9% of the variance across 33 neurobiological profiles, and represent major fundamental axes along which the human cortex is organised. These results offer new perspectives on the neurobiological properties underlying observable brain-cognitive associations. We provide important new data and a framework to study brain-behavioural associations in the future.

## Supporting information

Supplementary Analyses

Supplementary Tabular Data

## Conflicts of interest

None

## CRediT statement

Joanna E. Moodie: Conceptualization, Methodology, Writing -Original Draft, Formal analysis, Visualization; Colin Buchanan: Writing - Review & Editing; Eleanor Conole: Writing - Review & Editing; Anna Furtjes: Writing - Review & Editing; Aleks Stolicyn: Data Curation, Writing - Review & Editing; Janie Corley: Writing - Data Curation, Review & Editing; Karen Ferguson: Writing - Review & Editing; Maria Valdes Hernandez: Data Curation, Writing - Review & Editing; Susana Munoz Maniega: Data Curation, Writing - Review & Editing; Tom C. Russ: Writing - Review & Editing; Michelle Luciano: Data Curation, Writing - Review & Editing; Heather Whalley: Data Curation, Funding Acquisition, Writing - Review & Editing; Mark E. Bastin: Data Curation, Funding Acquisition, Writing - Review & Editing; Joanna Wardlaw: Data Curation, Funding Acquisition, Writing - Review & Editing; Ian Deary: Funding Acquisition, Writing - Review & Editing; Simon Cox: Conceptualization, Data Curation, Project Administration, Resources, Funding Acquisition, Methodology, Writing - Original Draft, Supervision

## Acknowledgements

We thank the participants of the three cohorts (UKB, Generation Scotland and LBC1936) for their participation and the research teams for their work in collecting, processing and giving access to these data for analysis. The UKB research was conducted using the UK Biobank Resource under Application number 10279. We are also thankful to the participants of all the studies involved in generating the neurobiological profiles, and to the people who collected and processed the data and made it openly available. SRC and JEM were supported by a Sir Henry Dale Fellowship, jointly funded by the Wellcome Trust and the Royal Society (221890/Z/20/Z). The LBC1936, supported by the BBSRC & ESRC (BB/W008793/1) (which also supports SMM, JC, and IJD), Age UK (Disconnected Mind project), the Medical Research Council (MR/M01311/1; MR/K026992/1), the US National Institutes of Health (R01AG054628) and the University of Edinburgh. CRB, MEB, IJD and SRC were supported by a National Institutes of Health (NIH) research grant R01AG054628. AF is supported by National Institutes of Health (NIH) grant R01AG073593. TCR is a member of the Alzheimer Scotland Dementia Research Centre funded by Alzheimer Scotland. MCVH and JW are funded by The Row Fogo Charitable Trust Centre for Research into Aging and the Brain (BRO-D.FID3668413). JW is also funded by the UK Dementia Research Institute which receives its funding from UK DRI, funded by the UK Medical Research Council, Alzheimer’s Society, and Alzheimer’s Research UK. AS was funded as part of the Generation Scotland study (Wellcome Trust reference 104036/Z/14/Z), which is led by HW and AM. ELSC is Junior Research Fellow in Applied AI at Lady Margeret Hall, University of Oxford (EPT-AI). For the purpose of open access, the author has applied a CC-BY public copyright licence to any Author Accepted Manuscript version arising from this submission.

## Notes

### Competing Interest Statement

The authors have declared no competing interest.

### Summary of Updates

Supplementary analyses file - reuploaded without tracked changes.

