## Supplementary Figures and Tables for "Brain maps of general cognitive function and spatial correlations with neurobiological cortical profiles"

#### Cognitive descriptives/models

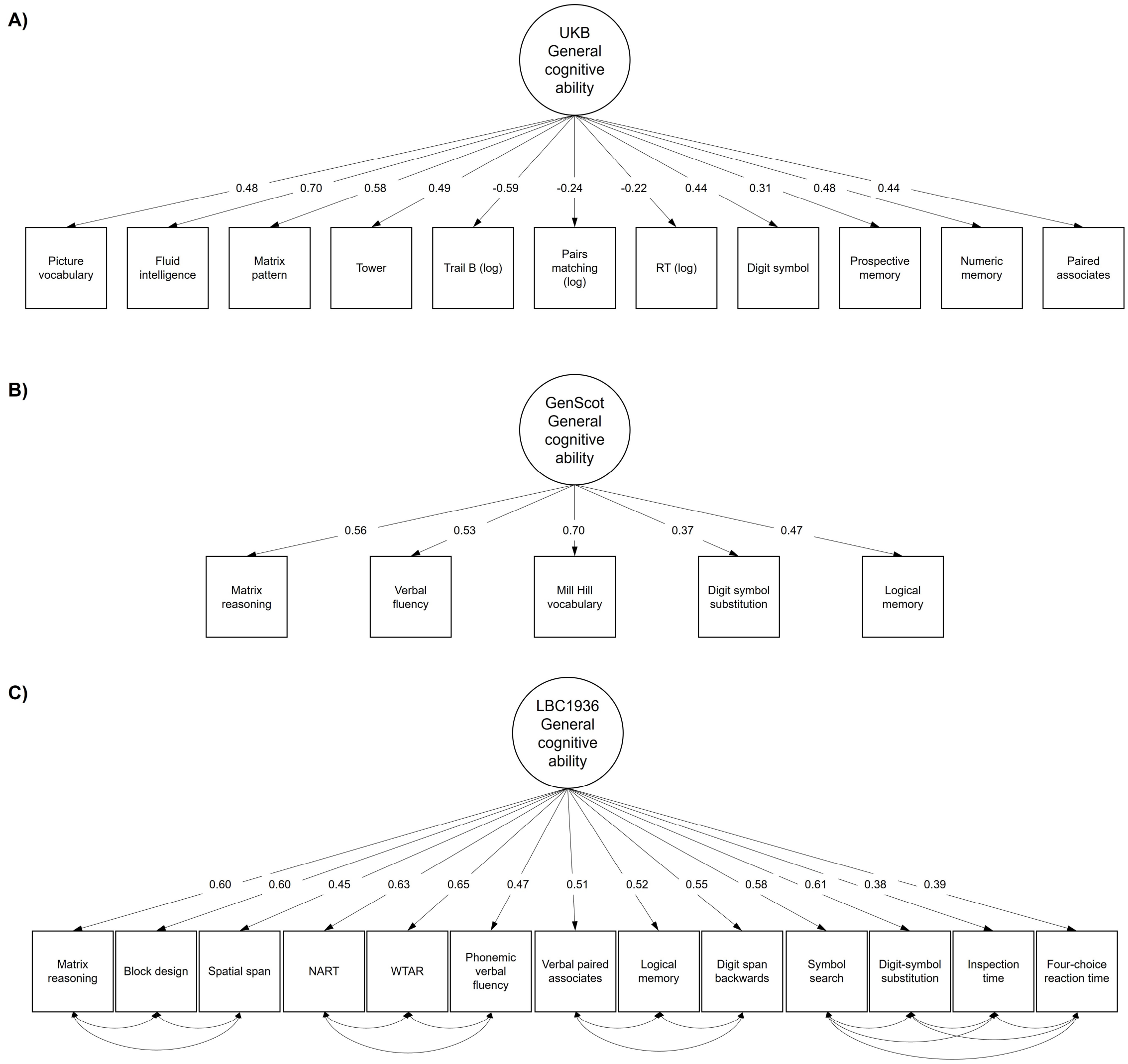


*Figure S1* Simplified path diagrams of cognitive ability latent models alongside density plots of the final cognitive ability prediction *z* scores for A) UKB, B) GenScot, and C) LBC1936. The within-domain residual variances for LBC1936 are in Table S8.

#### Vertex-wise descriptive stats

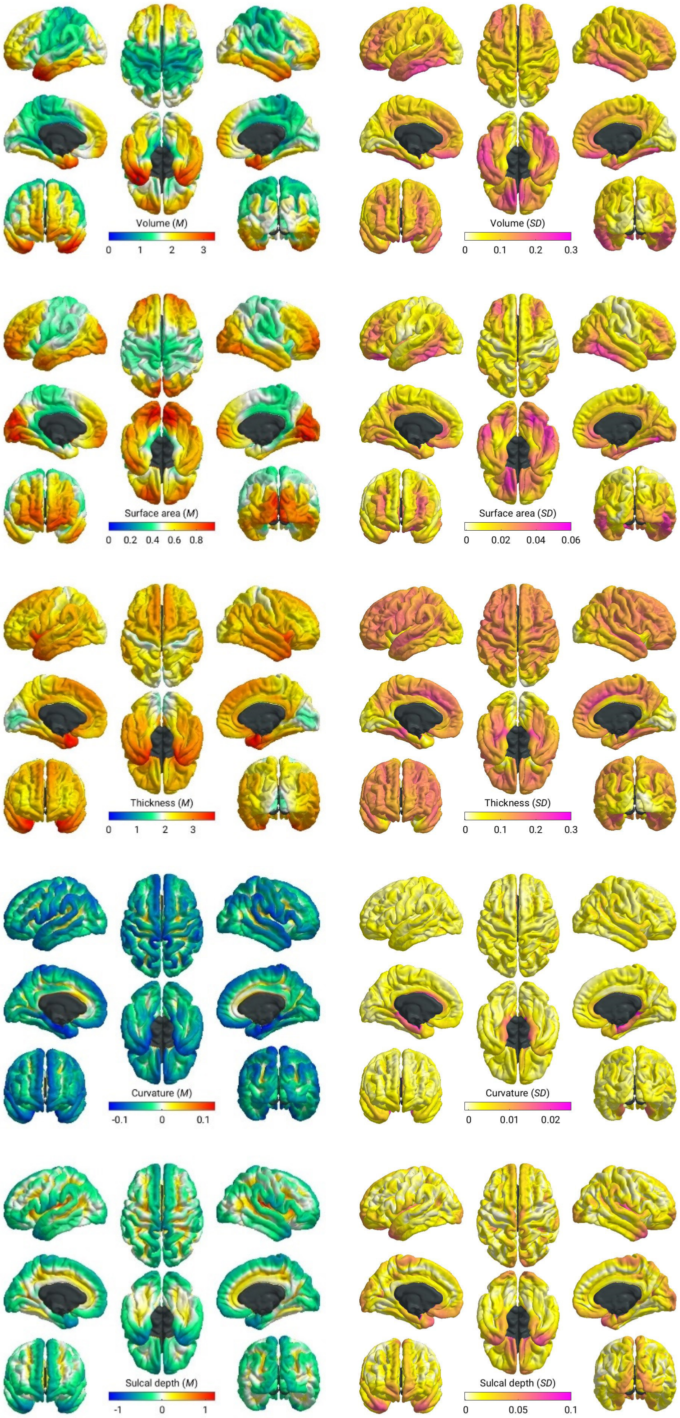


*Figure S2* Meta-analysed mean (left) and *SD* (right) values for each cortical morphometry measure. Mean volume is in $mm^3$, surface area is in $mm^2$, thickness is in mm, curvature in in $mm^{-1}$, sulcal depth is in mm.

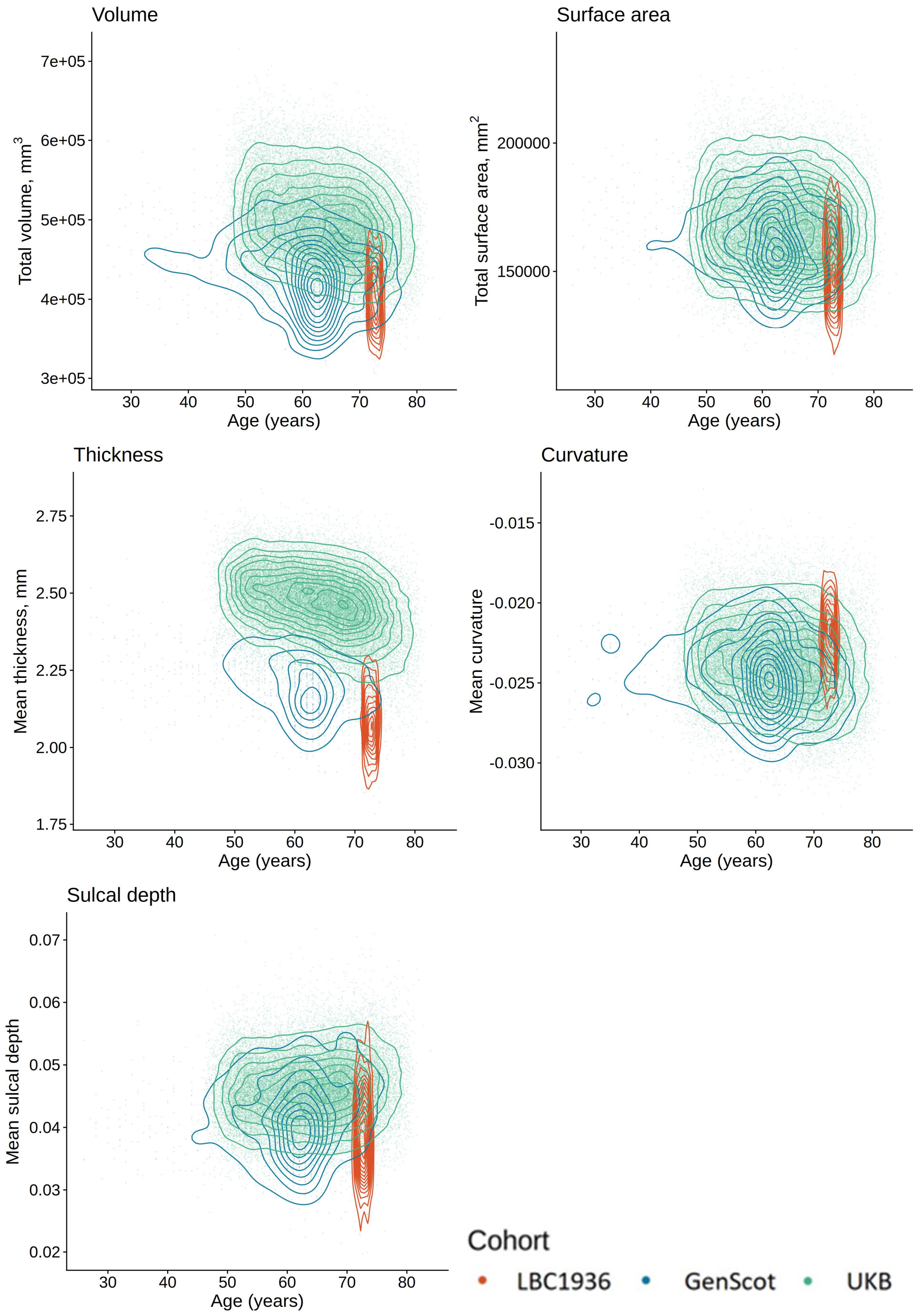


*Figure S3* Raw participant totals/means of the 5 vertex-wise measures, plotted by age and cohort.

#### Smoothing tolerances

LBC1936

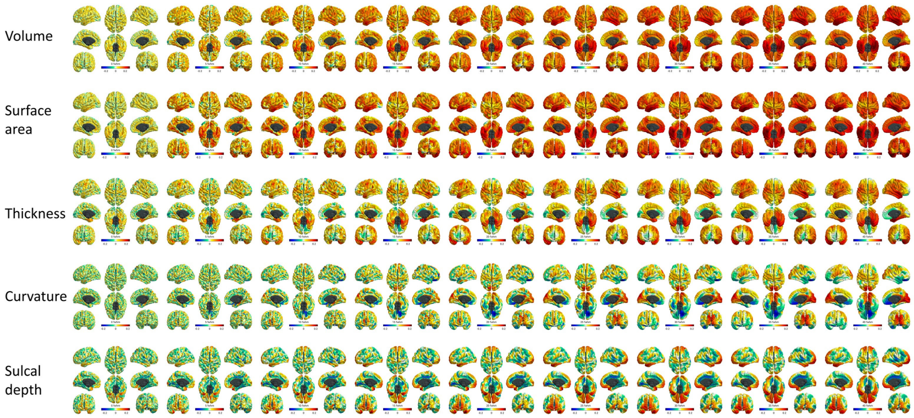


GenScot

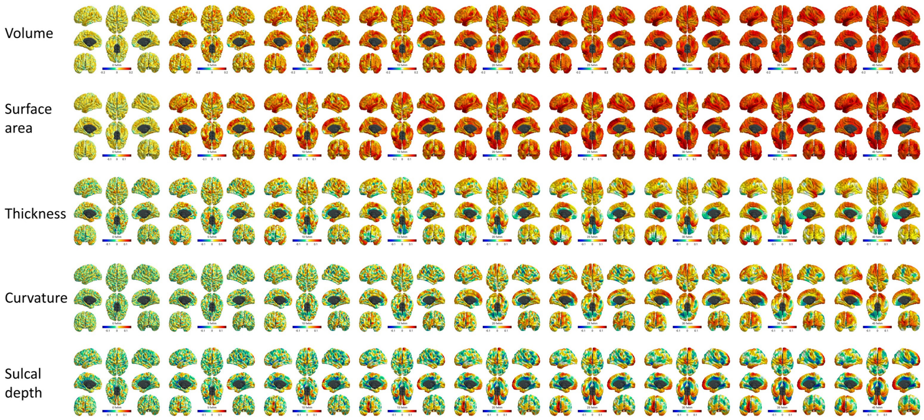


UKB

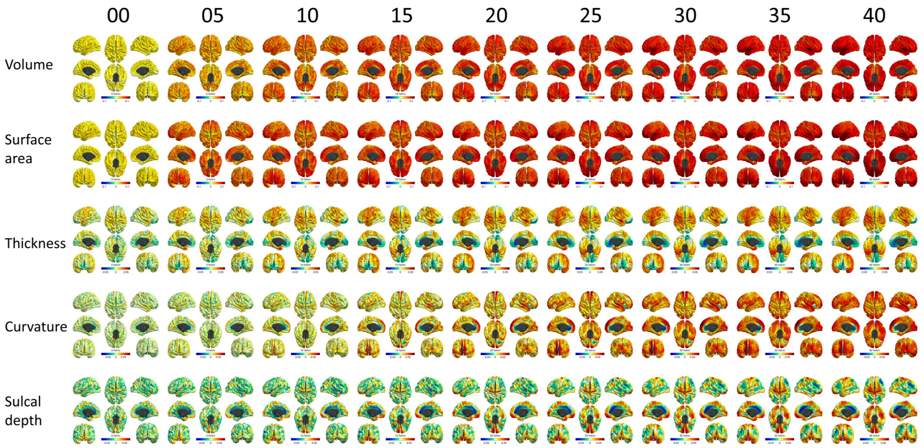


*Figure S4* Maps of g-associations in each cohort at the 9 smoothing tolerances for the 5 morphometry measures.

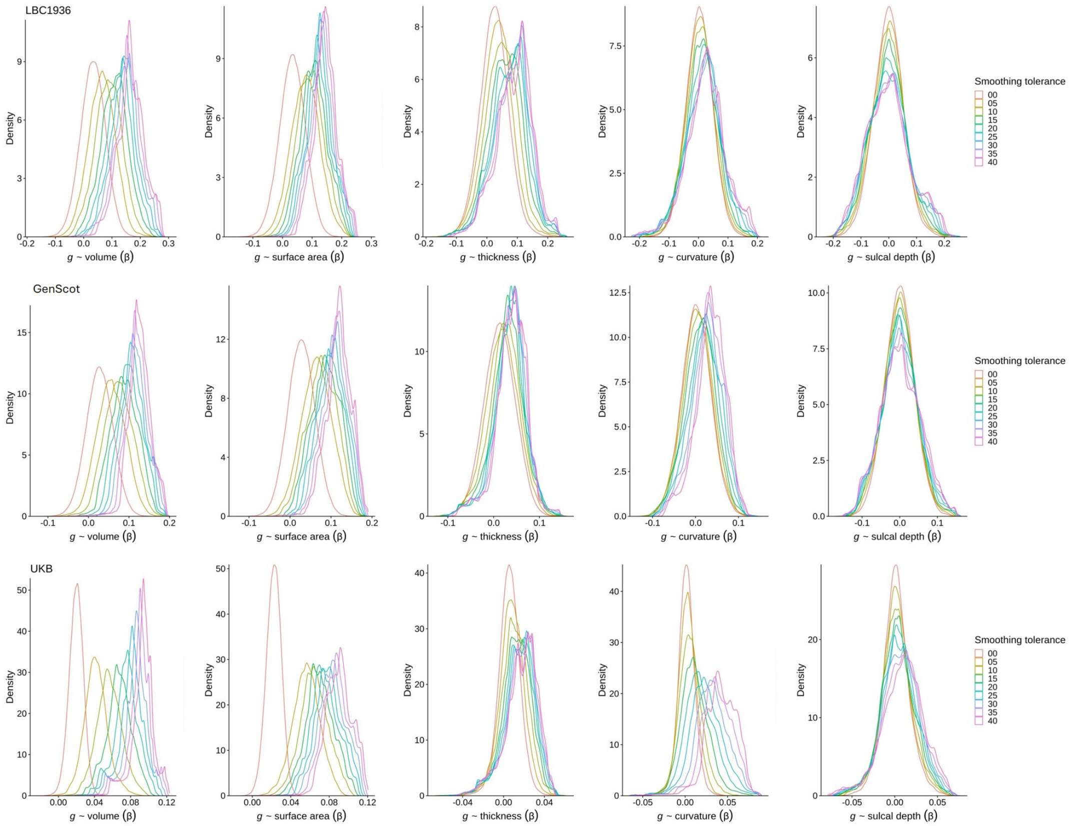


*Figure S5* Density plots showing *g*-associations for each cohort for each of the 9 smoothing tolerances (0, 5, 10, 15, 20, 25, 30, 35 and 40 mm FWHM).

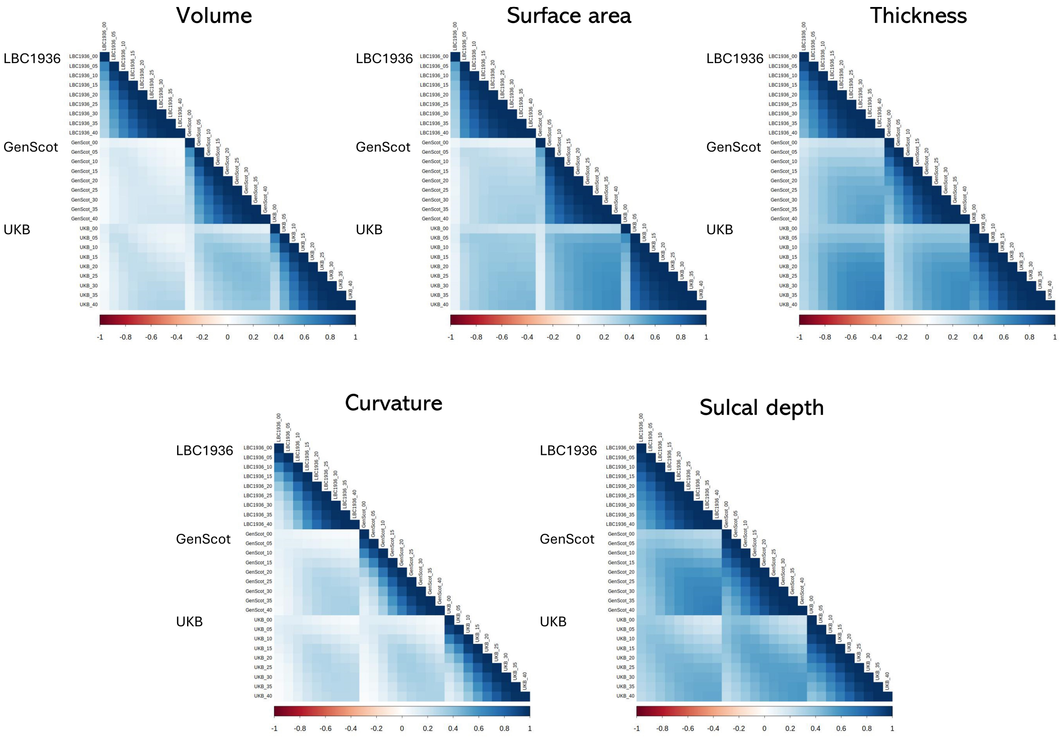


*Figure S6* Within and between cohort spatial correlations of vertex-wise *g*-associations for the 5 cortical morphometry measures at each of the 9 smoothing tolerances (0, 5, 10, 15, 20, 25, 30, 35, 40 mm FWHM). Between-cohort spatial correlations increase with increasing smoothing tolerance, as data converges towards the total surface estimates, while within-cohort correlations decrease with increasing smoothing tolerances.

#### Vertex-wise meta-analysis results

#### g associations

***g* ~ volume**

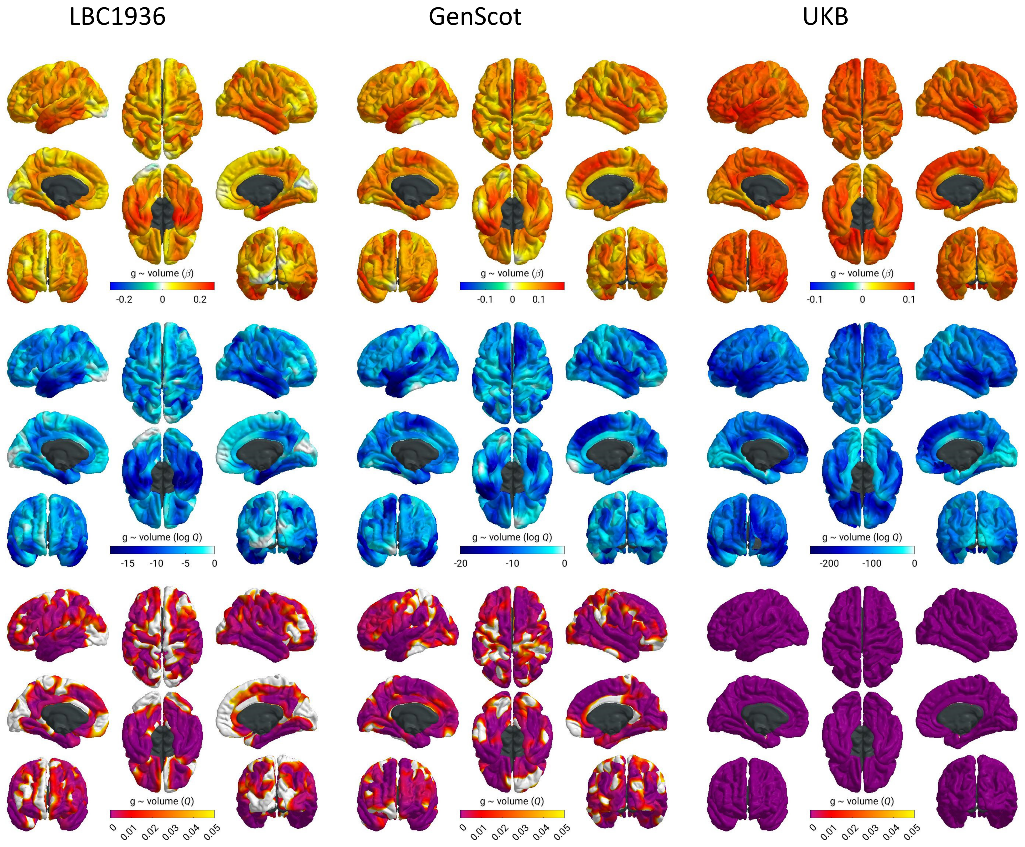


*Figure S7* *g*-volume associations for each cohort at 20 FWHM (top = β, middle = log *FDR Q*, bottom = *FDR Q*). The beta and log *FDR Q* scales are set at the maximum values for the relevant cohort across all measures.

***g* ~ surface area**

LBC1936 GenScot UKB

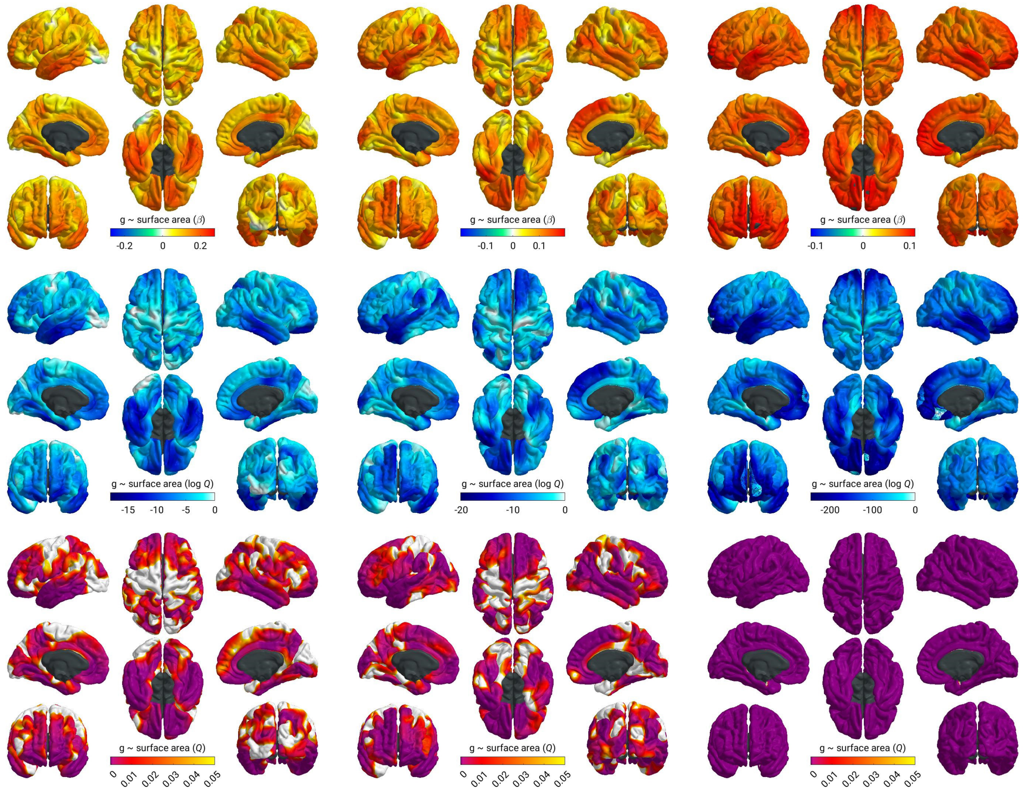


*Figure S8 g*-surface area associations for each cohort at 20 FWHM (top = β, middle = log *FDR Q*, bottom = *FDR Q*). The beta and log *FDR Q* scales are set at the maximum values for the relevant cohort across all measures.

***g* ~ thickness**

LBC1936 GenScot UKB

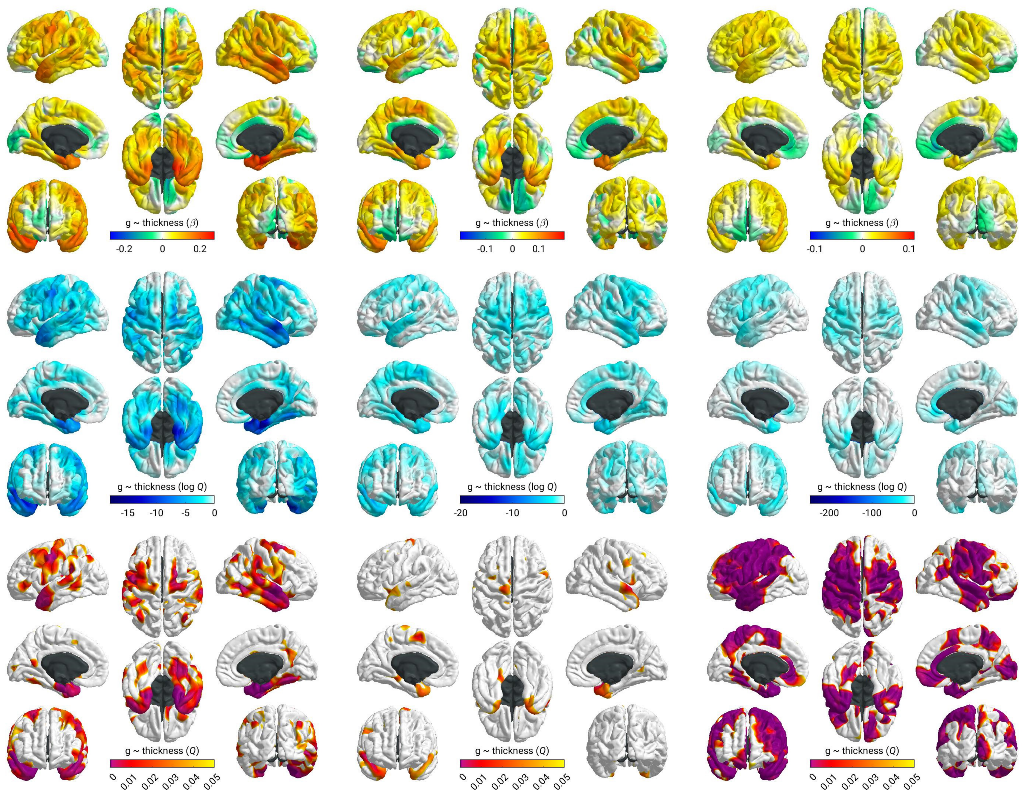


*Figure S9 g*-thickness associations for each cohort at 20 FWHM (top = β, middle = log *FDR Q*, bottom = *FDR Q*). The beta and log *FDR Q* scales are set at the maximum values for the relevant cohort across all measures.

#### *g* ~ curvature

LBC1936 GenScot UKB

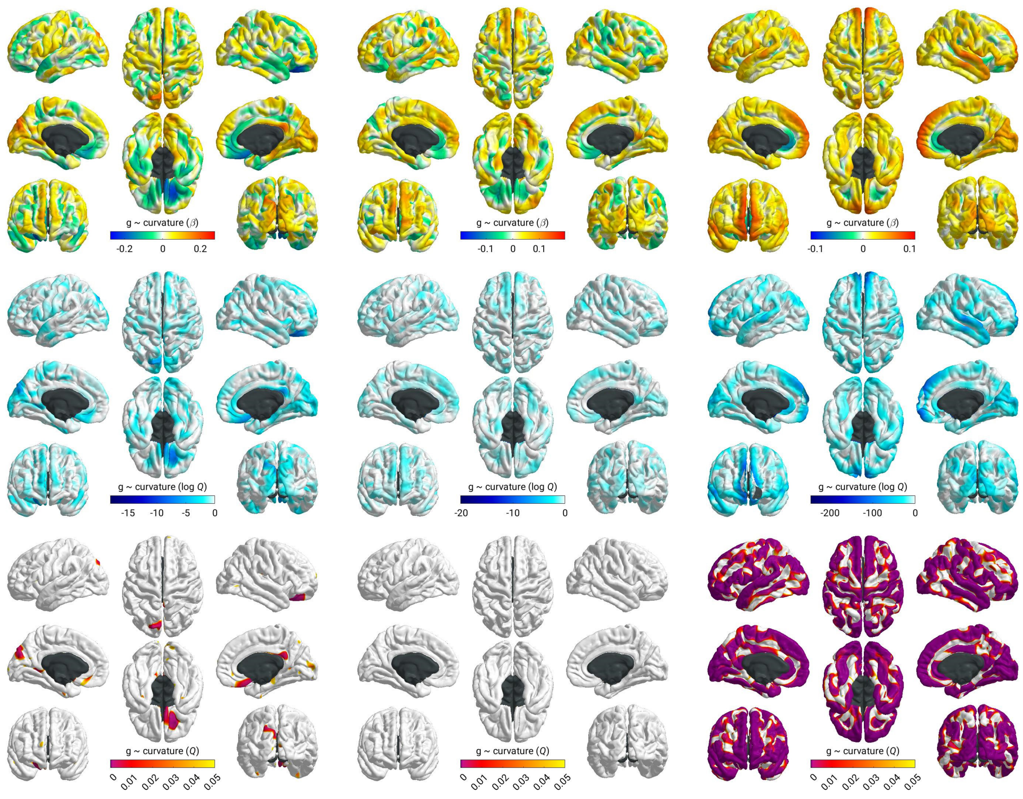


*Figure S10 g*-curvature associations for each cohort at 20 FWHM (top = β, middle = log *FDR Q*, bottom = *FDR Q*). The beta and log *FDR Q* scales are set at the maximum values for the relevant cohort across all measures.

#### *g* ~ sulcal depth

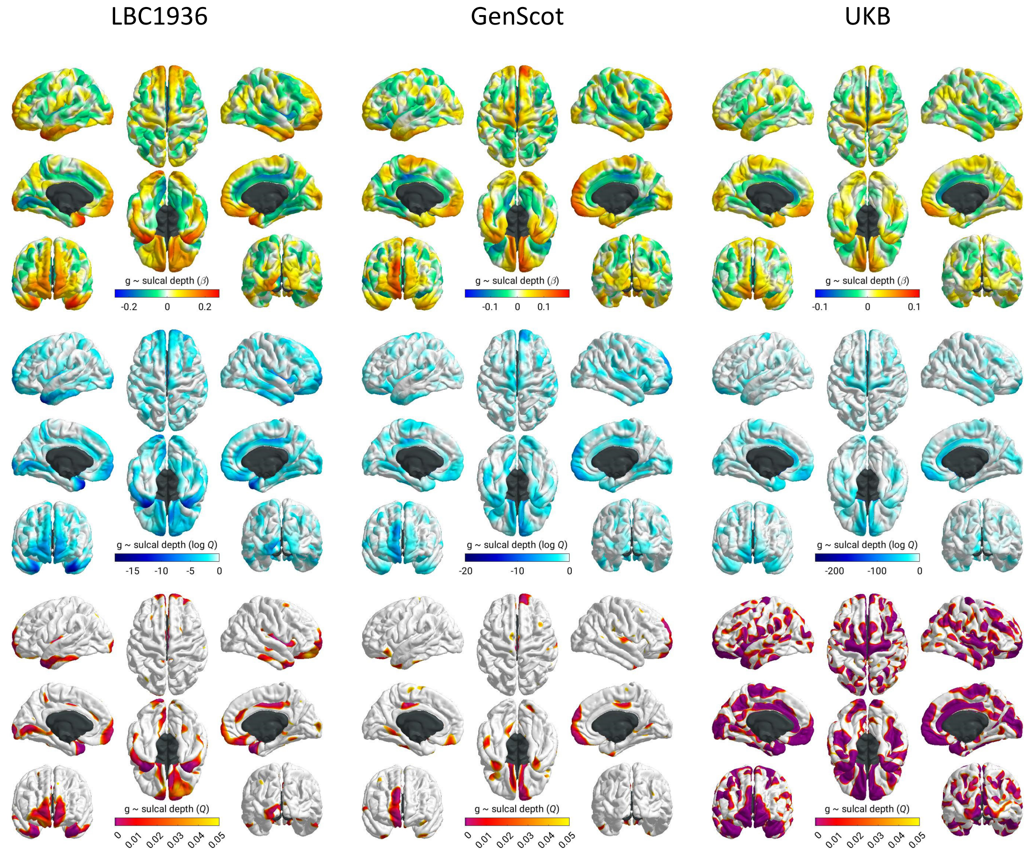


*Figure S11 g*-sulcal depth associations for each cohort at 20 FWHM (top = β, middle = log *FDR Q*, bottom = *FDR Q*). The beta and log *FDR Q* scales are set at the maximum values for the relevant cohort across all measures.

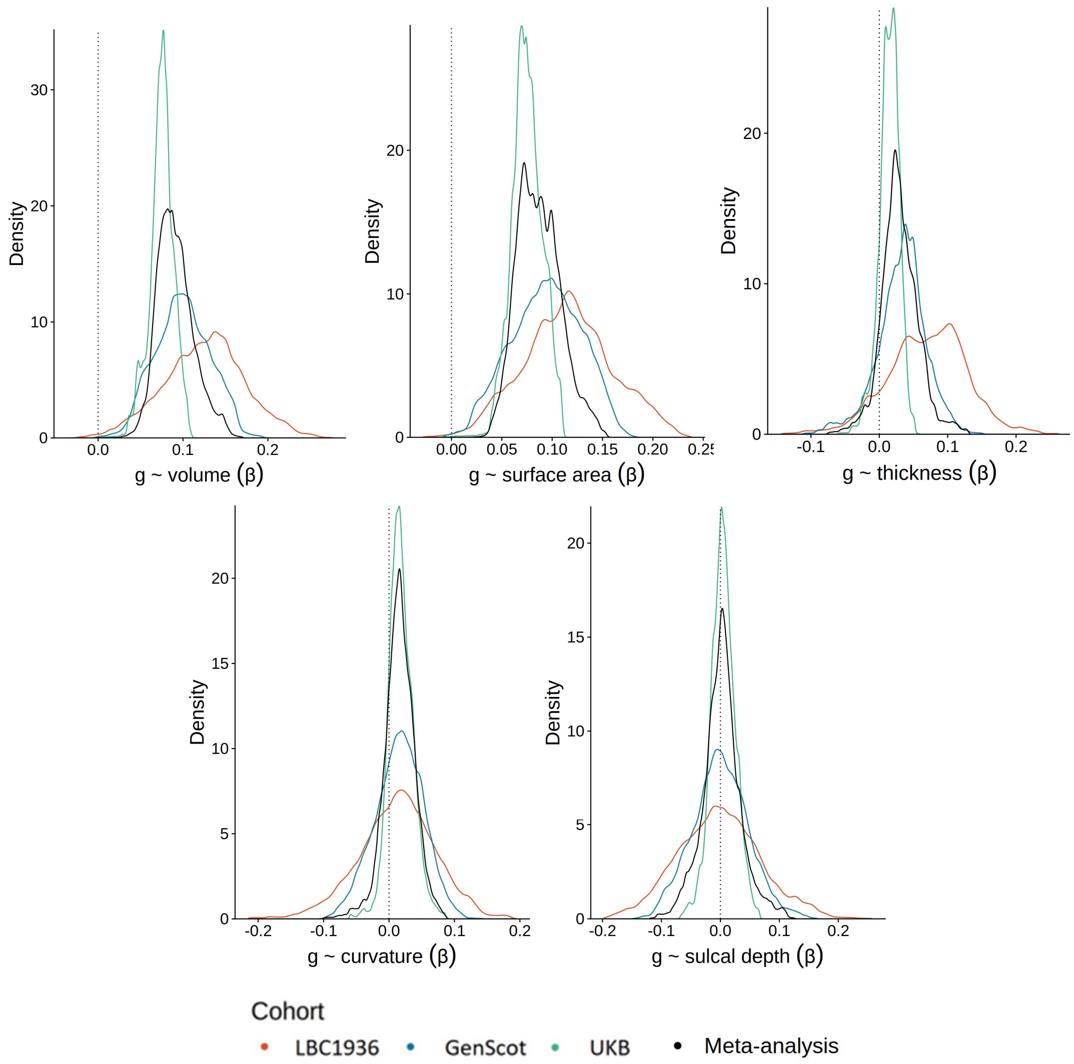


*Figure S12* Density distributions of *g*-association β values for the three cohorts and the meta-analysed *g*-associations for the 5 vertex-wise measures of morphometry. The vertical dotted line marks β = 0.

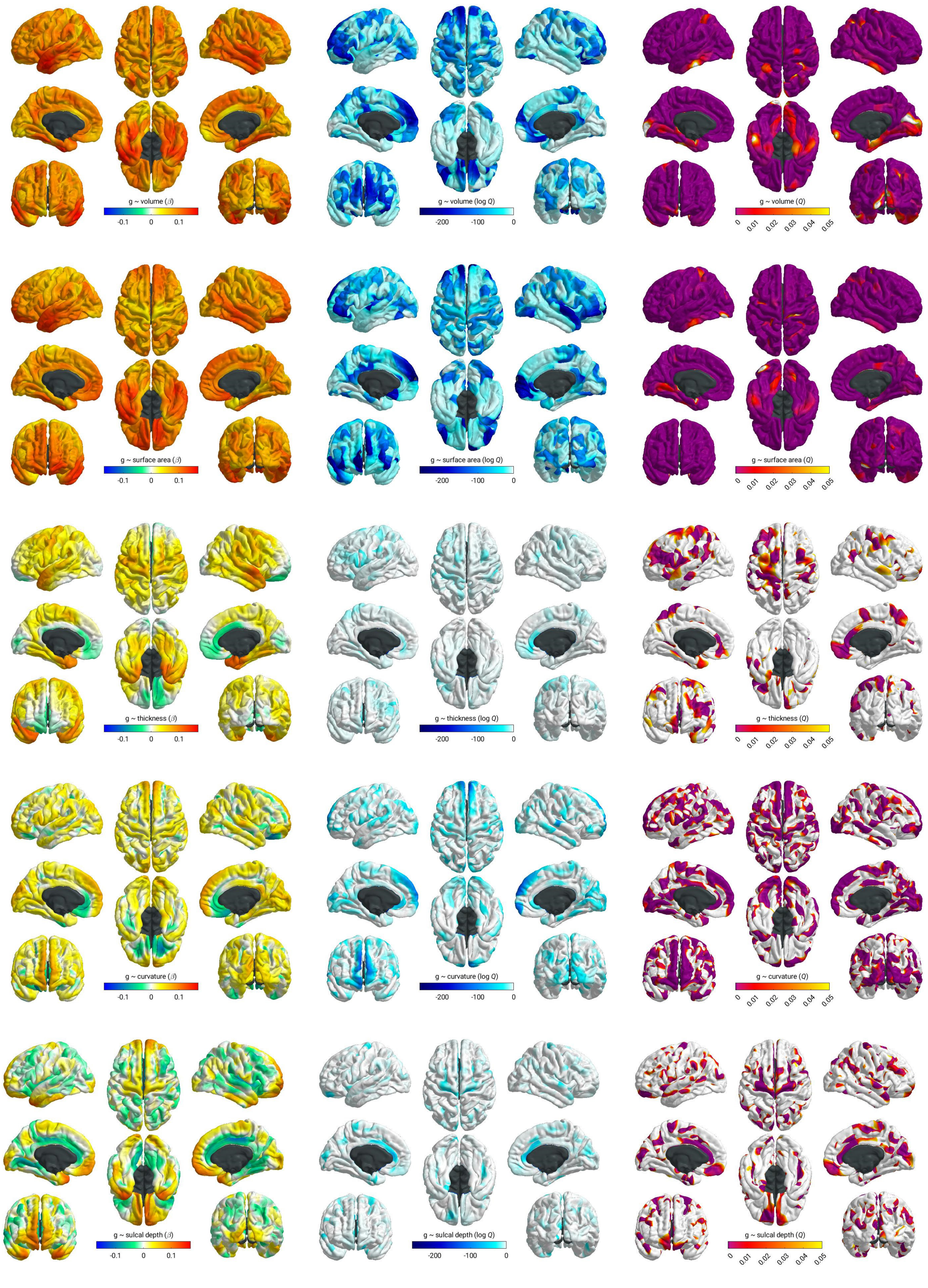


*Figure S13* Meta-analysed β, log *FDR Q* and *FDR Q* values for *g* ~ morphometry associations for the 5 vertex-wise measures (from top to bottom: volume, surface area, thickness, curvature and sulcal depth). The scale limits for the β maps set at 0.17+/-, which is the maximum absolute value for any measure. The scale limits for log *Q* maps is set at the minimum value for any measure (which is -263.24, or *FDR Q* = $4.75\text{x}10^{-115}$).

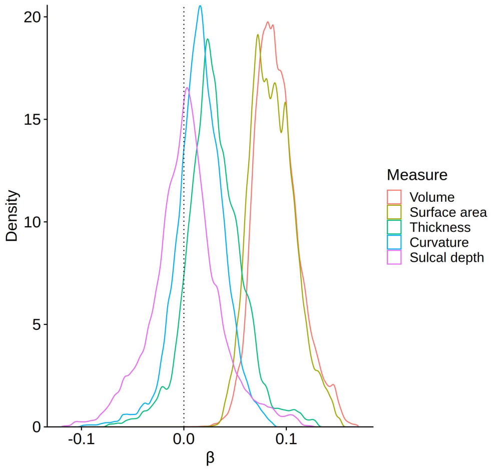


*Figure S14* Density distributions for the meta-analysed *g* ~ morphometry associations for the 5 measures of morphometry (volume, surface area, thickness curvature and sulcal depth). The vertical dotted line marks β = 0.

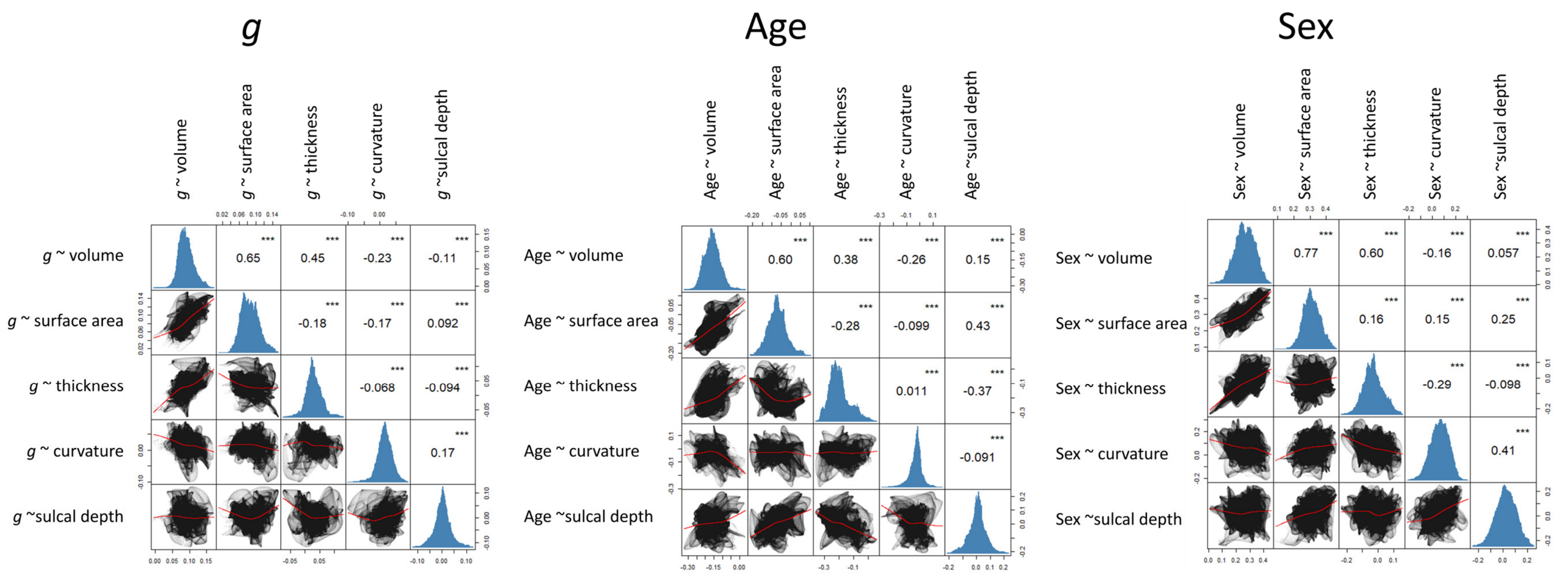


*Figure S15* Spatial correlation plots of the meta-analysed β values for vertex-wise associations of 5 measures of morphometry (volume, surface area, thickness, curvature and sulcal depth) and (left) *g*, (middle) age and (right) sex. These are linked to Table 3 in the main text.

#### Metabolism

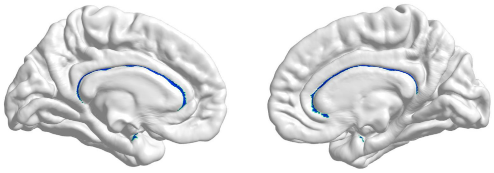

*Figure S16* The metabolism data was registered from fsLR 164k to fsaverage 164k, and there was a slight difference in the cortical masks, in that there were 2153 additional vertices included in the mask compared to the fsaverage mask. These vertices are shown in blue in this figure. They were not included in any of the correlations that include metabolism.

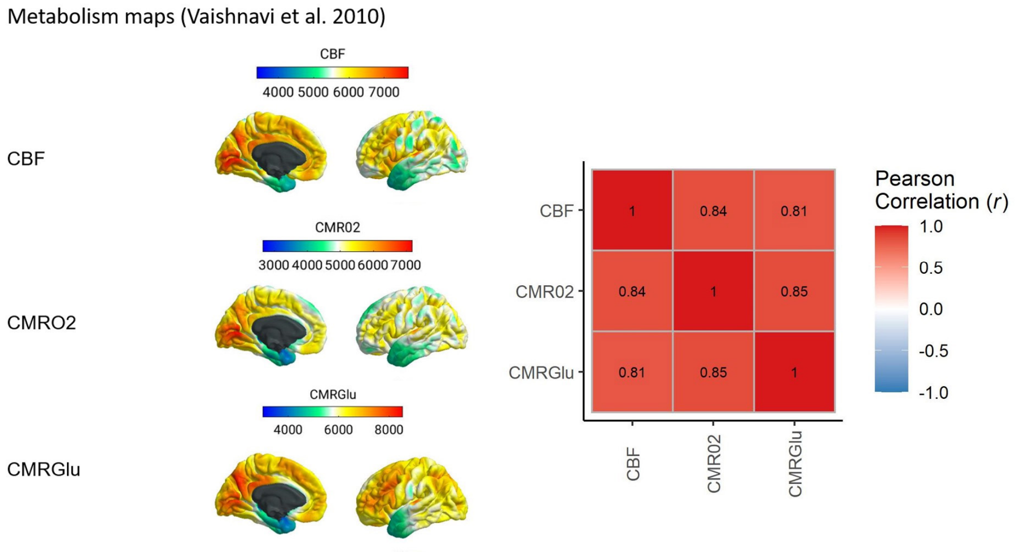


*Figure S17* Left: Metabolism data mapped to the cortex. Right: Spatial correlation plot showing the high correlations between three measures of cortical metabolism, which justifies a principal component analysis to create one measure of cortical metabolism.

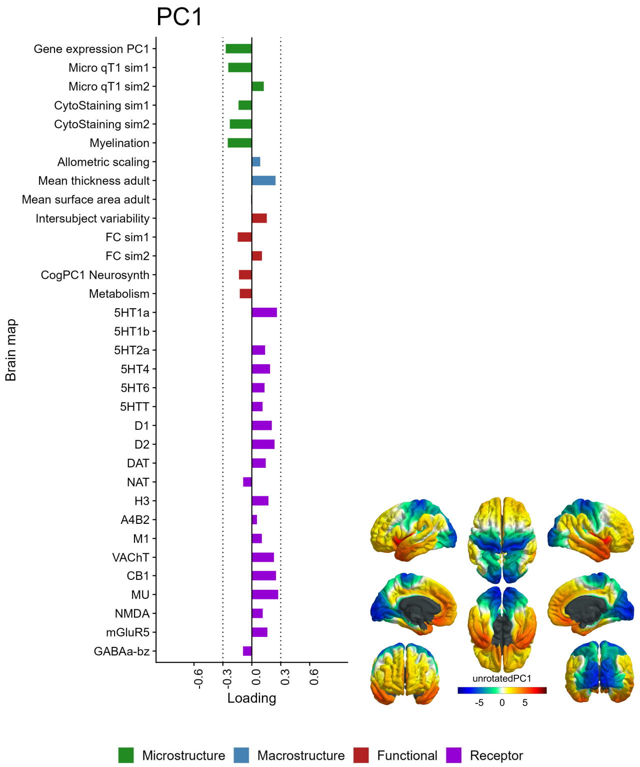


*Figure S18* Unrotated PC1 loadings. Note, the coefficient of factor congruence between the unrotated PC1 and PC1 after varimax rotation with 4 components was 0.9.

#### Global and subcortical brain structure descriptives

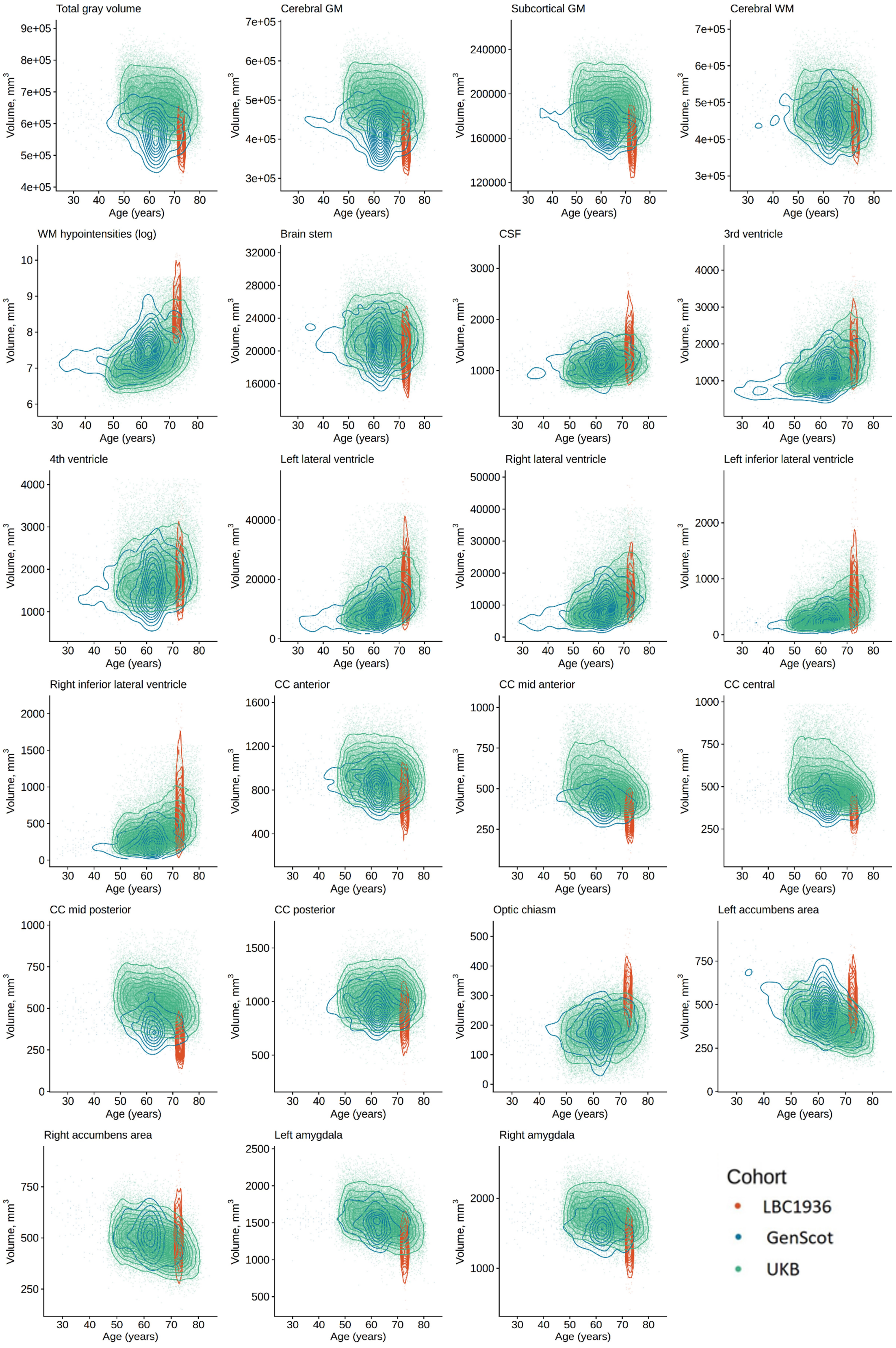

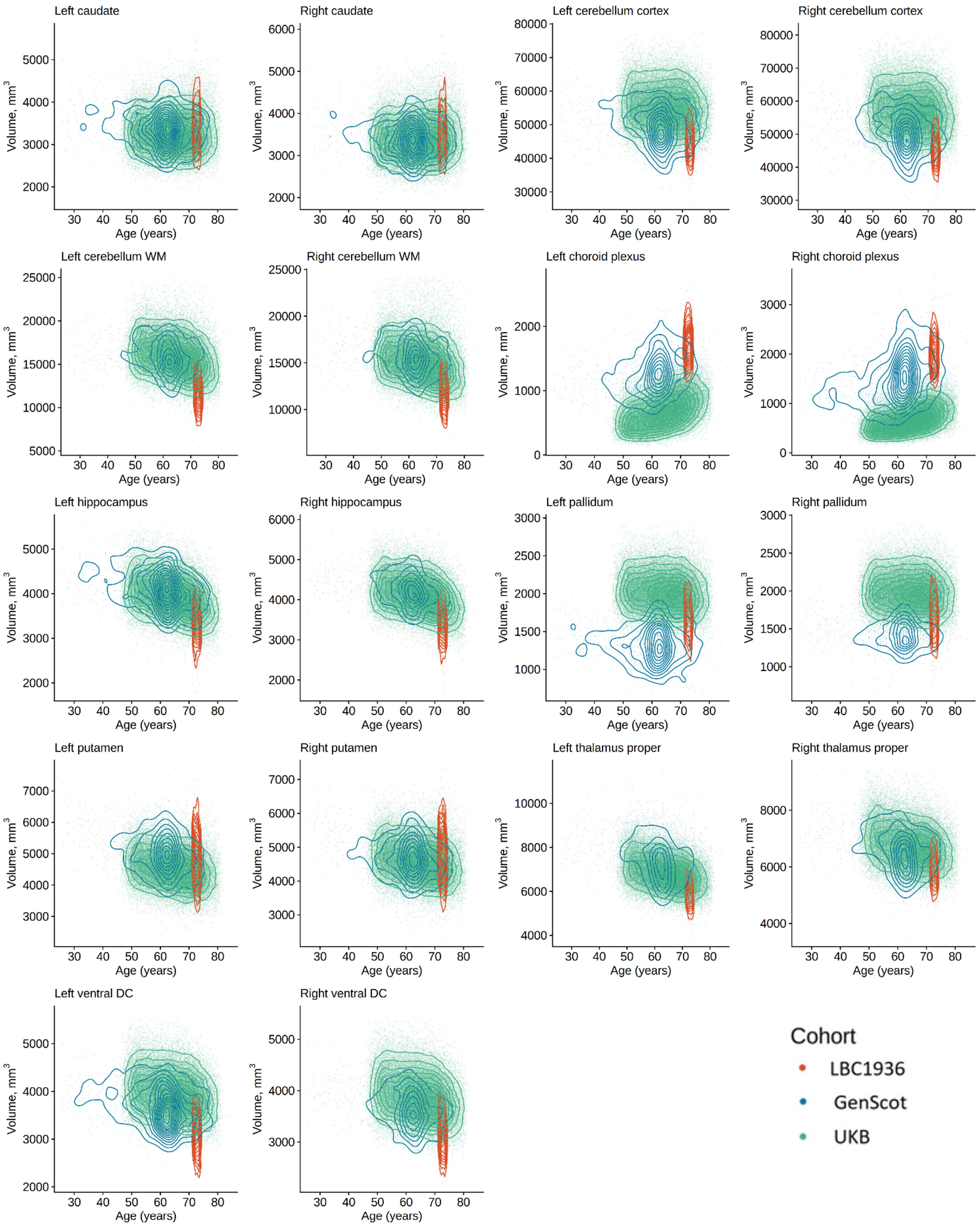


*Figure S19* Raw data plots of the volume of each global and subcortical brain structure, coloured by cohort.

#### Neurobiological profiles

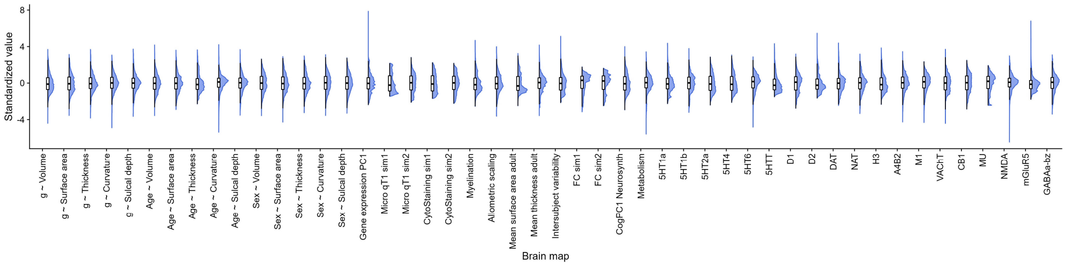


*Figure S20* Density distributions of all the vertex-wise maps (g-associations, age-associations, sex-associations and 33 neurobiological profiles)

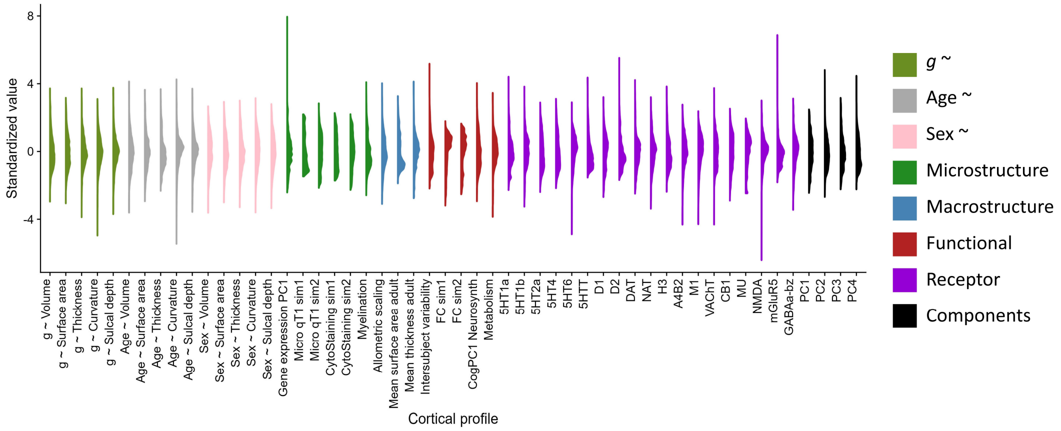


*Figure S21* Distributions of standardized values for all cortical profiles, including PCs, coloured by profile type.

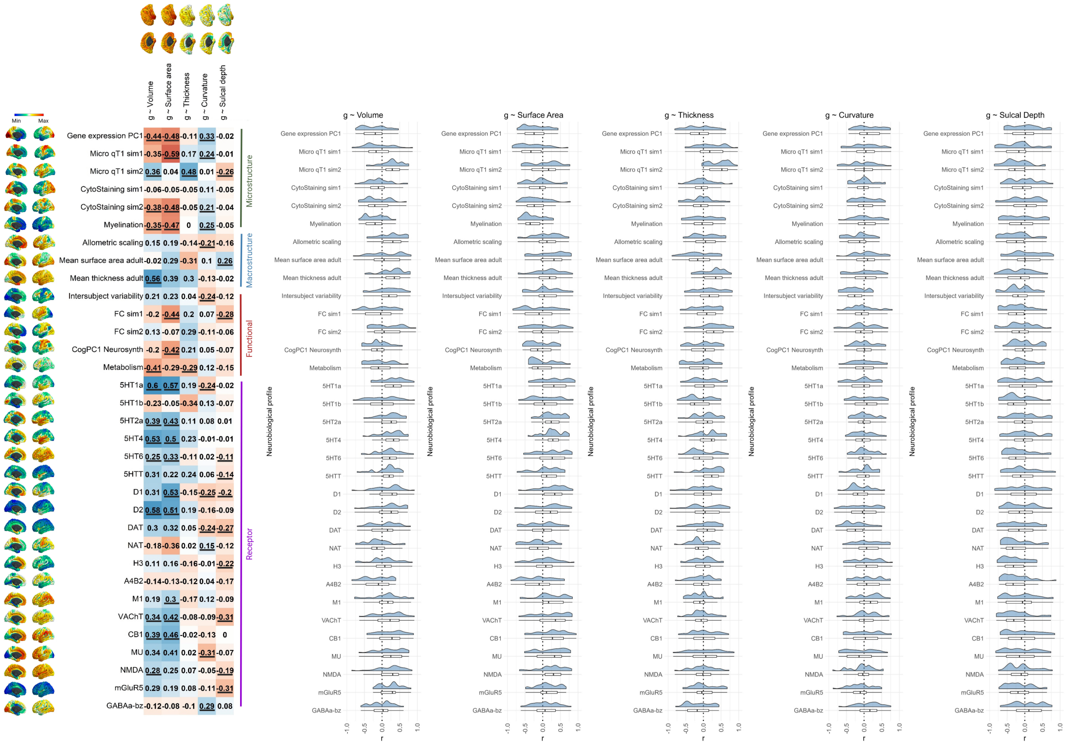

Figure S22 Extended version of main text Figure 5. A) Correlation plots showing spatial correlations between g-vertex-wise morphometry associations and various multiscale cortical profiles Those with *p_spin* values < .05 are underlined. B) Summary data of regional correlations for the cortex-level correlations in A).

### Regional correlations

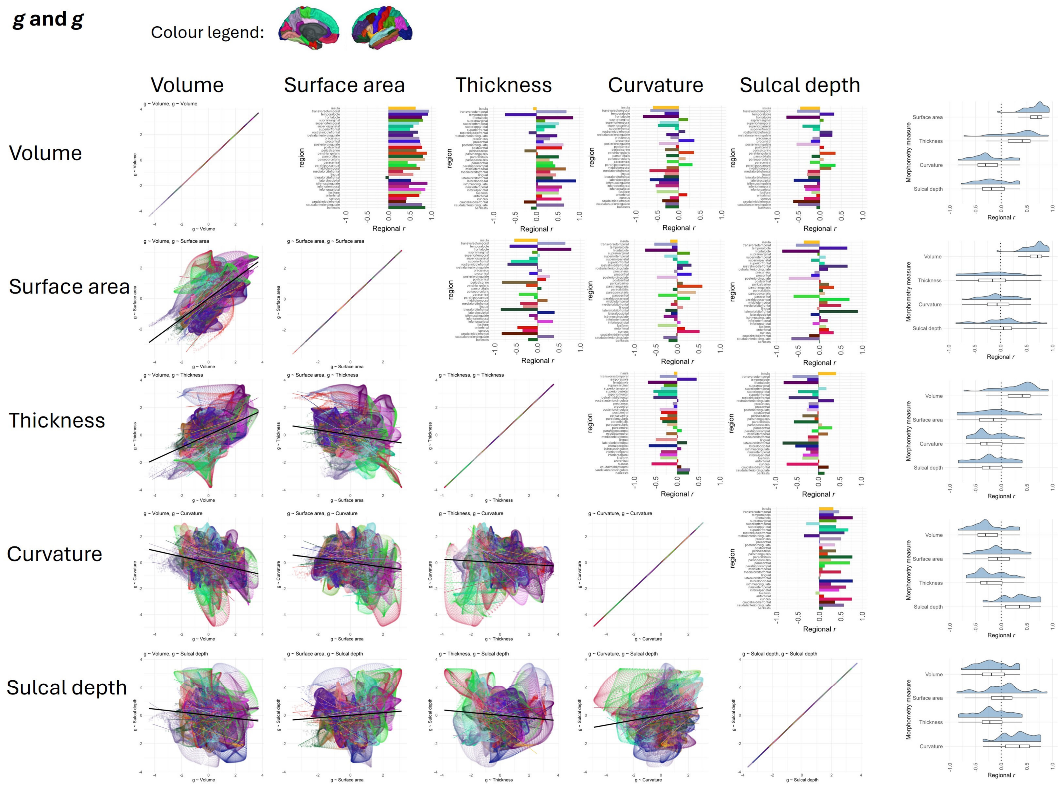


*Figure* S23 Within-region vertex-wise spatial correlations for **different morphometry measures with *g***. A) scatterplots showing each correlation, coloured by Desikan-Killiany region, B) summary of regional correlations, C) bar graph showing each regional correlation. The data underpinning these figures are in the Supplementary Tabular Data File.

#### Age and age

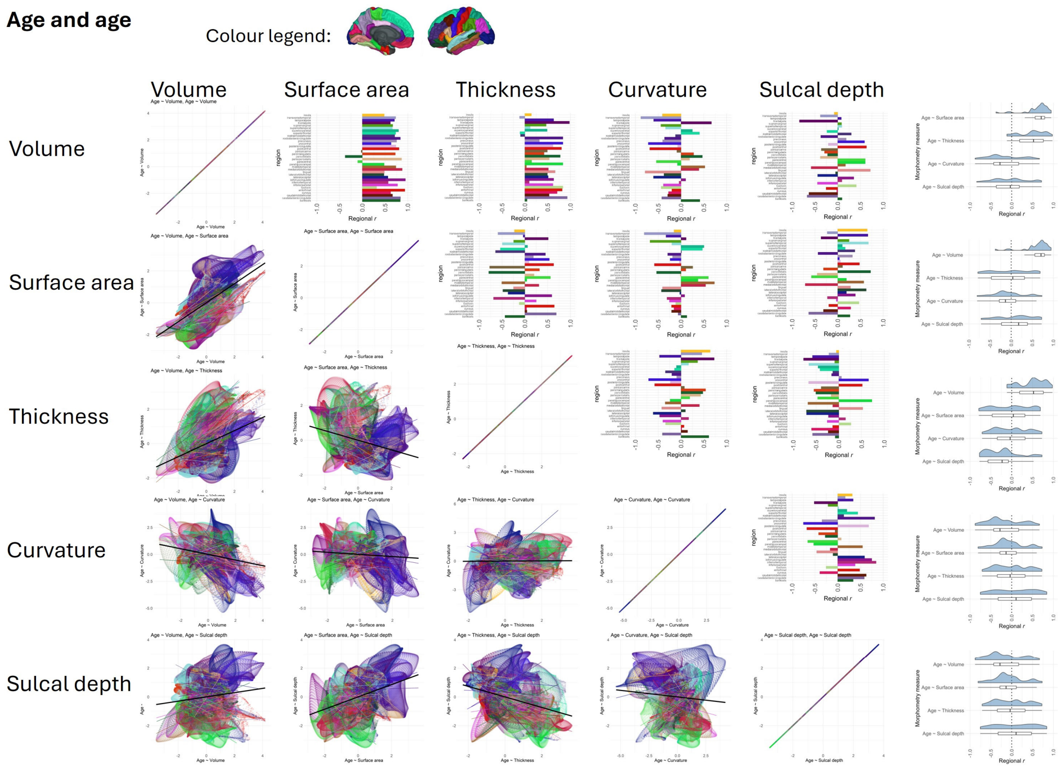


*Figure* S24 Within-region vertex-wise spatial correlations for **different morphometry measures with age.** A) scatterplots showing each correlation, coloured by Desikan-Killiany region, B) summary of regional correlations, C) bar graph showing each regional correlation. The data underpinning these figures are in the Supplementary Tabular Data File.

**Sex and sex**

Colour legend: 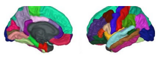

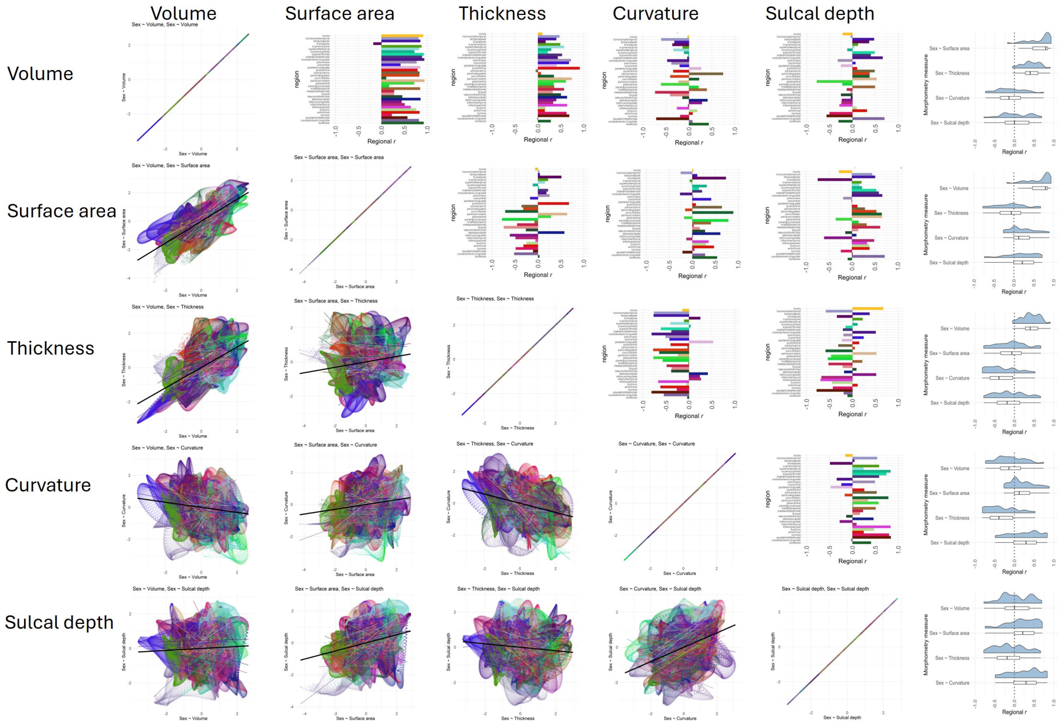


*Figure* S25 Within-region vertex-wise spatial correlations for **different morphometry measures with sex**. A) scatterplots showing each correlation, coloured by Desikan-Killiany region, B) summary of regional correlations, C) bar graph showing each regional correlation. The data underpinning these figures are in the Supplementary Tabular Data File.

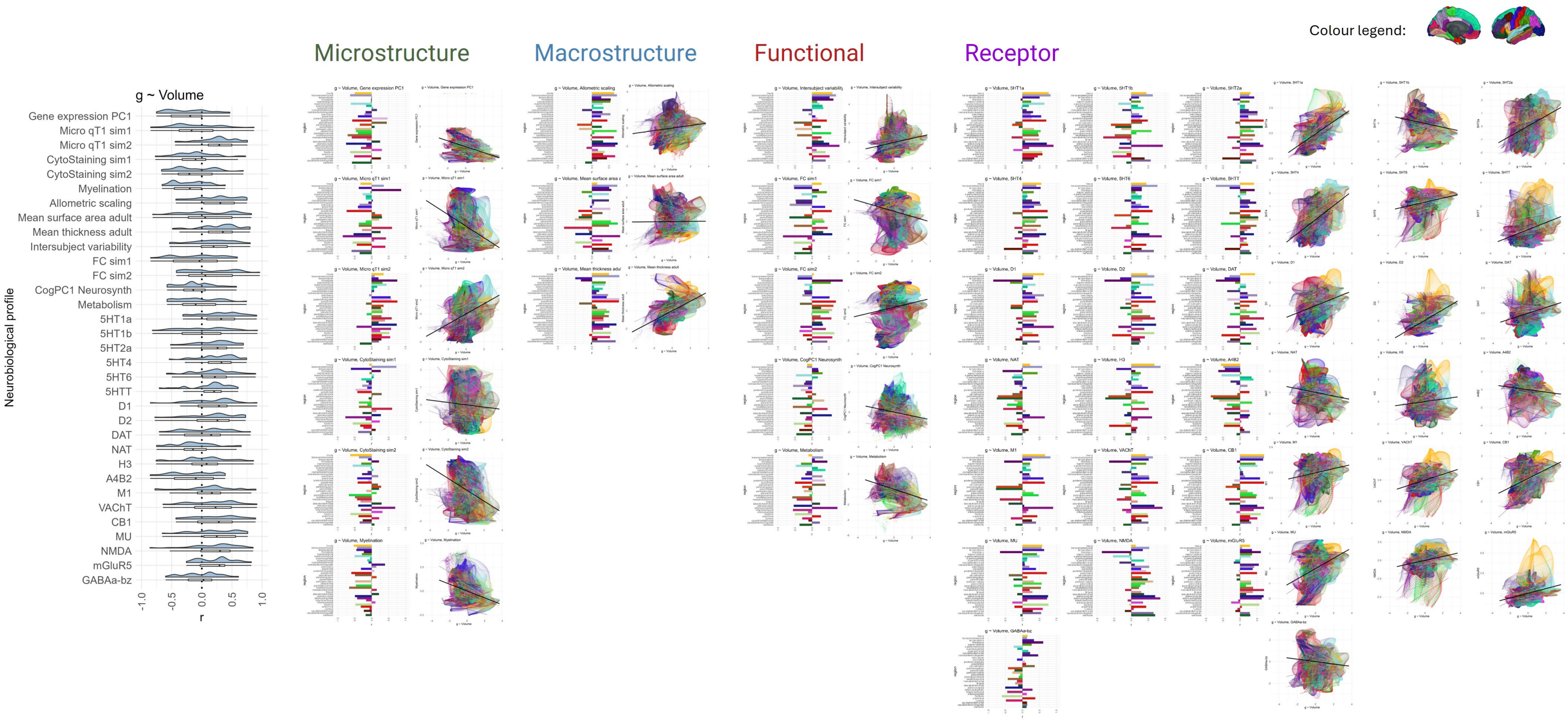


*Figure* S26 Within-region vertex-wise spatial correlations for ***g*-volume and neurobiological profiles** correlations. A) summary of regional correlations, B) bar graph showing each regional correlation and scatterplots showing each correlation, coloured by Desikan-Killiany region. The data underpinning these figures are in the Supplementary Tabular Data File.

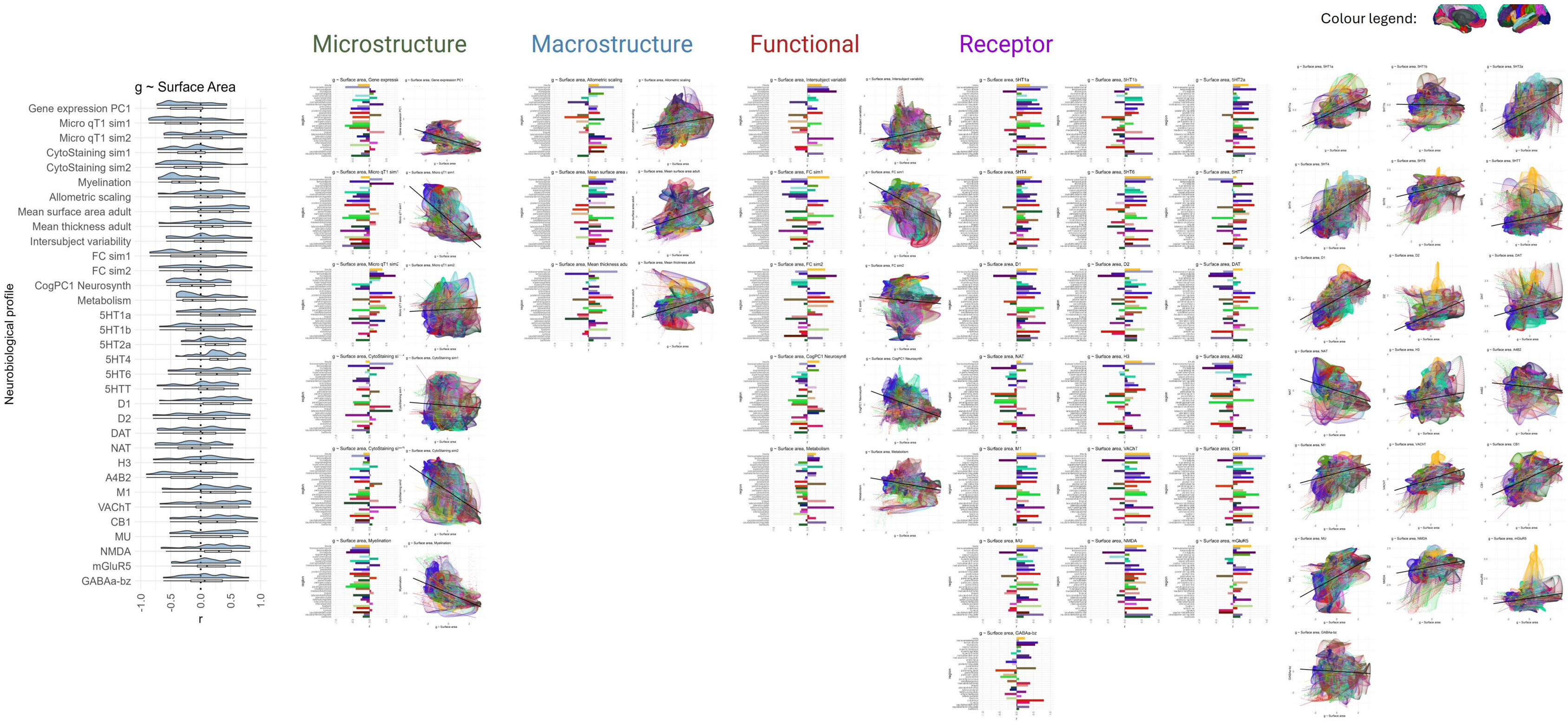


*Figure* S27 Within-region vertex-wise spatial correlations for ***g*-surface area and neurobiological profiles** correlations. A) summary of regional correlations, B) bar graph showing each regional correlation and scatterplots showing each correlation, coloured by Desikan-Killiany region. The data underpinning these figures are in the Supplementary Tabular Data File.

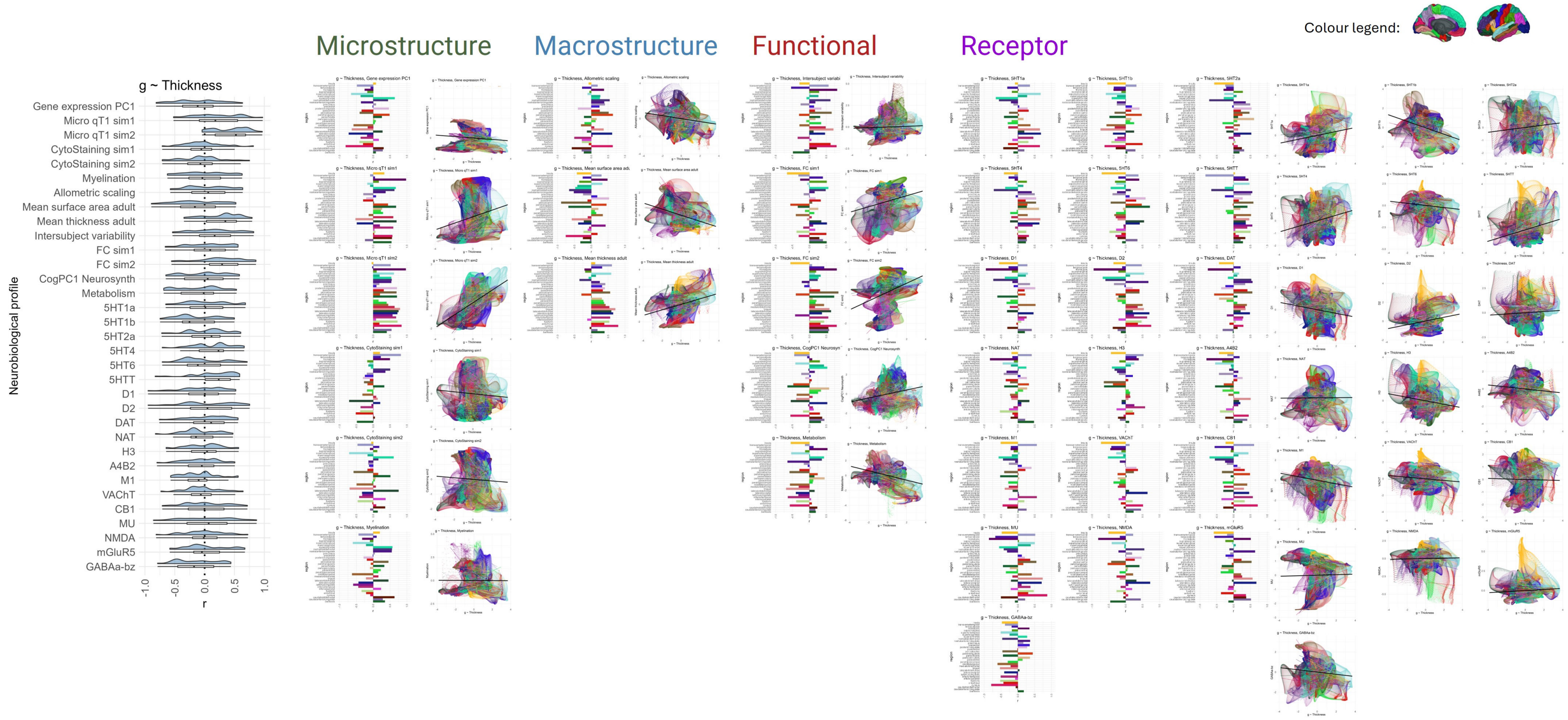


*Figure* S28 Within-region vertex-wise spatial correlations for ***g*-thickness and neurobiological profiles** correlations. A) summary of regional correlations, B) bar graph showing each regional correlation and scatterplots showing each correlation, coloured by Desikan-Killiany region. The data underpinning these figures are in the Supplementary Tabular Data File.

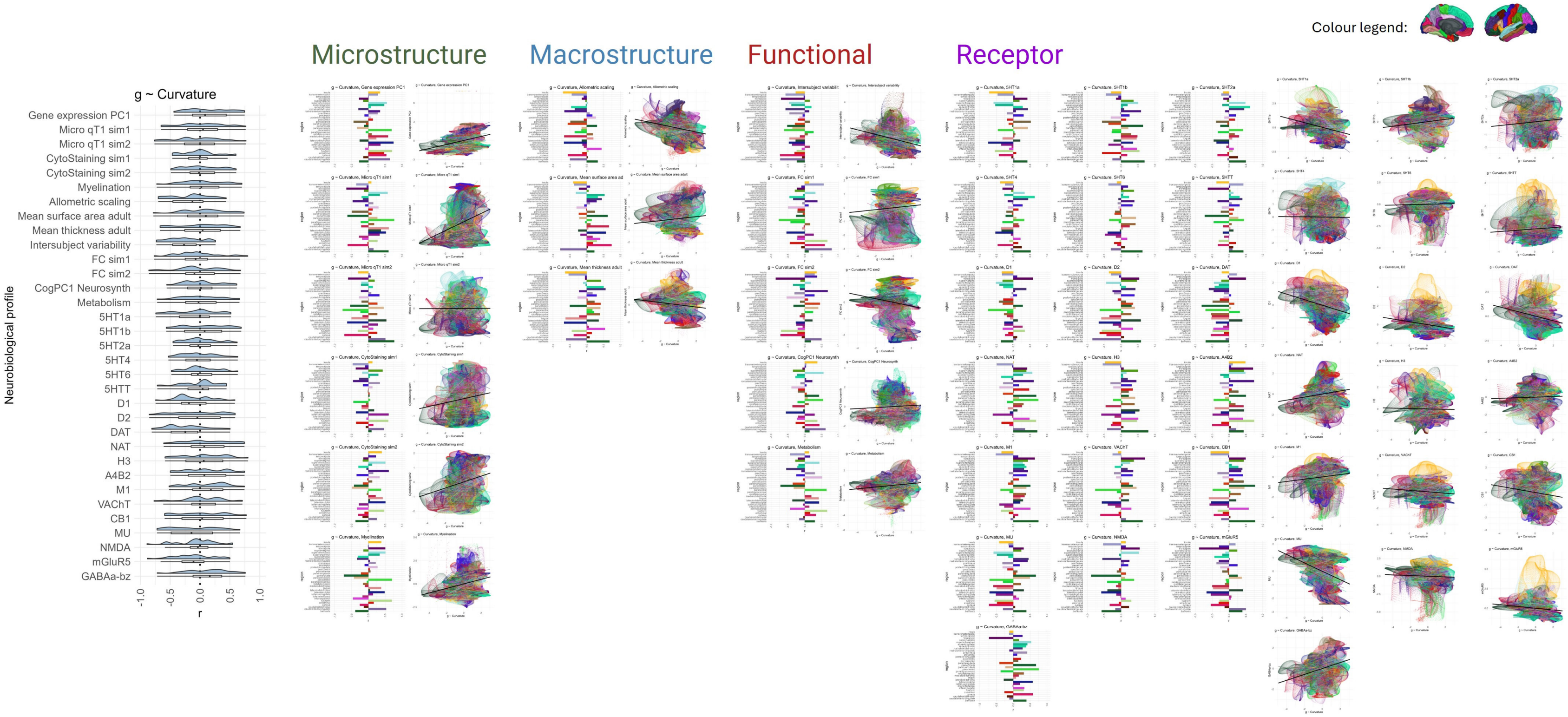


*Figure* S29 Within-region vertex-wise spatial correlations for ***g*-curvature and neurobiological profiles** correlations. A) summary of regional correlations, B) bar graph showing each regional correlation and scatterplots showing each correlation, coloured by Desikan-Killiany region. The data underpinning these figures are in the Supplementary Tabular Data File.

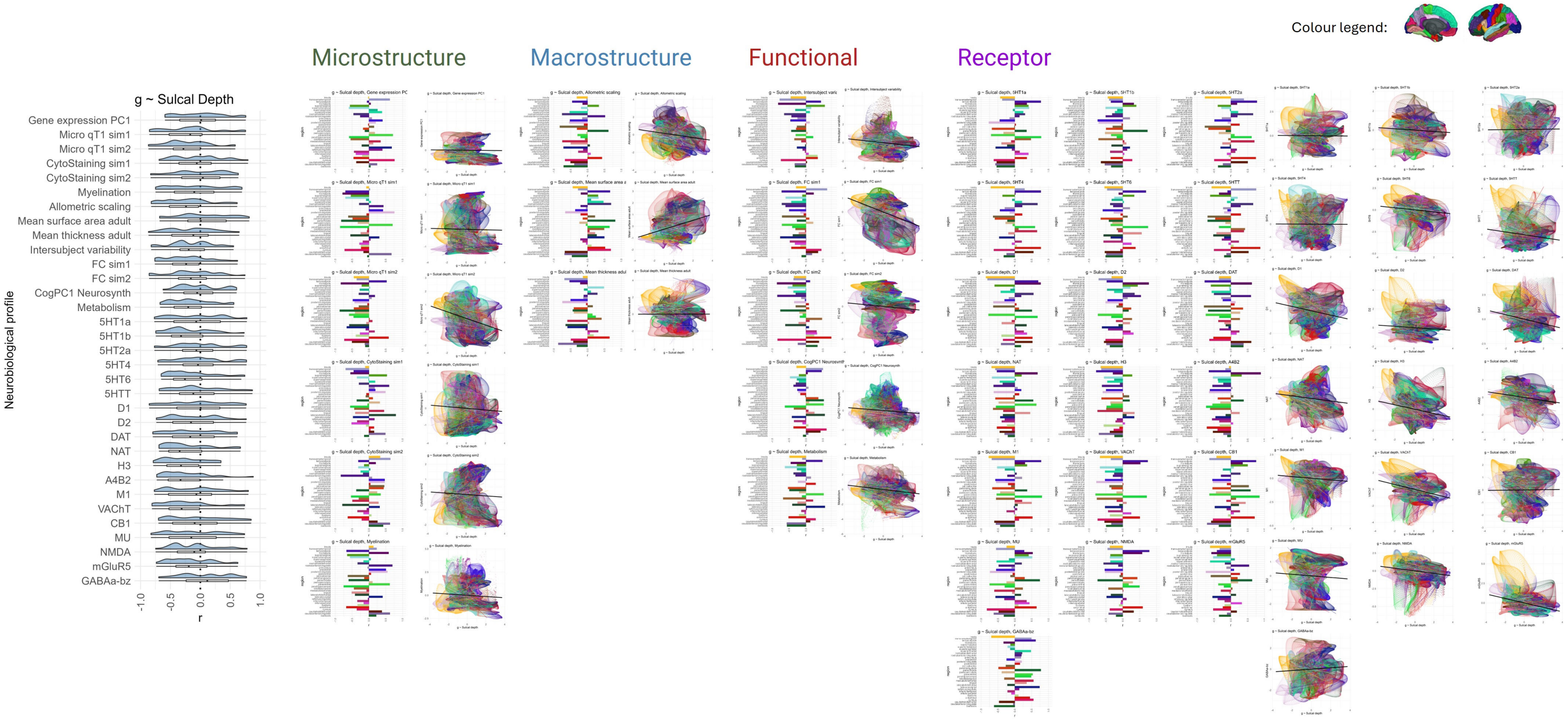


*Figure* S30 Within-region vertex-wise spatial correlations **for *g*-sulcal depth and neurobiological profiles** correlations. A) summary of regional correlations, B) bar graph showing each regional correlation and scatterplots showing each correlation, coloured by Desikan-Killiany region. The data underpinning these figures are in the Supplementary Tabular Data File.

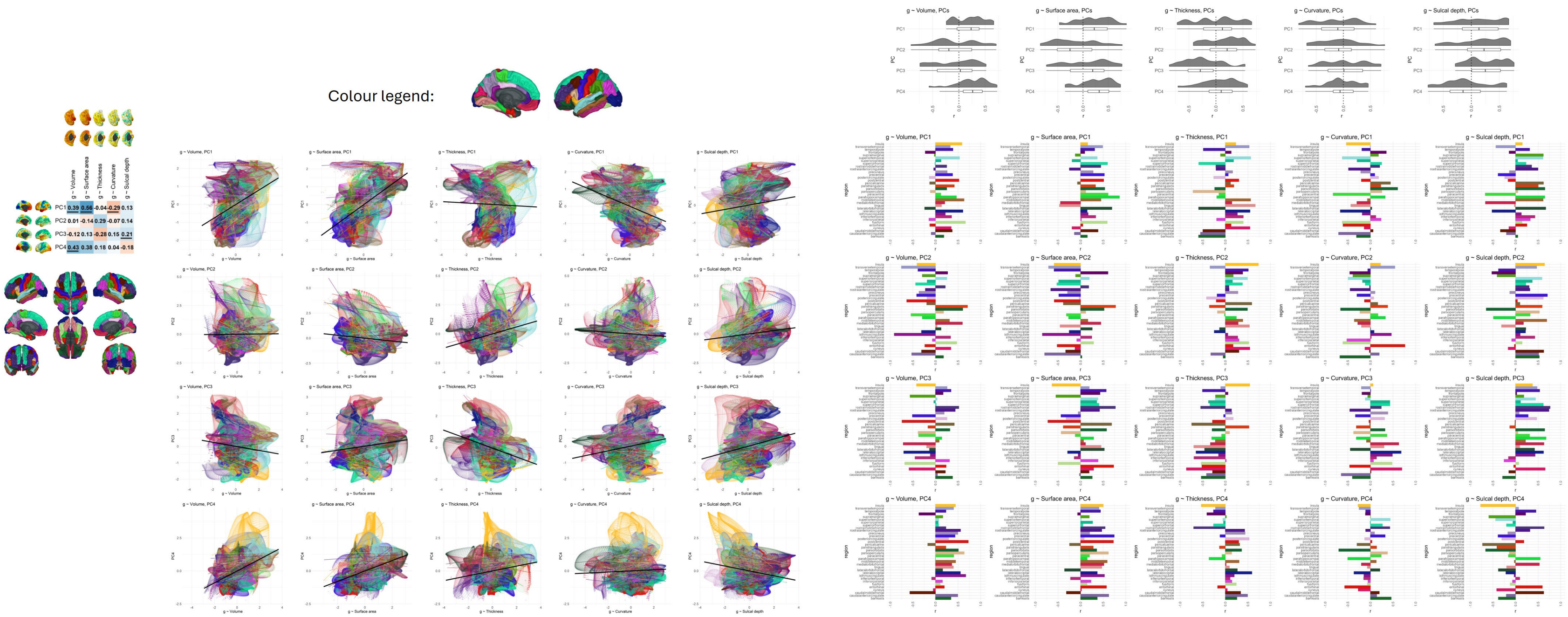


*Figure* S31 Regional spatial correlations for ***g* and PC** correlations. A) cortex-wide correlations, B) scatterplots showing each correlation, coloured by Desikan-Killiany region, C) summary of regional correlations, D) bar graph showing each regional correlation. The data underpinning these figures are in the Supplementary Tabular Data File.

#### Supplementary tables

Table S1 UK Biobank exclusions and database codes.

| Exclusion criteria | Root code | Specific code |
|---|---|---|
| Dementia | 20002 | 1263 |
| Parkinson’s disease | 20002 | 1262 |
| Stroke | 20002 | 1081 |
| Other chronic neurological problems | 20002 | 1258 |
| Other demyelinating diseases | 20002 | 1397 |
| Multiple sclerosis | 20002 | 1261 |
| Guillain–Barré syndrome | 20002 | 1256 |
| Brain cancer | 20001 | 1032 |
| Brain haemorrhage | 20002 | 1491 |
| Brain abscess | 20002 | 1245 |
| Brain aneurysm | 20002 | 1425 |
| Cerebral palsy | 20002 | 1433 |
| Encephalitis | 20002 | 1246 |
| Epilepsy | 20002 | 1264 |
| Head injury | 20002 | 1266 |
| Infection of nervous system | 20002 | 1244 |
| Ischaemic stroke | 20002 | 1583 |
| Meningeal cancer | 20001 | 1031 |
| Meningioma (benign) | 20002 | 1659 |
| Meningitis | 20002 | 1247 |
| Motor neurone disease | 20002 | 1259 |
| Neurological trauma | 20002 | 1240 |
| Spina bifida | 20002 | 1524 |
| Subdural haematoma | 20002 | 1083 |
| Subarachnoid haemorrhage | 20002 | 1086 |
| Transient ischaemic attack | 20002 | 1082 |

Table S2 Brief descriptions of UKB cognitive tests and index codes.

| Cognitive Test | Brief description | Code |
|---|---|---|
| Reaction time (s) | Time taken to respond in snap-type computer game | 20023 |
| Number span | Number of rounds completed (the maximum length of number string recalled) | 4282 |
| Verbal and numerical reasoning (called “fluid intelligence” in the UKB database) | Number of 13 verbal and numerical logic questions correct | 20016 |
| Trail making B (s) [1] | Time taken to complete trail B | 6350 |
| Matrix pattern (log) [2] | Number of puzzles solved | 6373 |
| Tower task | Number of puzzles solved | 21003 |
| Digit-symbol substitution [3] | Number of digit-symbol pairs matched | 23324 |

| | | |
|---|---|---|
| Pairs matching [4] | Number of incorrect matches in a 6-pair classic pairs game | 399 |
| Prospective memory | Did the participant remember to ignore the instruction on their first attempt? | 20018 |
| Paired associates | Number of novel word pairs matched in recall | 20197 |

Table S3 Brief descriptions of GenScot cognitive tests and index codes.

| Cognitive Test | Brief description | Code |
|---|---|---|
| Matrix reasoning [2] | Number of puzzles correct | mrtotc |
| Verbal fluency [5] | Number of words recalled beginning with C, F and L in 3x1 minute | vftot |
| Mill Hill vocabulary [6] | Number of word meanings explained correctly | mhv |
| Digit symbol substitution [7] | Number of digit-symbol pairs matched | digsym |
| Logical memory [7] | Story recall score (total from immediate and delayed tests) | mema + medela |

Table S4 Brief descriptions of LBC1936 cognitive tests and index codes.

| Cognitive test | Brief description | Code |
|---|---|---|
| Matrix reasoning [8] | Number of puzzles correct | matreas_w2 |
| Block design [8] | Number of puzzles correct | blkdes_w2 |
| Spatial span [7] | Number of block sequences correct | spantot_w2 |
| National adult reading test (NART) [9] | Number of words pronounced correctly | nart_w2 |
| Weschler Test of Adult reading (WTAR) [10] | Number of words pronounced correctly | wtar_w2 |
| Verbal fluency [11] | Number of words recalled beginning with C, F and L in 3x1 minute | vftot_w2 |
| Verbal paired associates [7] | Number of novel word pairs matched in recall (total from immediate and delayed tests) | vpatotal_w2 |
| Logical memory [7] | Number of story details recalled (out of a total possible of 25) Total from immediate and delayed tests | lmtotal_w2 |
| Digit span backwards [8] | Max number of a string of numbers recalled in reverse | digback_w2 |
| Symbol search | Number of symbols correctly detected | |
| Digit-symbol substitution [12] | Number of digit-symbol pairs matched | digsym_w2 |
| Inspection time [13] | Number of correct responses – is the left or right line longer? | Ittotal_w2 |

| | | |
|---|---|---|
| Four-choice reaction time (s) 14 | Time taken to press the indicated button (out of 4 buttons) | crtmean_w2 |

Table S5 UKB cognitive test summary statistics, and latent cognitive ability model estimates (for all paths to the latent factor, p < .001).

| Cognitive Test | *N* | *M* (*SD*) | β (*SE*) | Residual variance |
|---|---|---|---|---|
| Reaction time (log) | 35138 | 6.38 (0.17) | 0.22 (0.006) | 0.84 |
| Numeric memory | 25720 | 6.76 (1.27) | -0.46 (0.006) | 0.76 |
| Fluid intelligence | 34709 | 6.60 (2.05) | -0.70 (0.004) | 0.49 |
| Trail making B (log) | 24484 | 6.28 (0.36) | 0.59 (0.005) | 0.49 |
| Matrix pattern | 25135 | 7.95 (2.14) | -0.58 (0.005) | 0.60 |
| Tower task | 24909 | 9.86 (3.23) | -0.49 (0.006) | 0.71 |
| Digit-symbol substitution | 25134 | 18.87 (5.27) | -0.44 (0.005) | 0.62 |
| Pairs matching (log) | 35365 | 1.35 (0.63) | 0.24 (0.006) | 0.92 |
| Prospective memory | 35350 | 0.84 (0.37) | 0.31 (0.006) | 0.88 |
| Paired associates | 25404 | 7.90 (02.64) | -0.44 (0.006) | 0.75 |

Table S6 GenScot cognitive test summary statistics and latent cognitive ability model estimates (for all paths to the latent factor, p < .001).

| Cognitive Test | *N* | *M* (*SD*) | β (*SE*) | Residual variance |
|---|---|---|---|---|
| Matrix reasoning | 1043 | 8.30 (2.39) | 0.56 (0.029) | 0.65 |
| Verbal fluency | 1043 | 43.10 (11.92) | 0.53 (0.030) | 0.71 |
| Mill Hill vocabulary | 1043 | 31.64 (4.07) | 0.70 (0.028) | 0.45 |
| Digit symbol substitution | 1043 | 68.77 (15.13) | 0.37 (0.039) | 0.64 |
| Logical memory | 1043 | 31.91 (7.23) | 0.47 (0.030) | 0.72 |

Table S7 LBC1936 cognitive test summary statistics, and cognitive ability model estimates (for all paths, besides the path between Verbal Memory and Cognitive ability which was fixed, all $p < .001$).

| Cognitive test | *N* | *M (SD)* | β (*SE*) | Residual variance |
|---|---|---|---|---|
| Matrix reasoning | 634 | 13.52 (4.93) | 0.60 (0.03) | 0.62 |
| Block design | 634 | 34.38 (10.01) | 0.60 (0.03) | 0.6 |
| Spatial span | 634 | 14.79 (2,72) | 0.45 (0.04) | 0.77 |
| NART | 634 | 34.66 (8.10) | 0.63 (0.03) | 0.57 |
| WTAR | 634 | 41.27 (6.94) | 0.65 (0.03) | 0.56 |
| Phonemic verbal fluency | 635 | 43.55 (12.78) | 0.47 (0.04) | 0.77 |
| Verbal paired associates | 623 | 27.57 (9.48) | 0.51 (0.04) | 0.7 |
| Logical memory | 635 | 75.03 (17.84) | 0.52 (0.04) | 0.71 |
| Digit span backwards | 636 | 7.88 (2.31) | 0.55 (0.04) | 0.69 |
| Symbol search | 634 | 24.88 (6.05) | 0.58 (0.03) | 0.64 |
| Digit-symbol substitution | 634 | 56.68 (11.79) | 0.61 (0.03) | 0.57 |
| Inspection time | 634 | 111.78 (10.95) | 0.38 (0.04) | 0.84 |
| Four-choice reaction time (s) | 635 | 0.64 (0.08) | 0.39 (0.04) | 0.82 |

Table S8 Within-domain residual variances for LBC1936 general cognitive ability model.

| Cognitive test | β (*SE*) |
|---|---|
| Matrix reasoning ~~ block design | 0.25 (0.05) |
| Matrix reasoning ~~ spatial span | 0.09 (0.05) |
| Block design ~~ spatial span | 0.20 (0.05) |
| NART ~~ WTAR | 0.83 (0.02) |
| NART~~ verbal fluency | 0.16 (0.05) |
| WTAR ~~ verbal fluency | 0.16 (0.05) |
| Verbal paired associates ~~ logical memory | 0.35 (0.04) |
| Verbal paired associates ~~ digit span backward | -0.04 (0.05) |
| Logical memory ~~ digit span backward | 0.01 (0.05) |
| Symbol search ~~ digit symbol | 0.40 (0.04) |
| Symbol search ~~ inspection time | 0.17 (0.04) |
| Symbol search ~~ choice reaction time | 0.31 (0.04) |
| Digit symbol ~~ inspection time | 0.22 (0.04) |
| Digit symbol ~~ choice reaction time | 0.36 (0.04) |
| Inspection time ~~ choice reaction time | 0.25 (0.04) |

Table S9 Model fits for the latent cognitive ability models.

| Cohort | $X^2$ | *df* | CFI | TLI | RMSEA | SRMR |
|---|---|---|---|---|---|---|
| UKB | 2417 | 34 | 0.953 | 0.913 | 0.042 | 0.026 |
| GenScot | 1170 | 20 | 0.972 | 0.887 | 0.079 | 0.019 |
| LBC1936 | 168 | 35 | 0.966 | 0.930 | 0.061 | 0.037 |

Table S10 Between-cohort spatial correlations r for vertex-wise mean profiles for each measure. All $p < 2.2x10^{-16}$.

| Measure | *r* LBC-GenScot | *r* GenScot-UKB | *r* UKB-LBC |
|---|---|---|---|
| Volume | 0.986 | 0.966 | 0.961 |
| Surface area | 0.990 | 0.977 | 0.969 |
| Thickness | 0.934 | 0.907 | 0.843 |
| Curvature | 0.997 | 0.968 | 0.972 |
| Sulcal depth | 0.998 | 0.993 | 0.990 |

Table S11 Spatial correlations (Pearson's r) between absolute value g-associations (β) for the 5 vertex-wise measures (all $p < 2.2x10^{-16}$).

| | | Volume | Surface area | Thickness | Curvature |
|---|---|---|---|---|---|
| *g* | Volume | 1 | | | |
| | Surface area | 0.653 | 1 | | |
| | Thickness | 0.413 | -0.077 | 1 | |
| | Curvature | -0.093 | 0.113 | -0.236 | 1 |
| | Sulcal depth | 0.032 | 0.241 | -0.151 | 0.182 |
| Age | Volume | 1 | | | |
| | Surface area | 0.550 | 1 | | |
| | Thickness | 0.380 | -0.350 | 1 | |
| | Curvature | -0.248 | 0.006 | -0.127 | 1 |
| | Sulcal depth | -0.064 | 0.019 | -0.182 | 0.308 |
| Sex | Volume | 1 | | | |
| | Surface area | 0.769 | 1 | | |
| | Thickness | -0.426 | -0.166 | 1 | |
| | Curvature | -0.097 | 0.163 | 0.203 | 1 |
| | Sulcal depth | 0.126 | 0.207 | 0.024 | 0.237 |

*Table S12* Brief descriptions of the strongest meta-analysed vertex-wise mappings

| | g | | Age | | Sex | |
|---|---|---|---|---|---|---|
| Measure | +ve | -ve | +ve | -ve | +ve (male > female) | -ve (female > male) |
| Volume | Superior frontal, medial frontal, lateral temporal, parietal | - | | Lateral temporal, medial frontal, ventrolateral prefrontal | Insula, fusiform gyrus, superior frontal | |

| | | | | | | |
|---|---|---|---|---|---|---|
| Surface area | Superior frontal, medial frontal, medial orbitofrontal, anterior cingulate, lateral temporal, parietal regions | - | | Lateral temporal | Insula, fusiform, frontal | |
| Thickness | Temporal pole, entorhinal cortex, precentral | Anterior cingulate, medial orbitofrontal, medial occipital | | Dorsolateral prefrontal, superior temporal, fusiform gyrus | Lateral temporal, medial orbitofrontal | Superior frontal and parietal regions |
| Curvature | Medial frontal, medial occipital | Anterior cingulate | Insula | Medial occipital, temporal, superior frontal gyrus | Precentral, medial frontal, medial occipital | Anterior cingulate |
| Sulcal depth | Medial frontal, temporal pole | Cingulate, hippocampal gyrus, parieto-frontal regions | Anterior cingulate, medial frontal, insula | Medial orbitofrontal, posterior cingulate, lateral orbitofrontal | Medial frontal, medial occipital | Insula, caudate anterior, posterior and isthmus cingulate, lateral temporal |

Table S13 Descriptive statistics for the multi-scale cortical profiles. The fsaverage surface is not symmetrical, so there is no direct correspondence of the position of vertices on left and right hemispheres. Therefore, to calculate the left and right spatial correlations, we took the mean of the values (for each measure) at each of the 68 Desikan-Killiany regions (34 per region) and correlated the left and right values). For subcortical measures, out of the 42 structures, 26 were part of left/right pairings (13 left and 13 right) – it is these that are correlated in the L vs R column here. The cytoarchitectural, functional and microstructural eigenvectors are in the units they come in through BigBrainWarp. The neurotransmitter receptor density profiles are scaled here.

| Measure | *N* regions/vertices | Measure | M | SD | Min | Max | Range | Skew | Kurtosis | L vs R (*r*) | % +ve | % FDR Q < .05 |
|---|---|---|---|---|---|---|---|---|---|---|---|---|
| g_ global_subcortical | 42 | β | 0.09 | 0.07 | -0.06 | 0.19 | 0.25 | -0.67 | -0.57 | 0.970 | - | - |
| Age_global_subcortical | 42 | β | -0.11 | 0.28 | -0.43 | 0.49 | 0.92 | 0.90 | -0.79 | 0.993 | - | - |
| Sex_global_subcortical | 42 | β | 0.30 | 0.12 | 0.03 | 0.49 | 0.46 | -0.42 | -0.69 | 0.985 | - | - |
| g_Volume | 298790 | β | 0.09 | 0.02 | 0.00 | 0.17 | 0.17 | 0.50 | 0.32 | 0.870 | 100 | 99 |
| g_Surface area | 298790 | β | 0.09 | 0.02 | 0.01 | 0.15 | 0.14 | 0.37 | -0.15 | 0.855 | 100 | 100 |
| g_Thickness | 298790 | β | 0.03 | 0.03 | -0.08 | 0.13 | 0.21 | 0.21 | 1.18 | 0.940 | 89 | 27 |
| g_Curvature | 298790 | β | 0.02 | 0.02 | -0.10 | 0.09 | 0.19 | -0.43 | 1.44 | 0.948 | 78 | 40 |
| g_Sulcal depth | 298790 | β | 0.00 | 0.03 | -0.12 | 0.13 | 0.24 | 0.14 | 1.00 | 0.950 | 54 | 26 |
| Age_Volume | 298790 | β | -0.16 | 0.05 | -0.32 | 0.03 | 0.36 | 0.28 | 0.37 | 0.918 | 0 | 92 |
| Age_Surface area | 298790 | β | -0.07 | 0.05 | -0.21 | 0.11 | 0.32 | 0.24 | 0.34 | 0.855 | 8 | 59 |
| Age_Thickness | 298790 | β | -0.20 | 0.07 | -0.36 | 0.06 | 0.42 | 0.73 | 0.32 | 0.949 | 1 | 91 |
| Age_Curvature | 298790 | β | -0.03 | 0.05 | -0.29 | 0.17 | 0.46 | -0.65 | 2.57 | 0.948 | 20 | 55 |
| Age_Sulcal depth | 298790 | β | 0.01 | 0.06 | -0.21 | 0.23 | 0.44 | -0.08 | 0.82 | 0.881 | 57 | 45 |
| Sex_Volume | 298790 | β | 0.26 | 0.07 | 0.01 | 0.45 | 0.44 | -0.15 | -0.29 | 0.948 | 100 | 99 |
| Sex_Surface area | 298790 | β | 0.31 | 0.05 | 0.09 | 0.46 | 0.38 | 0.11 | -0.18 | 0.927 | 100 | 100 |
| Sex_Thickness | 298790 | β | -0.03 | 0.06 | -0.23 | 0.15 | 0.39 | 0.19 | -0.02 | 0.938 | 28 | 61 |
| Sex_Curvature | 298790 | β | 0.06 | 0.08 | -0.21 | 0.30 | 0.52 | -0.20 | -0.19 | 0.867 | 78 | 62 |
| Sex_Sulcal depth | 298790 | B | 0.02 | 0.08 | -0.24 | 0.25 | 0.49 | -0.08 | -0.20 | 0.936 | 62 | 69 |
| Allometric scaling | 296637 | β | 0.60 | 0.08 | 0.30 | 0.82 | 0.52 | 0.04 | -0.49 | 0.983 | 100 | - |
| Metabolism | 296637 | Principal component | 0.00 | 1.63 | -9.13 | 5.60 | 14.74 | -0.52 | 1.51 | 0.986 | 52 | - |
| Cytoarchitecture 1 | 298790 | Eigenvector | 0.00 | 0.31 | -0.52 | 0.69 | 1.21 | 0.26 | -1.01 | 0.954 | 47 | - |
| Cytoarchitecture 2 | 298790 | Eigenvector | 0.04 | 0.21 | -0.44 | 0.50 | 0.94 | -0.12 | -0.77 | 0.963 | 58 | - |

| | | | | | | | | | | | |
|---|---|---|---|---|---|---|---|---|---|---|---|
| Functional 1 | 298790 | Eigenvector | 0.02 | 0.28 | -0.88 | 0.52 | 1.40 | -0.95 | 0.11 | 0.982 | 63 | - |
| Functional 2 | 298790 | Eigenvector | 0.01 | 0.15 | -0.37 | 0.26 | 0.62 | -0.62 | -0.61 | 0.994 | 62 | - |
| Microstructure 1 | 298790 | Eigenvector | 0.00 | 0.07 | -0.10 | 0.16 | 0.26 | 0.54 | -0.90 | 0.982 | 43 | - |
| Microstructure 2 | 298790 | Eigenvector | 0.00 | 0.01 | -0.02 | 0.03 | 0.05 | -0.04 | -0.77 | 0.843 | 49 | - |
| 5HT1a | 298790 | Receptor density (z) | 0.00 | 1.00 | -2.27 | 4.37 | 6.64 | 0.72 | 1.05 | 0.991 | 46 | - |
| 5HT1b | 298790 | Receptor density (z) | 0.00 | 1.00 | -3.22 | 3.79 | 7.01 | -0.11 | 0.52 | 0.991 | 51 | - |
| 5HT2a | 298790 | Receptor density (z) | 0.00 | 1.00 | -2.42 | 2.87 | 5.29 | 0.31 | -0.66 | 0.956 | 47 | - |
| 5HT4 | 298790 | Receptor density (z) | 0.00 | 1.00 | -2.14 | 3.08 | 5.22 | 0.46 | -0.46 | 0.969 | 45 | - |
| 5HT6 | 298790 | Receptor density (z) | 0.00 | 1.00 | -4.83 | 2.88 | 7.71 | -1.06 | 2.02 | 0.943 | 58 | - |
| 5HTT | 298790 | Receptor density (z) | 0.00 | 1.00 | -1.52 | 4.32 | 5.84 | 1.33 | 1.83 | 0.952 | 39 | - |
| D1 | 298790 | Receptor density (z) | 0.00 | 1.00 | -2.78 | 3.18 | 5.96 | -0.09 | -0.38 | 0.962 | 54 | - |
| D2 | 298790 | Receptor density (z) | 0.00 | 1.00 | -1.66 | 5.47 | 7.13 | 1.26 | 2.20 | 0.962 | 39 | - |
| DAT | 298790 | Receptor density (z) | 0.00 | 1.00 | -2.46 | 4.40 | 6.85 | 0.63 | 1.22 | 0.993 | 49 | - |
| NAT | 298790 | Receptor density (z) | 0.00 | 1.00 | -3.34 | 3.18 | 6.52 | -0.02 | 0.32 | 0.982 | 52 | - |
| H3 | 298790 | Receptor density (z) | 0.00 | 1.00 | -2.35 | 3.86 | 6.21 | 0.78 | 0.50 | 0.818 | 44 | - |
| A4B2 | 298790 | Receptor density (z) | 0.00 | 1.00 | -4.27 | 3.45 | 7.72 | -0.47 | 0.70 | 0.980 | 52 | - |
| M1 | 298790 | Receptor density (z) | 0.00 | 1.00 | -4.33 | 2.36 | 6.69 | -0.85 | 0.94 | 0.993 | 56 | - |
| VAChT | 298790 | Receptor density (z) | 0.00 | 1.00 | -4.42 | 3.71 | 8.13 | 0.01 | 0.61 | 0.972 | 50 | - |
| CB1 | 298790 | Receptor density (z) | 0.00 | 1.00 | -2.86 | 2.51 | 5.37 | -0.17 | -0.60 | 0.991 | 51 | - |
| MU | 298790 | Receptor density (z) | 0.00 | 1.00 | -2.44 | 1.93 | 4.37 | -0.67 | -0.22 | 0.972 | 58 | - |
| NMDA | 298790 | Receptor density (z) | 0.00 | 1.00 | -6.47 | 2.98 | 9.45 | -1.20 | 4.93 | 0.872 | 56 | - |
| mGluR5 | 298790 | Receptor density (z) | 0.00 | 1.00 | -1.80 | 6.80 | 8.60 | 2.41 | 9.59 | 0.996 | 42 | - |
| GABAa-bz | 298790 | Receptor density (z) | 0.00 | 1.00 | -3.41 | 3.10 | 6.50 | -0.19 | 0.34 | 0.997 | 54 | - |

Table S14 Descriptive statistics of estimates within each cohort, and the between-cohort relative correlations (Pearson's r) of the 46 global and subcortical estimates (all $p < .05$). Note, the summary estimates are for the standardised betas.

| | LBC1936 | | | GenScot | | | UKB | | | Between-cohort relative consistency (*r*) | | |
|---|---|---|---|---|---|---|---|---|---|---|---|---|
| | Mean (SD) | Min | Max | Mean (SD) | Min | Max | Mean (SD) | Min | Max | LBC1936-GenScot | GenScot-UKB | UKB-LBC1936 |
| *g* | 0.127 (0.116) | -0.139 | 0.283 | 0.091 (0.057) | -0.051 | 0.174 | 0.072 (0.045) | -0.025 | 0.133 | 0.724 | 0.858 | 0.820 |
| Age | - | - | - | -0.134 (0.270) | -0.401 | 0.460 | -0.094 (0.283) | -0.452 | 0.502 | - | 0.967 | - |
| Sex | 0.260 (0.121) | -0.016 | 0.468 | 0.275 (0.122) | 0.074 | 0.469 | 0.345 (0.142) | 0.016 | 0.568 | 0.795 | 0.914 | 0.825 |
| Allometry | 0.581 (0.094) | 0.232 | 0.843 | 0.593 (0.096) | 0.299 | 0.853 | 0.599 (0.088) | 0.012 | 0.822 | 0.754 | 0.726 | 0.779 |

*Table S15* Meta-analysis outcomes for ***g*** ~ global and subcortical brain structure volume associations (ordered by decreasing beta values).

| Region | β | *SE* | *z* | *p* | *FDR Q* | Cochrane's *Q* | *p* (*Q*) | $I^2$ |
|---|---|---|---|---|---|---|---|---|
| Total grey matter | 0.191 | 0.044 | 4.319 | 0.000 | 0.000 | 26.144 | 0.000 | 0.005 |
| Cerebral GM | 0.183 | 0.042 | 4.372 | 0.000 | 0.000 | 24.207 | 0.000 | 0.005 |
| Cerebral WM | 0.180 | 0.043 | 4.174 | 0.000 | 0.000 | 22.571 | 0.000 | 0.005 |
| TBV | 0.174 | 0.037 | 4.729 | 0.000 | 0.000 | 16.617 | 0.000 | 0.004 |
| Subcortical GM | 0.156 | 0.024 | 6.420 | 0.000 | 0.000 | 8.029 | 0.018 | 0.001 |
| Brain stem | 0.148 | 0.028 | 5.243 | 0.000 | 0.000 | 12.785 | 0.002 | 0.002 |
| Left ventral DC | 0.148 | 0.035 | 4.231 | 0.000 | 0.000 | 12.320 | 0.002 | 0.003 |
| Right Cerebellum GM | 0.146 | 0.032 | 4.592 | 0.000 | 0.000 | 10.217 | 0.006 | 0.003 |
| Right hippocampus | 0.141 | 0.028 | 5.016 | 0.000 | 0.000 | 9.691 | 0.008 | 0.002 |
| Right ventral DC | 0.139 | 0.027 | 5.245 | 0.000 | 0.000 | 8.395 | 0.015 | 0.002 |
| Left Cerebellum GM | 0.133 | 0.021 | 6.312 | 0.000 | 0.000 | 5.356 | 0.069 | 0.001 |
| CC posterior | 0.131 | 0.039 | 3.332 | 0.001 | 0.001 | 16.077 | 0.000 | 0.004 |
| CC anterior | 0.127 | 0.051 | 2.506 | 0.012 | 0.017 | 19.777 | 0.000 | 0.007 |
| Left Cerebellum WM | 0.123 | 0.022 | 5.678 | 0.000 | 0.000 | 5.942 | 0.051 | 0.001 |
| CC mid posterior | 0.122 | 0.053 | 2.284 | 0.022 | 0.029 | 24.455 | 0.000 | 0.008 |
| Left hippocampus | 0.117 | 0.011 | 10.370 | 0.000 | 0.000 | 2.020 | 0.364 | 0.000 |
| Right pallidum | 0.115 | 0.031 | 3.680 | 0.000 | 0.000 | 7.803 | 0.020 | 0.002 |
| Left pallidum | 0.113 | 0.040 | 2.798 | 0.005 | 0.007 | 10.759 | 0.005 | 0.004 |
| Left thalamus | 0.111 | 0.004 | 24.822 | 0.000 | 0.000 | 1.950 | 0.377 | 0.000 |
| Right putamen | 0.111 | 0.029 | 3.786 | 0.000 | 0.000 | 11.547 | 0.003 | 0.002 |
| Left putamen | 0.111 | 0.028 | 4.019 | 0.000 | 0.000 | 10.393 | 0.006 | 0.002 |
| Right thalamus | 0.110 | 0.004 | 26.765 | 0.000 | 0.000 | 1.336 | 0.513 | 0.000 |
| CC mid anterior | 0.107 | 0.054 | 1.985 | 0.047 | 0.056 | 22.577 | 0.000 | 0.008 |
| CC central | 0.106 | 0.063 | 1.689 | 0.091 | 0.101 | 30.329 | 0.000 | 0.011 |
| Right Cerebellum WM | 0.102 | 0.020 | 5.100 | 0.000 | 0.000 | 4.562 | 0.102 | 0.001 |
| Right amygdala | 0.095 | 0.004 | 21.707 | 0.000 | 0.000 | 1.862 | 0.394 | 0.000 |
| Right accumbens | 0.093 | 0.029 | 3.243 | 0.001 | 0.002 | 6.352 | 0.042 | 0.002 |
| Left amygdala | 0.090 | 0.011 | 8.533 | 0.000 | 0.000 | 2.308 | 0.315 | 0.000 |
| Left caudate | 0.075 | 0.019 | 3.945 | 0.000 | 0.000 | 4.180 | 0.124 | 0.001 |
| Left accumbens | 0.067 | 0.005 | 14.578 | 0.000 | 0.000 | 3.582 | 0.167 | 0.000 |
| Right caudate | 0.063 | 0.005 | 13.529 | 0.000 | 0.000 | 1.585 | 0.453 | 0.000 |
| 4th ventricle | 0.035 | 0.005 | 7.426 | 0.000 | 0.000 | 3.583 | 0.167 | 0.000 |
| Optic chiasm | 0.014 | 0.004 | 3.123 | 0.002 | 0.003 | 0.159 | 0.923 | 0.000 |
| CSF | 0.012 | 0.041 | 0.305 | 0.760 | 0.779 | 10.487 | 0.005 | 0.004 |
| Left choroid plexus | 0.011 | 0.007 | 1.725 | 0.085 | 0.096 | 2.176 | 0.337 | 0.000 |
| Right choroid plexus | 0.009 | 0.004 | 2.171 | 0.030 | 0.038 | 0.069 | 0.966 | 0.000 |
| Right lateral ventricle | 0.001 | 0.005 | 0.182 | 0.856 | 0.856 | 1.595 | 0.451 | 0.000 |
| Left lateral ventricle | -0.002 | 0.005 | -0.551 | 0.581 | 0.611 | 2.527 | 0.283 | 0.000 |
| 3rd ventricle | -0.026 | 0.016 | -1.645 | 0.100 | 0.108 | 4.074 | 0.130 | 0.000 |
| Left inferior lateral ventricle | -0.028 | 0.004 | -6.383 | 0.000 | 0.000 | 1.056 | 0.590 | 0.000 |
| Right inferior lateral ventricle | -0.057 | 0.027 | -2.130 | 0.033 | 0.041 | 9.015 | 0.011 | 0.002 |
| WM hypointensities | -0.063 | 0.032 | -1.981 | 0.048 | 0.056 | 9.619 | 0.008 | 0.002 |

*Table S16* Meta-analysis outcomes for global and subcortical brain structure volume ~ **age** associations (ordered by decreasing beta values).

| Region | β | *SE* | *z* | *p* | *FDR Q* | Cochrane's *Q* | *p* (*Q*) | $I^2$ |
|---|---|---|---|---|---|---|---|---|
| Left accumbens | -0.431 | 0.024 | -17.823 | 0.000 | 0.000 | 4.781 | 0.029 | 0.001 |
| Right accumbens | -0.375 | 0.004 | -92.348 | 0.000 | 0.000 | 0.284 | 0.594 | 0.000 |
| Left hippocampus | -0.364 | 0.004 | -90.425 | 0.000 | 0.000 | 0.022 | 0.881 | 0.000 |
| Right hippocampus | -0.363 | 0.004 | -89.332 | 0.000 | 0.000 | 0.714 | 0.398 | 0.000 |
| Left amygdala | -0.342 | 0.036 | -9.608 | 0.000 | 0.000 | 7.419 | 0.006 | 0.002 |
| Left thalamus | -0.336 | 0.024 | -13.798 | 0.000 | 0.000 | 4.508 | 0.034 | 0.001 |
| CC mid anterior | -0.333 | 0.004 | -75.824 | 0.000 | 0.000 | 0.102 | 0.750 | 0.000 |
| Right thalamus | -0.328 | 0.014 | -24.056 | 0.000 | 0.000 | 1.981 | 0.159 | 0.000 |
| Right ventral DC | -0.327 | 0.039 | -8.372 | 0.000 | 0.000 | 11.414 | 0.001 | 0.003 |
| Total grey matter | -0.319 | 0.036 | -8.883 | 0.000 | 0.000 | 12.571 | 0.000 | 0.002 |
| CC central | -0.315 | 0.018 | -17.378 | 0.000 | 0.000 | 2.258 | 0.133 | 0.000 |
| Cerebral WM | -0.307 | 0.030 | -10.158 | 0.000 | 0.000 | 8.086 | 0.004 | 0.002 |
| Left putamen | -0.306 | 0.060 | -5.070 | 0.000 | 0.000 | 24.243 | 8.49E-07 | 0.007 |
| Right putamen | -0.306 | 0.061 | -4.978 | 0.000 | 0.000 | 26.496 | 2.64E-07 | 0.007 |
| CC mid posterior | -0.301 | 0.061 | -4.962 | 0.000 | 0.000 | 21.761 | 3.09E-06 | 0.007 |
| Right amygdala | -0.287 | 0.004 | -69.675 | 0.000 | 0.000 | 0.447 | 0.504 | 0.000 |
| Left ventral DC | -0.285 | 0.015 | -19.026 | 0.000 | 0.000 | 1.989 | 0.158 | 0.000 |
| Subcortical GM | -0.281 | 0.062 | -4.545 | 0.000 | 0.000 | 31.543 | 1.95E-08 | 0.007 |
| TBV | -0.250 | 0.051 | -4.861 | 0.000 | 0.000 | 14.376 | 0.000 | 0.005 |
| Left Cerebellum GM | -0.231 | 0.065 | -3.530 | 0.000 | 0.000 | 32.031 | 1.52E-08 | 0.008 |
| Left Cerebellum WM | -0.228 | 0.072 | -3.181 | 0.001 | 0.002 | 26.952 | 2.09E-07 | 0.010 |
| Right Cerebellum WM | -0.224 | 0.052 | -4.313 | 0.000 | 0.000 | 13.029 | 0.000 | 0.005 |
| Cerebral GM | -0.220 | 0.004 | -54.317 | 0.000 | 0.000 | 0.503 | 0.478 | 0.000 |
| Right Cerebellum GM | -0.219 | 0.047 | -4.626 | 0.000 | 0.000 | 18.329 | 1.86E-05 | 0.004 |
| CC anterior | -0.211 | 0.054 | -3.907 | 0.000 | 0.000 | 14.619 | 0.000 | 0.005 |
| Left pallidum | -0.152 | 0.005 | -33.826 | 0.000 | 0.000 | 0.003 | 0.960 | 0.000 |
| Brain stem | -0.142 | 0.035 | -4.087 | 0.000 | 0.000 | 7.097 | 0.008 | 0.002 |
| Right pallidum | -0.137 | 0.013 | -10.335 | 0.000 | 0.000 | 1.491 | 0.222 | 0.000 |
| CC posterior | -0.090 | 0.066 | -1.366 | 0.172 | 0.176 | 19.031 | 1.29E-05 | 0.008 |
| Left caudate | -0.082 | 0.057 | -1.445 | 0.148 | 0.156 | 15.094 | 0.000 | 0.006 |
| Right caudate | -0.071 | 0.075 | -0.950 | 0.342 | 0.342 | 26.200 | 3.08E-07 | 0.011 |
| 4th ventricle | 0.115 | 0.013 | 8.551 | 0.000 | 0.000 | 1.460 | 0.227 | 0.000 |
| CSF | 0.219 | 0.004 | 49.383 | 0.000 | 0.000 | 0.063 | 0.802 | 0.000 |
| Optic chiasm | 0.227 | 0.004 | 51.311 | 0.000 | 0.000 | 0.321 | 0.571 | 0.000 |
| Right inferior lateral ventricle | 0.283 | 0.064 | 4.404 | 0.000 | 0.000 | 20.098 | 7.36E-06 | 0.008 |
| Left choroid plexus | 0.327 | 0.067 | 4.902 | 0.000 | 0.000 | 26.307 | 2.91E-07 | 0.009 |
| Left inferior lateral ventricle | 0.354 | 0.035 | 10.135 | 0.000 | 0.000 | 7.104 | 0.008 | 0.002 |
| Right choroid plexus | 0.358 | 0.043 | 8.394 | 0.000 | 0.000 | 11.995 | 0.001 | 0.003 |
| Left lateral ventricle | 0.364 | 0.015 | 23.712 | 0.000 | 0.000 | 1.890 | 0.169 | 0.000 |
| Right lateral ventricle | 0.378 | 0.017 | 22.423 | 0.000 | 0.000 | 2.150 | 0.143 | 0.000 |
| 3rd ventricle | 0.397 | 0.004 | 106.683 | 0.000 | 0.000 | 0.166 | 0.684 | 0.000 |
| WM hypointensities | 0.487 | 0.020 | 23.985 | 0.000 | 0.000 | 3.368 | 0.066 | 0.001 |

*Table S17* Meta-analysis outcomes global and subcortical brain structure volume ~ **sex** associations (ordered by decreasing beta values).

| Region | β | *SE* | *z* | *p* | *FDR Q* | Cochrane's *Q* | *p* (*Q*) | $I^2$ |
|---|---|---|---|---|---|---|---|---|
| Total grey matter | 0.490 | 0.043 | 11.475 | 0.000 | 0.000 | 46.446 | 0.000 | 0.005 |
| Subcortical GM | 0.466 | 0.054 | 8.569 | 0.000 | 0.000 | 48.029 | 0.000 | 0.008 |
| TBV | 0.466 | 0.053 | 8.743 | 0.000 | 0.000 | 51.550 | 0.000 | 0.008 |
| Cerebral WM | 0.462 | 0.045 | 10.214 | 0.000 | 0.000 | 27.210 | 0.000 | 0.006 |
| Cerebral GM | 0.455 | 0.042 | 10.918 | 0.000 | 0.000 | 36.697 | 0.000 | 0.005 |
| Brain stem | 0.447 | 0.034 | 12.968 | 0.000 | 0.000 | 15.692 | 0.000 | 0.003 |
| Right Cerebellum GM | 0.439 | 0.038 | 11.505 | 0.000 | 0.000 | 29.706 | 0.000 | 0.004 |
| Left ventral DC | 0.432 | 0.040 | 10.743 | 0.000 | 0.000 | 18.679 | 0.000 | 0.004 |
| Right ventral DC | 0.423 | 0.041 | 10.213 | 0.000 | 0.000 | 22.223 | 0.000 | 0.005 |
| Left Cerebellum GM | 0.420 | 0.039 | 10.793 | 0.000 | 0.000 | 21.197 | 0.000 | 0.004 |
| 3rd ventricle | 0.392 | 0.033 | 11.868 | 0.000 | 0.000 | 9.580 | 0.008 | 0.003 |
| Left amygdala | 0.366 | 0.018 | 19.988 | 0.000 | 0.000 | 4.112 | 0.128 | 0.001 |
| Right amygdala | 0.350 | 0.050 | 7.002 | 0.000 | 0.000 | 41.785 | 0.000 | 0.007 |
| Left choroid plexus | 0.347 | 0.022 | 15.447 | 0.000 | 0.000 | 5.002 | 0.082 | 0.001 |
| Right choroid plexus | 0.345 | 0.029 | 11.914 | 0.000 | 0.000 | 10.724 | 0.005 | 0.002 |
| Right thalamus | 0.344 | 0.056 | 6.116 | 0.000 | 0.000 | 59.414 | 0.000 | 0.009 |
| Right putamen | 0.339 | 0.043 | 7.859 | 0.000 | 0.000 | 26.758 | 0.000 | 0.005 |
| Left putamen | 0.338 | 0.048 | 7.088 | 0.000 | 0.000 | 30.591 | 0.000 | 0.006 |
| CSF | 0.336 | 0.017 | 20.136 | 0.000 | 0.000 | 3.324 | 0.190 | 0.000 |
| Left pallidum | 0.328 | 0.051 | 6.462 | 0.000 | 0.000 | 31.327 | 0.000 | 0.007 |
| Left thalamus | 0.323 | 0.043 | 7.581 | 0.000 | 0.000 | 27.543 | 0.000 | 0.005 |
| Right pallidum | 0.311 | 0.069 | 4.497 | 0.000 | 0.000 | 40.437 | 0.000 | 0.014 |
| Left hippocampus | 0.297 | 0.031 | 9.444 | 0.000 | 0.000 | 11.803 | 0.003 | 0.002 |
| Right hippocampus | 0.297 | 0.023 | 12.995 | 0.000 | 0.000 | 6.072 | 0.048 | 0.001 |
| Left inferior lateral ventricle | 0.285 | 0.020 | 14.590 | 0.000 | 0.000 | 4.337 | 0.114 | 0.001 |
| Right lateral ventricle | 0.274 | 0.005 | 59.872 | 0.000 | 0.000 | 2.519 | 0.284 | 0.000 |
| Right inferior lateral ventricle | 0.268 | 0.041 | 6.600 | 0.000 | 0.000 | 9.980 | 0.007 | 0.004 |
| Left lateral ventricle | 0.267 | 0.004 | 62.887 | 0.000 | 0.000 | 2.459 | 0.292 | 0.000 |
| 4th ventricle | 0.260 | 0.026 | 9.976 | 0.000 | 0.000 | 5.696 | 0.058 | 0.001 |
| Right caudate | 0.248 | 0.040 | 6.280 | 0.000 | 0.000 | 17.563 | 0.000 | 0.004 |
| Left caudate | 0.247 | 0.031 | 7.996 | 0.000 | 0.000 | 9.699 | 0.008 | 0.002 |
| Optic chiasm | 0.229 | 0.064 | 3.566 | 0.000 | 0.000 | 45.542 | 0.000 | 0.012 |
| Right accumbens | 0.192 | 0.051 | 3.794 | 0.000 | 0.000 | 37.051 | 0.000 | 0.007 |
| Right Cerebellum WM | 0.189 | 0.029 | 6.562 | 0.000 | 0.000 | 8.804 | 0.012 | 0.002 |
| Left Cerebellum WM | 0.178 | 0.044 | 4.031 | 0.000 | 0.000 | 23.445 | 0.000 | 0.005 |
| Left accumbens | 0.169 | 0.047 | 3.564 | 0.000 | 0.000 | 30.080 | 0.000 | 0.006 |
| WM hypointensities | 0.136 | 0.023 | 6.026 | 0.000 | 0.000 | 5.299 | 0.071 | 0.001 |
| CC posterior | 0.120 | 0.005 | 24.525 | 0.000 | 0.000 | 1.245 | 0.537 | 0.000 |
| CC anterior | 0.115 | 0.042 | 2.741 | 0.006 | 0.006 | 11.039 | 0.004 | 0.004 |
| CC mid anterior | 0.059 | 0.013 | 4.424 | 0.000 | 0.000 | 3.780 | 0.151 | 0.000 |
| CC mid posterior | 0.058 | 0.007 | 8.899 | 0.000 | 0.000 | 1.591 | 0.451 | 0.000 |
| CC central | 0.028 | 0.018 | 1.556 | 0.120 | 0.120 | 3.409 | 0.182 | 0.000 |

*Table S18* **Between-cohort age moderation outcomes** for meta-analysis of ***g*** ~ global and subcortical brain structure volume associations (ordered by decreasing beta values, structures for which *FDR Q* < .05 are in bold font).

| Region | β | *SE* | *z* | *p* | *FDR Q* | Cochrane's *Q* | *p* (*Q*) | $I^2$ |
|---|---|---|---|---|---|---|---|---|
| CC central | 0.015 | 0.008 | 1.894 | 0.058 | 0.321 | 13.890 | 0.000 | 0.004 |
| CC mid anterior | 0.012 | 0.007 | 1.764 | 0.078 | 0.327 | 10.980 | 0.001 | 0.003 |
| CC anterior | 0.012 | 0.006 | 1.964 | 0.050 | 0.321 | 7.935 | 0.005 | 0.002 |
| CC mid posterior | 0.012 | 0.008 | 1.541 | 0.123 | 0.351 | 14.066 | 0.000 | 0.004 |
| **Left pallidum** | **0.011** | **0.003** | **3.205** | **0.001** | **0.028** | **0.488** | **0.485** | **0.000** |
| Total GM | 0.010 | 0.007 | 1.450 | 0.147 | 0.363 | 17.900 | 0.000 | 0.003 |
| Cerebral WM | 0.009 | 0.007 | 1.330 | 0.184 | 0.386 | 14.642 | 0.000 | 0.004 |
| Cerebral GM | 0.009 | 0.007 | 1.235 | 0.217 | 0.396 | 18.346 | 0.000 | 0.004 |
| Right pallidum | 0.009 | 0.004 | 2.207 | 0.027 | 0.287 | 1.850 | 0.174 | 0.000 |
| Right accumbens | 0.009 | 0.003 | 2.468 | 0.014 | 0.190 | 0.262 | 0.609 | 0.000 |
| Left ventral DC | 0.008 | 0.005 | 1.669 | 0.095 | 0.333 | 6.221 | 0.013 | 0.001 |
| TBV | 0.008 | 0.005 | 1.527 | 0.127 | 0.351 | 7.914 | 0.005 | 0.002 |
| Right cerebellum GM | 0.008 | 0.004 | 1.781 | 0.075 | 0.327 | 5.051 | 0.025 | 0.001 |
| CC posterior | 0.007 | 0.008 | 0.952 | 0.341 | 0.530 | 12.227 | 0.000 | 0.004 |
| Left accumbens | 0.006 | 0.003 | 1.880 | 0.060 | 0.321 | 0.047 | 0.828 | 0.000 |
| Right ventral DC | 0.006 | 0.005 | 1.338 | 0.181 | 0.386 | 5.102 | 0.024 | 0.001 |
| Left cerebellum GM | 0.006 | 0.004 | 1.684 | 0.092 | 0.333 | 2.084 | 0.149 | 0.000 |
| Subcortical GM | 0.005 | 0.005 | 1.172 | 0.241 | 0.405 | 5.852 | 0.016 | 0.001 |
| Right Cerebellum WM | 0.005 | 0.004 | 1.260 | 0.208 | 0.396 | 2.436 | 0.119 | 0.001 |
| Right hippocampus | 0.005 | 0.006 | 0.828 | 0.408 | 0.591 | 8.143 | 0.004 | 0.002 |
| Right putamen | 0.005 | 0.006 | 0.703 | 0.482 | 0.614 | 10.637 | 0.001 | 0.003 |
| Right amygdala | 0.004 | 0.003 | 1.355 | 0.176 | 0.386 | 0.027 | 0.870 | 0.000 |
| Left pallidum | 0.004 | 0.006 | 0.551 | 0.582 | 0.679 | 9.879 | 0.002 | 0.003 |
| Left amygdala | 0.003 | 0.004 | 0.729 | 0.466 | 0.612 | 1.707 | 0.191 | 0.000 |
| Left thalamus | 0.003 | 0.003 | 0.780 | 0.435 | 0.609 | 1.363 | 0.243 | 0.000 |
| Left cerebellum WM | 0.002 | 0.005 | 0.431 | 0.667 | 0.737 | 5.751 | 0.016 | 0.002 |
| Brain stem | 0.002 | 0.007 | 0.319 | 0.750 | 0.807 | 12.652 | 0.000 | 0.003 |
| Right thalamus | 0.002 | 0.003 | 0.563 | 0.573 | 0.679 | 1.021 | 0.312 | 0.000 |
| Left hippocampus | 0.000 | 0.004 | 0.070 | 0.944 | 0.944 | 2.020 | 0.155 | 0.000 |
| Optic chiasm | 0.000 | 0.004 | -0.092 | 0.926 | 0.944 | 0.151 | 0.698 | 0.000 |
| Right choroid plexus | -0.001 | 0.003 | -0.167 | 0.867 | 0.911 | 0.041 | 0.840 | 0.000 |
| Left inferior lateral ventricle | -0.002 | 0.003 | -0.660 | 0.509 | 0.629 | 0.620 | 0.431 | 0.000 |
| Right caudate | -0.003 | 0.004 | -0.732 | 0.464 | 0.612 | 1.039 | 0.308 | 0.000 |
| Right inferior lateral ventricle | -0.003 | 0.006 | -0.465 | 0.642 | 0.728 | 8.330 | 0.004 | 0.003 |
| Left caudate | -0.004 | 0.004 | -0.850 | 0.395 | 0.591 | 2.860 | 0.091 | 0.001 |
| Right lateral ventricle | -0.004 | 0.003 | -1.144 | 0.253 | 0.408 | 0.286 | 0.593 | 0.000 |
| Left choroid plexus | -0.004 | 0.003 | -1.206 | 0.228 | 0.398 | 0.721 | 0.396 | 0.000 |
| Left lateral ventricle | -0.005 | 0.003 | -1.499 | 0.134 | 0.351 | 0.280 | 0.597 | 0.000 |
| 3rd ventricle | -0.005 | 0.003 | -1.604 | 0.109 | 0.351 | 1.330 | 0.249 | 0.000 |
| 4th ventricle | -0.007 | 0.004 | -1.872 | 0.061 | 0.321 | 0.079 | 0.779 | 0.000 |
| WM hypointensities | -0.007 | 0.006 | -1.302 | 0.193 | 0.386 | 6.308 | 0.012 | 0.002 |
| **CSF** | **-0.011** | **0.003** | **-3.213** | **0.001** | **0.028** | **0.167** | **0.683** | **0.000** |

Table S19 **Between-cohort age moderation outcomes** for meta-analysis of global and subcortical brain structure volume ~ **sex** associations (ordered by decreasing beta values, FDR Q < .05 associations are in bold font).

| Region | B | *SE* | *z* | *p* | *FDR Q* | Cochrane's *Q* | *p* (*Q*) | $I^2$ |
|---|---|---|---|---|---|---|---|---|
| Right inferior lateral ventricle | 0.011 | 0.004 | 3.144 | 0.002 | 0.070 | 0.293 | 0.589 | 0.000 |
| Optic chiasm | 0.010 | 0.013 | 0.753 | 0.451 | 0.885 | 37.910 | 0.000 | 0.015 |
| 3rd ventricle | 0.009 | 0.003 | 2.894 | 0.004 | 0.080 | 1.177 | 0.278 | 0.000 |
| Left inferior lateral ventricle | 0.007 | 0.004 | 2.043 | 0.041 | 0.515 | 0.322 | 0.570 | 0.000 |
| Left accumbens | 0.007 | 0.010 | 0.700 | 0.484 | 0.885 | 23.248 | 0.000 | 0.008 |
| Right lateral ventricle | 0.006 | 0.004 | 1.607 | 0.108 | 0.649 | 0.030 | 0.863 | 0.000 |
| Left lateral ventricle | 0.005 | 0.004 | 1.472 | 0.141 | 0.705 | 0.437 | 0.509 | 0.000 |
| WM hypointensities | 0.005 | 0.005 | 1.025 | 0.305 | 0.885 | 3.118 | 0.077 | 0.001 |
| TBV | 0.004 | 0.013 | 0.266 | 0.790 | 0.889 | 52.072 | 0.000 | 0.015 |
| Right amygdala | 0.002 | 0.013 | 0.152 | 0.879 | 0.928 | 41.708 | 0.000 | 0.013 |
| Left amygdala | 0.000 | 0.005 | 0.036 | 0.972 | 0.992 | 4.541 | 0.033 | 0.001 |
| Right accumbens | 0.000 | 0.013 | 0.010 | 0.992 | 0.992 | 37.107 | 0.000 | 0.014 |
| CSF | -0.001 | 0.005 | -0.147 | 0.883 | 0.928 | 3.299 | 0.069 | 0.001 |
| CC posterior | -0.002 | 0.004 | -0.432 | 0.666 | 0.885 | 1.394 | 0.238 | 0.000 |
| Right Cerebellum WM | -0.002 | 0.007 | -0.306 | 0.759 | 0.886 | 9.262 | 0.002 | 0.003 |
| Right hippocampus | -0.002 | 0.006 | -0.392 | 0.695 | 0.885 | 6.589 | 0.010 | 0.002 |
| Right Cerebellum GM | -0.003 | 0.010 | -0.326 | 0.745 | 0.886 | 30.782 | 0.000 | 0.008 |
| CC mid posterior | -0.003 | 0.004 | -0.794 | 0.427 | 0.885 | 0.823 | 0.364 | 0.000 |
| Right choroid plexus | -0.003 | 0.007 | -0.494 | 0.622 | 0.885 | 10.429 | 0.001 | 0.003 |
| Right thalamus | -0.003 | 0.014 | -0.247 | 0.805 | 0.889 | 60.073 | 0.000 | 0.017 |
| Total grey matter | -0.004 | 0.011 | -0.344 | 0.731 | 0.886 | 46.853 | 0.000 | 0.010 |
| Left hippocampus | -0.004 | 0.008 | -0.495 | 0.621 | 0.885 | 12.092 | 0.001 | 0.004 |
| Right caudate | -0.004 | 0.009 | -0.430 | 0.667 | 0.885 | 17.350 | 0.000 | 0.007 |
| Left caudate | -0.004 | 0.007 | -0.591 | 0.555 | 0.885 | 9.246 | 0.002 | 0.003 |
| Left Cerebellum WM | -0.004 | 0.011 | -0.410 | 0.682 | 0.885 | 23.913 | 0.000 | 0.009 |
| Right putamen | -0.005 | 0.010 | -0.443 | 0.658 | 0.885 | 27.327 | 0.000 | 0.009 |
| Left thalamus | -0.005 | 0.010 | -0.451 | 0.652 | 0.885 | 28.376 | 0.000 | 0.009 |
| Cerebral WM | -0.005 | 0.010 | -0.461 | 0.645 | 0.885 | 37.239 | 0.000 | 0.009 |
| CC central | -0.005 | 0.004 | -1.175 | 0.240 | 0.875 | 1.609 | 0.205 | 0.000 |
| Left putamen | -0.006 | 0.011 | -0.527 | 0.598 | 0.885 | 30.545 | 0.000 | 0.010 |
| Brain stem | -0.006 | 0.007 | -0.860 | 0.390 | 0.885 | 14.411 | 0.000 | 0.004 |
| Left Cerebellum GM | -0.007 | 0.008 | -0.804 | 0.421 | 0.885 | 20.881 | 0.000 | 0.005 |
| Left choroid plexus | -0.007 | 0.004 | -1.779 | 0.075 | 0.527 | 1.530 | 0.216 | 0.000 |
| 4th ventricle | -0.007 | 0.005 | -1.436 | 0.151 | 0.705 | 2.834 | 0.092 | 0.001 |
| Left pallidum | -0.007 | 0.012 | -0.606 | 0.544 | 0.885 | 29.964 | 0.000 | 0.011 |
| CC mid anterior | -0.007 | 0.004 | -1.871 | 0.061 | 0.515 | 0.674 | 0.412 | 0.000 |
| Right ventral DC | -0.008 | 0.008 | -0.963 | 0.336 | 0.885 | 19.884 | 0.000 | 0.005 |
| Left ventral DC | -0.009 | 0.007 | -1.150 | 0.250 | 0.875 | 15.513 | 0.000 | 0.004 |
| Subcortical GM | -0.009 | 0.012 | -0.777 | 0.437 | 0.885 | 46.622 | 0.000 | 0.011 |
| Cerebral GM | -0.009 | 0.009 | -1.032 | 0.302 | 0.885 | 23.283 | 0.000 | 0.006 |
| CC anterior | -0.011 | 0.006 | -1.905 | 0.057 | 0.515 | 4.645 | 0.031 | 0.002 |
| Right pallidum | -0.014 | 0.012 | -1.247 | 0.213 | 0.875 | 30.823 | 0.000 | 0.011 |

[1] R.M. Reitan, & D. Wolfson (1985). The Halstead–Reitan neuropsycholgical test battery: Therapy and clinical interpretation. Neuropsychological Press, Tucson, AZ.

[2] Ritchie, K., Roquefeuil, *g.* de, Ritchie, C., *et al.*: COGNITO: computerized assessment of information processing. *J Psychol Psychother.* 2014; 4(2): 1000136.

[3] A. Smith. (1991). Symbol digit modalities test. Western Psychological Services, Los Angeles, CA

[4] https://biobank.ctsu.ox.ac.uk/crystal/crystal/docs/Pairs.pdf

[5] Benton A, Hamsher K, Sivan A: Multilingual aphasia examination. 3rd ed. San Antonio, TX: Psychological Corporation; 1994.

[6] Raven JC: Raven's progressive matrices and vocabulary scales. Oxford, England: Oxford Psychologists Press Ltd. 1989.

[7] Wechsler, D. (1997b) Wechsler Adult Intelligence Scale-Third Edition (WAIS-III). San Antonio: Harcourt Assessment Inc.

[8] Wechsler, D. (1997a) Wechsler Adult Intelligence Scale-Third Edition (WAIS-III). San Antonio: Harcourt Assessment Inc.

[9] Nelson, H.E. & Wilson, J. (1991). National Adult Reading Test (NART), Windsor, UK.

[10] Wechsler, D. (2001). Wechsler test of adult Reading. Psychological Corporation, San Antonio, TX.

[11] Lezak, M.D., Howieson, D.B., Loring, D.W., Hannay, H.J. & Fischer, J.S. (2004) Neuropsychological assessment (4th ed.), Oxford University Press, New York, NY.

[12] Smith, A. (1991). Symbol digit modalities test. Western Psychological Services, Los Angeles, CA

[13] Deary, I. J., Simonotto, E., Meyer, M., Marshall, A., Marshall, I., Goddard, N., & Wardlaw, J. M. (2004). The functional anatomy of inspection time: an event-related fMRI study. *NeuroIm*age, *22*(4), 1466–1479. https://doi.org/10.1016/j.neuroimage.2004.03.047

[14] Deary, I. J., Der, G., & Ford, *g.* (2001). Reaction times and intelligence differences: A population-based cohort study. *Intelligence*, 29 (5), 389-399. https://doi.org/10.1016/S0160-2896(01)00062-9
