## Supplementary Analyses for "Brain maps of general cognitive function and spatial correlations with neurobiological cortical profiles"

### Supplementary analysis 1: Smoothing tolerances

The smoothing tolerance chosen for vertex-wise analyses could have an impact on the results. Noise in the data, due to registration inaccuracies, is minimized when the cortex is parcellated into larger regions (i.e., greater smoothing) but, when the cortex is parcellated into smaller regions (i.e., less smoothing), the % variance explained increases (41F^[1]^) due to the additional information provided. Thus, at the vertex-wise level, there is a balance to be struck between the benefits of reducing noise in the data, and the problem that increasing to higher levels of smoothing will, at a point, remove fine-grained spatial information and thus reduce the spatial specificity of detected associations. Lerch and Evans (2005) analysed the effect of different smoothing tolerances on cortical thickness measurement sensitivity, and they concluded an optimal kernel size of 30 mm (42F^[2]^, *N =* 25). Some studies use 30 mm (43F^[3],^44F^[4]^), and other common choices are 5 mm (45F^[5]^), 10 mm (46F^[6],^47F^[7],^48F^[8]^), 15 mm (49F^[9],^50F^[10]^) or 20 mm (51F^[11],^52F^[12]^).

Here, we calculated the *g*-morphometry associations for each vertex-wise measure (volume, surface area, thickness, curvature and sulcal depth) for 9 smoothing kernels (0, 5, 10, 15, 20, 25, 30, 35 and 40 mm), see Supplementary Figures S4 to S6. The trade-off between noise reduction and loss of fine-grained information is clearest with standardised βs that are all in the same direction. In Figure i, the trade-off can be most clearly seen in the *g-*volume estimates – the means of the standardised βs increase with increasing smoothing tolerance alongside a u-shaped trend in maximum densities of the standardised βs with increasing smoothing tolerances (as localized results are first located through noise reduction and then lost as results converge towards the total surface effect). Generally, the optimum smoothing tolerance for these types of associations across different morphometry measures appears to be between 10-20 mm (see Figure iB). Therefore, the *a priori* selection of 20 fwhm for the main *g-*morphometry analyses in this paper, in line with our previous work (e.g. ^[13]^), appears appropriate. These results may aid future choices for similar analyses.

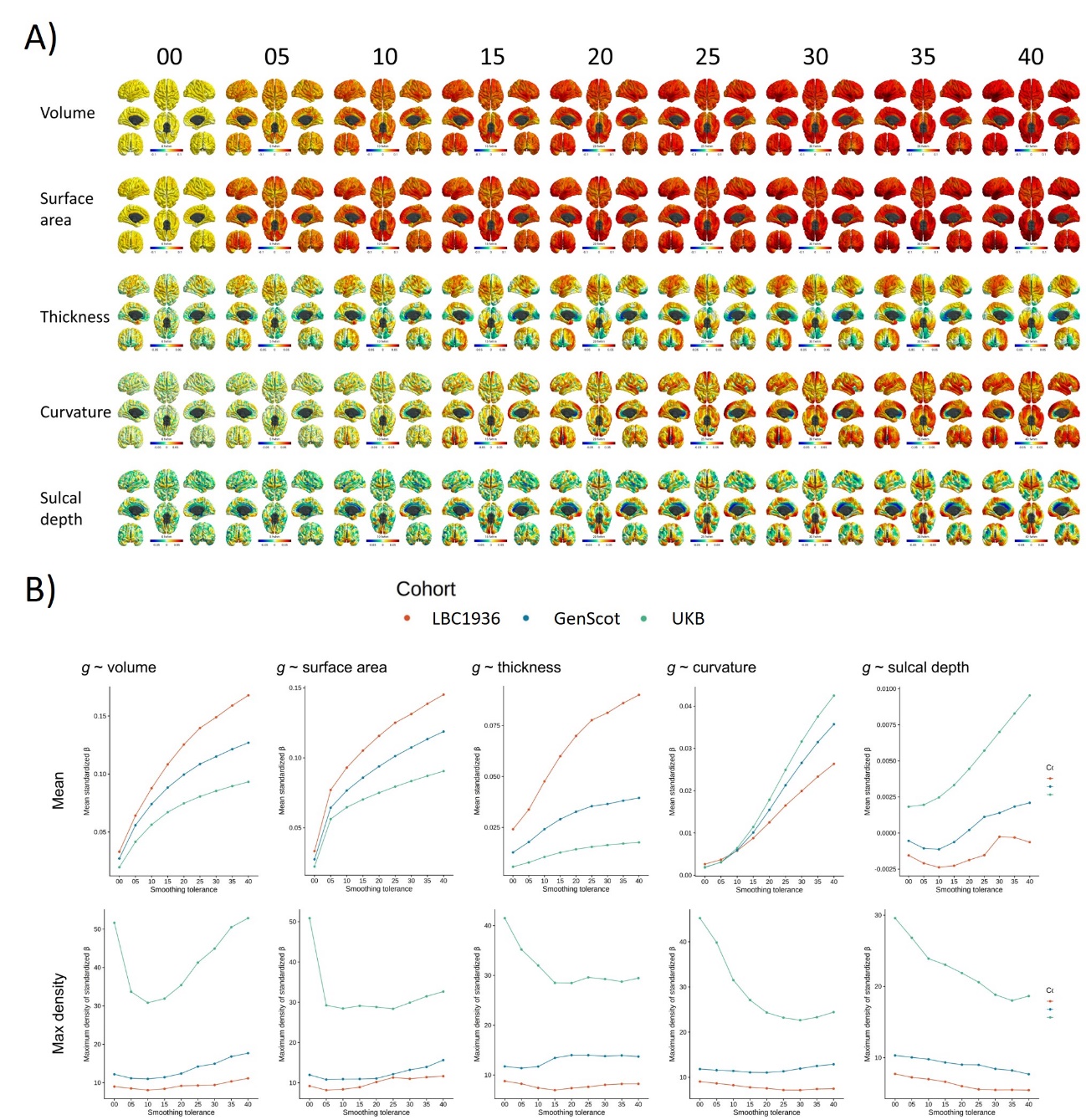

*Figure i* A) UKB *g-*morphometry associations at the 9 included smoothing tolerances. B) Summaries of the vertex-wise volume associations with *g* across smoothing tolerances. Top: mean standardised *β* at each smoothing tolerance; Bottom: maximum density of standardised *β* at each smoothing tolerance.

### Supplementary analysis 2: Global and subcortical volumes analyses

Although much previous work on *g-*brain associations focuses on the cortex, sub-cortical structures are becoming increasingly recognized for their associations with cognitive function. For example, reduced volumes of sub-cortical structures are reliably associated with neurodevelopmental (53F^[14]^), psychiatric (54F^[15]^), and neurodegenerative (55F^[16]^) disorders. The cerebellum is also now considered to be key for several cognitive functions, although it has been historically known for its role in motor functions (56F^[17]^). Other subcortical structures are thought to have specialised roles in cognition, such as the thalamus in memory (57F^[18]^), and the amygdala in risk-based decision making (58F^[19]^). Larger ventricular volumes are also indicative of poorer health, e.g., high blood pressure (59F^[20]^) and diabetes (60F^[21]^), and have been found to be negatively associated with performance on cognitive tests (61F^[22],^62F^[23],^63F^[24])^.

#### g ~ global and subcortical brain structures

In addition to calculating vertex-wise associations (presented in the main paper), we looked at *g-*associations for volume-based global and subcortical measures, as provided in the FreeSurfer aseg outputs. We previously provided the equivalent associations for the 68 Desikan-Killiany regions (^[25]^). Volumes are plotted by cohort and age for global and subcortical structures in Supplementary Figure S19. First, we calculated these for each cohort, and there was strong between-cohort consistency in the relative *g-*association magnitudes across structures (all Pearson’s *r* > 0.724, all *p* < 2.2x10^16^, see Supplementary Table S14). The meta-analysed estimates (see Figure ii and Supplementary Table S15) show that, generally, higher *g* is associated with larger grey matter volumes, and smaller ventricular volumes. The strongest associations were for total grey matter volume and total cortex volume (β = 0.191 and β = 0.183, respectively). Other relatively large or multi-structure measures also have high positive associations: subcortical grey matter (β = 0.156), the brainstem (β = 0.148), and right and left cerebellum grey matter (β = 0.146 and β = 0.133, respectively). Smaller structures with relatively strong positive β values include the bilateral hippocampi and ventral DC (β range 0.117 to 0.148), and the anterior and posterior parts of the corpus callosum (β = 0.127 and 0.131 respectively). Negative associations are comparably weaker and show that higher *g* is associated with smaller inferior lateral ventricles (left β = -0.028 and right β = -0.057), and fewer WM hypointensities (β = -0.063). The similarity in magnitude between subcortical and cortical associations underscores the importance of subcortical structures in studies of *g.*

There was an age moderation effect for *g-*CSF associations (β = -0.011, *FDR Q* = .031), for which there were positive associations for UKB (β = 0.029, *SE =* 0.005, *p* < 2.2x10^16^) and GenScot (β = 0.072, *SE =* 0.027, *p =* .009), and a negative association in LBC1936 (β = -0.077, *SE =* 0.037, *p =* .040). This suggests that greater CSF volume in earlier life is associated with higher *g* but, in later life, increased CSF is associated with lower *g*. In younger ages, CSF volume is more strongly related to intracranial volume (ICV), but as the TBV-ICV association weakens due to atrophy, CSF becomes an important marker of atrophic differences – which can be seen in this divergence of effect sizes. There was also an age moderation effect for *g-*left pallidum associations (β = 0.011, SE = 0.003, *p* = .001); the positive association was stronger in the older cohort with a narrow age range, LBC1936 (β = 0.202, SE = 0.036, *p* = 6.^6x10-8^) than for GenScot (β = 0.057, SE = 0.028, *p* = .044) and UKB (β = 0.091, SE = 0.005, *p* < 2.^2x10-16^). However, there are noticeable differences in the magnitude of the raw volume estimations for the left pallidum between the three cohorts, which might suggest that this result is simply due to image processing differences. For all other age moderation effects for ­global and subcortical volumes with *g* associations, FDR Q > .05, see Supplementary Table S18.

#### Age and sex associations with global and subcortical volumes

For age associations, only GenScot and UKB cohorts were included as the age range of LBC1936 is narrow (mean age = 72.67 years, *SD =* 0.41 years, age range = 71 to 74 years). The between-cohort correlations between the relative β magnitudes across global and subcortical volume associations with age was *r =* 0.967, *p* < 2.2x10^16^ (see Figure ii). Similarly, for sex associations the between-cohort correlations for sex associations with global and subcortical volumes were all high, showing good relative consistency of between-cohort effects - all r > 0.79, see Supplementary Table S14. Between-cohort age moderation effects were not calculated for these age associations, as there were only two cohorts, but there were no age moderation effects on sex-volume associations (all FDR Q > .05, see Supplementary Table S19).

With increasing age, grey matter structures decrease in volume and both ventricles and WM hypointensities increase in volume. Subcortical structures with the largest negative associations between volume and age include the accumbens areas (β = -0.431 and -0.375, left and right), hippocampi (β = -0.364 and -0.363, left and right) and thalami (β = -0.336 and -0.328, left and right), which all have larger associations than total grey matter (β = -0.319). White matter hypointensity volumes have the strongest positive associations with age of all the presently included measures (β = 0.487, *p* < .001).

All meta-analysed sex associations with global and subcortical volumes were positive – in other words, males tend to have larger volumes for all structures than females. The strongest sex-associations were with larger scale measures – e.g., TBV β = 0.466, cerebral GM β = 0.455, cerebral WM β = 0.462, subcortical GM β = 0.466. The smallest associations are for the portions of the corpus callosum (β range = 0.028 to 0.120). See Figure ii for details.

Importantly for the current main focus of this paper, the relative correlation of global and subcortical volumes by *g* and by age β estimates is r = -0.860, p = 2.86x10^13^, suggesting there is strong agreement between global and subcortical volumes that are most strongly associated with *g* and those that change the most with age. The relative correlation between global and subcortical volumes by *g* and by sex associations is r = 0.305, p = .0496, which does not reach the significance threshold.

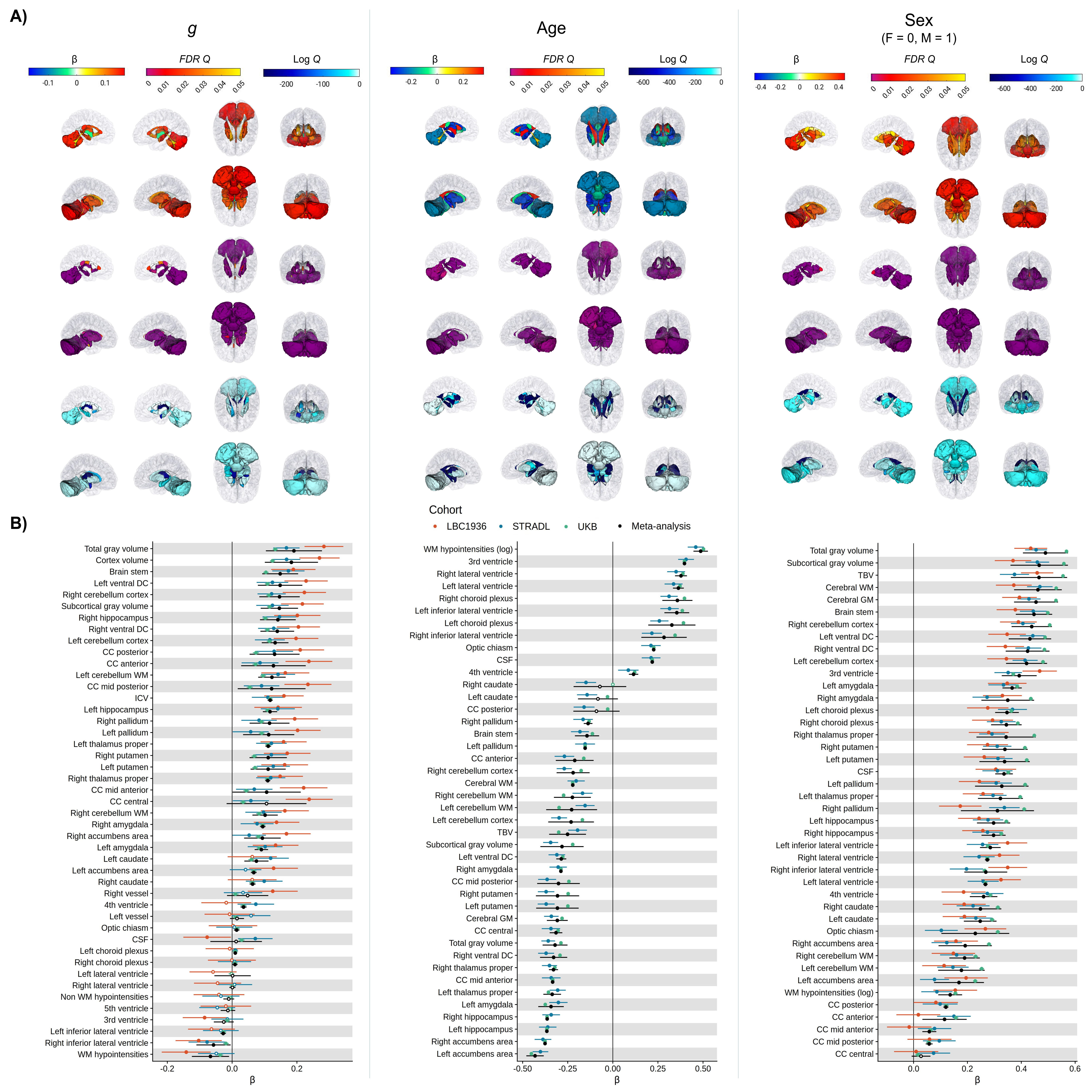

*Figure ii* Associations between *g*, age and sex and global and subcortical structures. A) The meta-analysed subcortical estimates mapped to the brain (top A: β estimates, middle A: log FDR Q values, bottom A: FDR Q values). For β estimates and log Q values, the colour scale limits are set to the maximum of the vertex-wise or present results for each independent variable. For age and sex, some *p* values were estimated at 0, with an R print limit of 1000. For these values, for log Q values, they are set at the limit of the scale which is -704.3. These associations are shown on the ICBM 2009b non-linear asymmetric mni template brain. B) Forest plot showing standardised β estimates for each cohort and meta-analysed estimates.

### Supplementary Analysis 3: Do regional *g*-morphometry associations differ by sex?

Although we controlled for sex in the main analyses, we decided to additionally conduct a supplementary analysis to test whether regional *g-*morphometry associations differ by sex. Across global and subcortical analyses, there were high correlations between the β values derived from males and those from females (all *r* > 0.80, all *p* < .0001, see Figure iii). However, for vertex-wise analyses, whilst there were similarly high correlations for male and female groups for UKB (all *r* > 0.753), correlations tended to be smaller for both LBC1936 and GenScot (see Table i). There are several possible explanations for these findings. It may be that smaller sample sizes lead to less stable association patterns (N_maleUKB_ = 17393, N_femaleUKB_ = 19358, N_maleGenScot_ = 408, N_femaleGenScot_ = 606, N_maleLBC1936_ = 294, N_femaleLBC1936_ = 328). Additionally, the LBC1936 is a narrow-age cohort (mean age = 72.67 years, *SD =* 0.41) and the GenScot imaging sample was selected for depression – sex differences in the way that brain morphometry relates to *g* might be more pronounced in these samples than in the more generally sampled UKB. If meaningful sex differences exist in the general population, we would have expected to see them in the UKB sample.

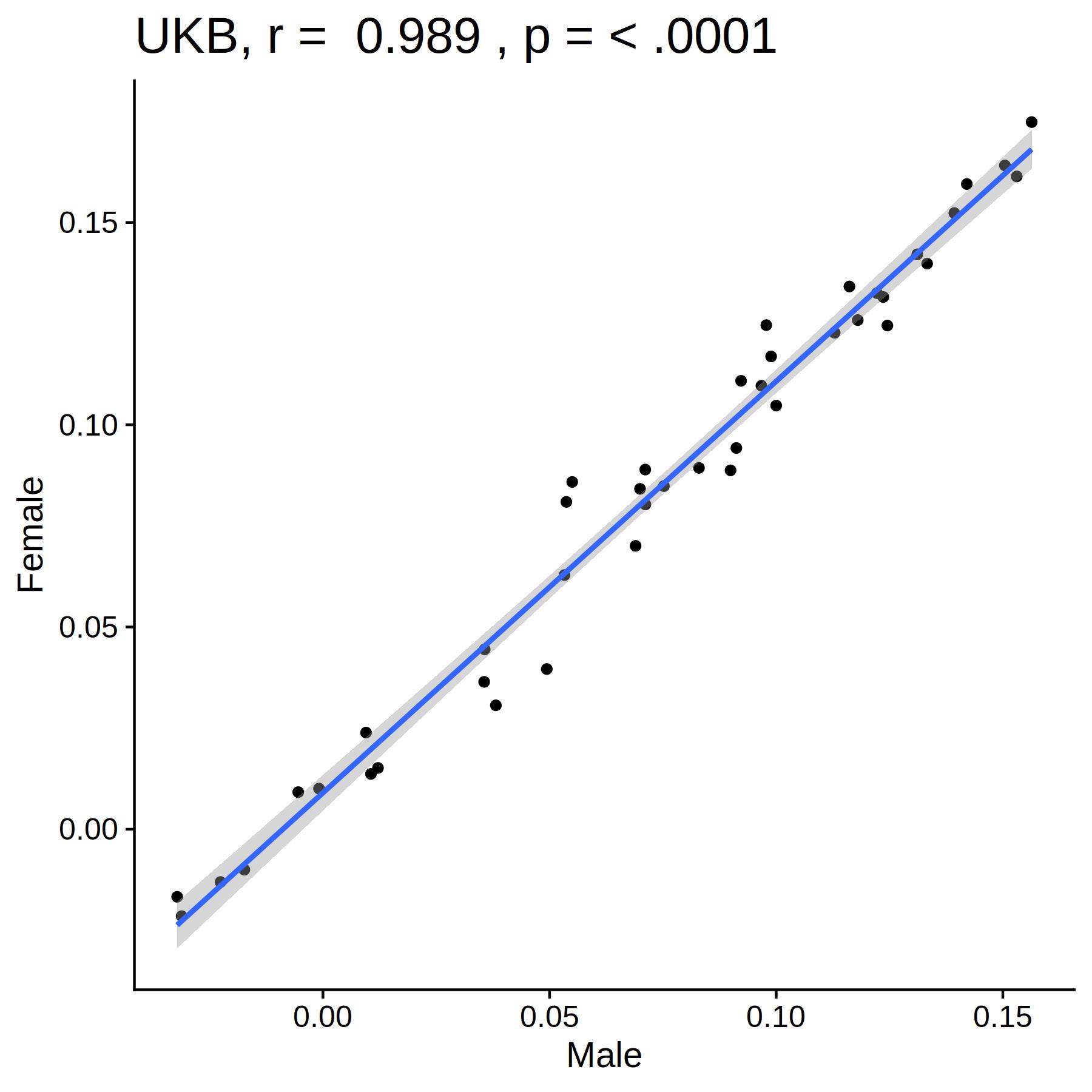

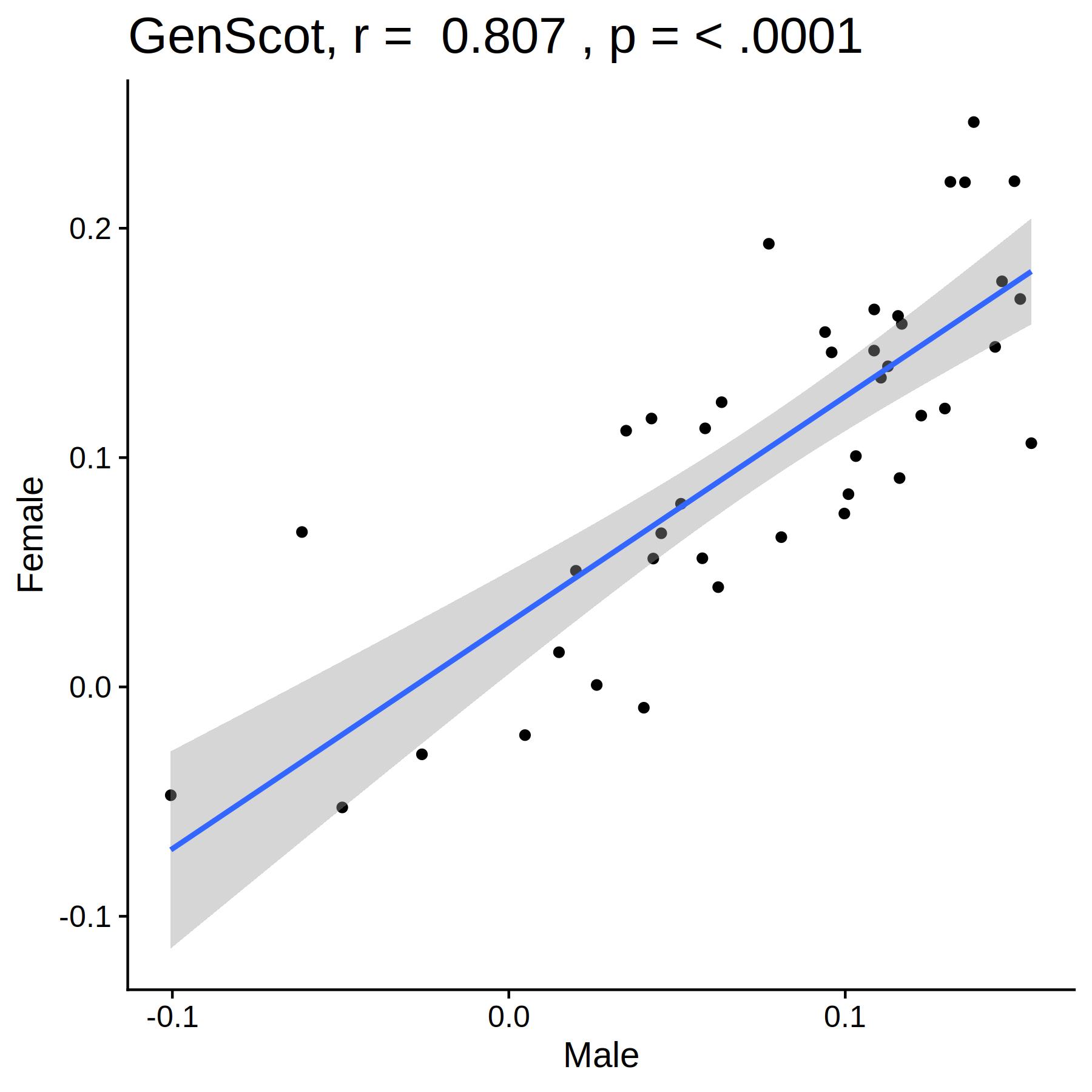

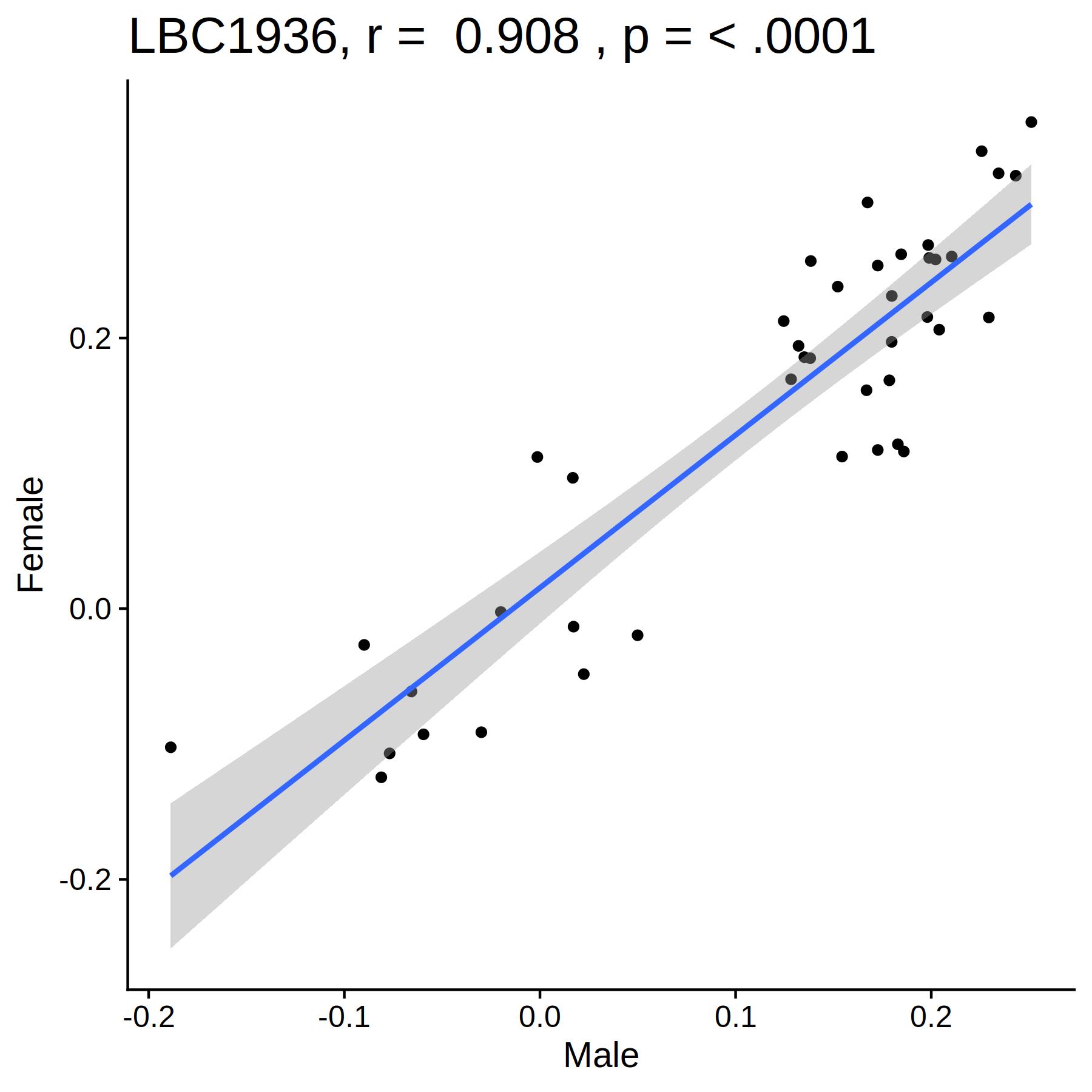

*Figure iii* Scatter plots showing correlations between *g*-volume estimates for global and subcortical structures for males against females.

*Table i* Correlations between male and female *g-*morphometry profiles in each cohort (Pearson’s *r*).

|  | LBC1936 | GenScot | UKB |
| --- | --- | --- | --- |
| Volume | 0.281 | 0.152 | 0.808 |
| Surface area | 0.368 | 0.247 | 0.847 |
| Thickness | 0.307 | 0.349 | 0.753 |
| Curvature | 0.374 | 0.172 | 0.831 |
| Sulcal depth | 0.536 | 0.426 | 0.844 |
